# Plastid fatty acid export (FAX) proteins in *Arabidopsis thaliana* - the role of FAX1 and FAX3 in growth and development

**DOI:** 10.1101/2023.02.09.527856

**Authors:** Wassilina Bugaeva, Anne Könnel, Janick Peter, Julia Mees, Valentin Hankofer, Cordula Schick, Alexander Schmidt, Alexander Banguela-Castillo, Katrin Philippar

## Abstract

In plant cells, fatty acid (FA) synthesis occurs in the plastid stroma and thus requires subsequent FA export for lipid assembly in the endoplasmic reticulum. In this context, the membrane-intrinsic protein FAX1 has been described to mediate FA-export across the plastid inner envelope (IE). In *Arabidopsis,* FAX1 function is crucial for pollen cell wall formation, male fertility, cellular lipid homeostasis and plant biomass. Based on conserved structural features and sequence motifs, we here define the plant FAX-protein family localized in plastids. Besides their membrane-intrinsic domain, the plastid-targeted FAX1-FAX3 contain distinct N-terminal stretches. Among them, the apolipoprotein-like α-helical bundle of FAX2 is the most prominent. Further, we could unequivocally localize FAX2 and FAX3 proteins together with FAX1 to the IE membrane of chloroplasts and develop a topology model for FAX1, FAX2, and FAX3. In yeast, all plastid FAX proteins – i.e. FAX1, FAX2, FAX3, FAX4 – can complement for FA-transport function. For FAX1 we show a new function in pollen tube growth as well as together with FAX3 in seed/embryo development and in rosette leaf growth. Since in comparison to *fax1* single knockout mutants, *fax1/fax3* double knockouts are seedling lethal and not able to develop mature rosette leaves, we conclude that FAX1 and FAX3 function together in vegetative leaf growth.

**Highlight:** We define distinct structural features of plant FAX proteins in plastids and demonstrate that FAX1 and FAX3 have complementary functions in vegetative leaf growth.

## Introduction

Within the plant cell, fatty acids (FAs) are essential building blocks for structural – i.e. membrane building – and energy-storage acyl lipids. Further, FAs are basic units for plant-produced lipophilic biopolymers such as cutin and wax on the epidermal surface or sporopollenin and tryphine covering the outer pollen cell wall (for overview see Li-Beisson et al., 2013). *De novo* synthesis of FAs takes place in the plastid stroma, while breakdown by ß- oxidation is accomplished in peroxisomes (Li-Beisson et al., 2013, Troncoso-Ponce et al., 2015). In the cyclic pathway of FA synthesis, the acyl chain is repeatedly elongated by the FA synthase complex and typically reaches lengths of C16 or C18, called long-chain FAs. In general, for synthesis and metabolic processing, FAs are activated via a thioester linkage to either the acyl carrier protein or coenzyme A (CoA), respectively (Kalinger et al., 2020). After synthesis, long-chain FAs are assembled into acyl lipids either in plastids via the so-called prokaryotic pathway or in the endoplasmic reticulum (ER) by the eukaryotic pathway. In the model plant *Arabidopsis thaliana* about 38% of the FAs remain in the plastid, where they are mainly utilized for assembly of structural lipids that build the chloroplast internal membrane systems (Browse et al., 1986, Somerville and Browse, 1991). The majority (62% of newly synthesized FAs), however, is exported from the plastid stroma to the endoplasmic reticulum (ER). Here, all phospholipids for non-plastid, cellular lipid bilayer membranes, and energy storage triacylglycerol (TAG) oils (Li-Beisson et al., 2013, Bates, 2016, Xu and Shanklin, 2016), as well as precursors for complex extracellular lipophilic compounds such as epidermal wax and cutin (Beisson et al., 2012, Lee and Suh, 2015, Fich et al., 2016) or sporopollenin and tryphine of the outer pollen cell wall (Ariizumi and Toriyama, 2011, Shi et al., 2015, Wan et al., 2020) are produced. However, also a considerable amount of these eukaryotic lipids is relocated into plastids for lipid assembly. In consequence, a eukaryotic-type acyl-glycerol backbone has to be imported into plastids (LaBrant et al., 2018, Lavell and Benning, 2019).

In principle, transport and distribution of lipophilic compounds in the plant cell can be accomplished by membrane transport proteins (Li et al., 2016, Li-Beisson et al., 2017, Lavell and Benning, 2019) and by exchange via membrane contacts between organelles or vesicular traffic (Hurlock et al., 2014, Block and Jouhet, 2015, Michaud and Jouhet, 2019). Well-studied primary active transport systems for lipids are ABC transporter such as the TGD complex in the inner chloroplast envelope (IE) membrane for import of eukaryotic lipids into plastids (for overview see LaBrant et al. [2018], Lavell and Benning, [2019]). At-ABCA9 and Cr-ABCA2 mediate uptake of acyl-CoA into the ER lumen in *Arabidopsis* seeds (Kim et al., 2013) and in the green microalgae *Chlamydomonas reinhardtii* (Jang et al., 2020), respectively. At-ABCD1 in peroxisome membranes binds acyl-CoA as substrate but due to its thioesterase activity most likely transports free FAs into peroxisomes for ß-oxidation (Baker et al., 2015, Carrier et al., 2019). Further, an entire set of ABC transporter proteins of subfamily G is responsible for transport of precursors of wax, cutin, sporopollenin, tryphine compounds across the plasma membrane (for review see Do et al. [2017], Dhara and Raichaudhuri [2021], Grafe and Schmitt [2021]).

In 2015 we were able to describe the crucial role of the FAX1 (fatty acid export1) protein in export of FAs across the IE membrane of plastids in *Arabidopsis* (Li et al., 2015). FAX1 contains a so-called Tmemb_14 domain with four membrane-intrinsic α-helices and is able to mediate FA transport in heterologous yeast cells. The directional transport of FAs in yeast (Black and DiRusso, 2003, Claus et al., 2019) and most likely also across the plastid envelope (Li et al., 2016) is accomplished by a process called vectorial acylation or esterification, which involves membrane intrinsic transport proteins and the ATP-dependent coupling of FAs to CoA by acyl-CoA synthetases. In yeast these are the proteins Fat1 (membrane-intrinsic) and Faa1/Faa4 (acyl-CoA synthetases), in plastids it is proposed that FAX1 and FAX2 proteins in the IE can act together with the long chain acyl-CoA synthetase LACS9 at the outer envelope (OE) membrane (Li et al., 2016, Li-Beisson et al., 2017, Tian et al., 2019). *At-FAX1* is strongly expressed in tapetum cells in anthers and thereby is essential for deposition of the lipophilic sporopollenin and tryphine compounds on the outer pollen cell wall (Li et al., 2015, Zhu et al., 2020). In consequence, *fax1* knockout (ko) mutants show male sterility and are impaired in tapetum programmed cell death and transcriptional regulation (Li et al., 2015, Zhu et al., 2020). Furthermore, we could demonstrate by the detailed study of *At-FAX1* ko and overexpressing lines that the function of FAX1 is important for cellular lipid homeostasis -e.g. for ER-produced TAG oils and phospholipids -, cuticular wax composition and plant biomass (Li et al., 2015). In addition, it was shown by colleagues that seed-specific overexpression of At-FAX1 increases seed oil content in *Arabidopsis* and that the FAX1 ortholog in *Brassica napus* as well contributes to seed oil production (Tian et al., 2018, Xiao et al., 2021). Further, FAX1 relatives in green and red microalgae are implicated in TAG lipid metabolism as well (Li et al., 2019, Takemura et al., 2019, Chen et al., 2022, Chen et al., 2023). In this context we could show only very recently that in *Chlamydomonas reinhardtii* FAX1 is localized in the chloroplast envelope and represents a bottleneck for TAG production (Peter et al., 2022). Further, for the first time we could provide unequivocal evidence for ER membrane insertion of Cr-FAX5, which in *Chlamydomonas* functions in the same metabolic pathway as Cr-FAX1 – i.e. TAG synthesis (Peter et al., 2022). Detailed analysis of conserved motifs in the Tmemb14_domain of FAX1 together with the observation that a strong FAX1-overexpressing *Arabidopsis* line contains heavily curved, honey comb-like IE membranes in plastids, lead us to the hypothesis that the amphiphilic third helix of FAX1 Tmemb_14 contributes to IE membrane bending and stability (Könnel et al., 2019). Thus, we propose that FA transport by FAX1 at least in part might be mediated by membrane bending and possible membrane contacts between IE and OE.

Since in *Arabidopsis* FAX1 belongs to a family of seven proteins, in the recent years several publications addressed the function of plastid-predicted FAX1-relatives by mutant analysis. For At-FAX2, a function in seed TAG oil production could be shown by CRISPR/Cas9 generated knockout and seed-specific overexpression (Tian et al., 2019). A more recent study claims that together with FAX2 also FAX4 has a role in seed oil accumulation in *Arabidopsis* (Li et al., 2020). However, if At-FAX4 really has a prominent role in seed oil delivery so far still remains questionable, since the study by Li and co-workers (2020) used a *fax2/fax4* double mutant line that was generated from single T-DNA insertion lines in the 5’ UTR of both genes. The identical *fax2* single mutant line by Tian et al. (2019), however, was shown to contain considerable amounts of *FAX2* transcripts and displayed no phenotype. Further, in our hands *fax4* mutants with 5’UTR insertion of T-DNA as well contain residual transcripts (see Results section of this study). Thus, most-likely normal expression of *At-FAX4* can complement the only partial loss of FAX2 function in the 5’UTR single *fax2* mutant of Li et al. (2020), but not the complete CRISPR/Cas9 *fax2* knockout (Tian et al., 2019). If in addition to *At-FAX2* also *At-FAX4* transcripts are reduced in the *fax2/4* double mutant generated by Li and co-workers, a seed phenotype appears (Li et al., 2020). The ortholog of At-FAX3 in poplar trees (*Populus trichocarpa*) was described to be involved in plastid export of the FA-derived lipid molecule OPDA (12-oxo-phytodienoic acid), which is synthesized in plastids and in the cytosol is the precursor for synthesis of the wounding hormone jasmonic acid (Zhao et al., 2021). In all studies on FAX2, FAX3, and FAX4 proteins (Tian et al., 2019, Li et al., 2020, Zhao et al., 2021), however, subcellular localization in plastid envelopes so far was only documented by GFP-targeting in *Arabidopsis* protoplasts, which does not allow to discriminate undoubtedly between the OE and IE membranes.

In this study, we define the plant FAX protein family in plastids by the detailed analysis of conserved domains and motifs in their protein sequences. The Tmemb_14 domain of all FAX proteins contains a mixture of hydrophobic and amphiphilic helices in combination with positively and negatively charged amino acid residue that is conserved among subfamily members. Interestingly, plastid-localized FAX1, FAX2, and FAX3 have specific N-terminal domains that most likely contribute to protein-protein interaction and FA-transport function. Here the most prominent is the apolipoprotein-like helix bundle of FAX2 proteins. By combined GFP-targeting and immunoblot analysis, we for the first time could unequivocally localize FAX2 and FAX3 proteins together with FAX1 to the IE membrane of chloroplasts. Further, proteolysis of IE membrane vesicles revealed a membrane topology model for FAX1, FAX2, and FAX3. In yeast, all plastid localized At-FAX1-4 proteins are able to complement FA-transport function of Fat1. For FAX1 we can show an *in planta* function in pollen tube growth as well as together with FAX3 in seed/embryo development and in rosette leaf growth. Since in comparison to *fax1* single ko, *fax1/fax3* double ko mutants are seedling lethal and not able to develop mature rosette leaves, we conclude that the FAX3 protein can complement the loss of FAX1 function in vegetative leaf growth.

## Materials and methods

### Plant material and growth conditions

Experiments on *Arabidopsis thaliana* wild-type plants were performed with ecotype Columbia-0 (Col-0; Lehle seeds, Round Rock, USA) and Wassilewskija (WS-4; INRA, Versailles Arabidopsis Stock Center, France). The T-DNA insertion lines *fax2-1* (FLAG_426D07) and *fax3-1* (FLAG_454B11) were purchased from INRA, Versailles Arabidopsis Stock Center; *fax4-1* (SAIL788B10), *fax4-2* (SAIL1176C06) and *fax4-3* (GK-930E07) from Nottingham Arabidopsis Stock Center, UK. *Fax1-1* (SAIL_66_B09) and *fax1-2* (GABI_559E01) are as previously published (Li et al., 2015). All mutant lines were PCR-genotyped and the presence of *FAX1*, *FAX2*, *FAX3, FAX4* transcripts was controlled by RT-PCR (Supplementary Fig. 8). The *fax1/fax3* double mutant (*fax1/3* dm) was generated by manually pollinating stigmata of *fax1-2* plants with mature pollen from homozygous *fax3-1* mutants. In the subsequent F1 generation, progeny with a heterozygous genotype for both mutant alleles was selected and was further screened for segregation of T-DNA insertions in F2 and F3 filial generations.

To synchronize germination, all seeds were kept at 4°C for 2–3 days prior to growth. Plants were either grown on sterile plates (0.8 % Agar-Agar [Roth], 1 % sucrose, 0.5x MS salts, 0.05 % MES at pH 5.8) or on soil (Potground H, Geeste, Germany). Before sowing on agar plates, seeds were surface-sterilized with 5 % hypochloride and ethanol. Plant growth occurred in growth chambers with a 16 h light (21°C; photon flux density of 100 µmol m^-2^ s^-1^) and 8 h dark (16°C) cycle.

### Segregation and seed development analyses of fax1/3 double mutants

Because in F2, F3 progeny no double homozygous *fax1/3* mutant lines (*fax1/3* dm_ho/ho) could be identified, we chose two independent lines with *fax1-2* heterozygous and *fax3-1* homozygous genotype (*fax1/3* dm_he/ho, lines #4y10 and #5r10) for further analysis in the F4 generation. For segregation analysis, we repeatedly germinated defined amounts of seeds on agar (20) and soil (50) media, and determined the percentage of seedlings developed after three weeks of growth. Subsequently, the phenotype of plantlets was documented by photography and seedlings were completely processed for PCR-genotyping.

Seed development was monitored in mature siliques from the F3 generation of line #4y10 with a *fax1-2* heterozygous and *fax3-1* homozygous background (*fax1/3* dm_he/ho). The corresponding segregated wild-type allele for *fax1-2*, i.e. *fax1/3* dm_wt/ho, was selected as control. Mature green siliques were harvested from six and five different plants of *fax1/3* dm_he/ho and *fax1/3* dm_wt/ho, respectively. Subsequently, 5-10 siliques from each individual plant were destained in 100 % EtOH and phenotyped with a binocular loupe. We therefore determined the amount of fully developed seeds as well as that of empty white seed coats and gaps in each silique.

### Transcript quantification

To monitor transcript content, total RNA was extracted from rosettes of 4- or 5-week-old *Arabidopsis* wildtype and the respective mutant lines, using the NucleoSpin RNA Plant and Fungi kit (Macherey-Nagel). In general, rosette tissue of three individual plants was pooled for each sample to decrease biological variation. For quantitative real-time RT-PCR (qRT-PCR), RNA was reverse-transcribed into cDNA and quantification of transcripts was performed as described previously (Duy et al., 2007) using a LightCycler system (Roche). All signals were normalized to that of actin 2 (At3g18780), actin 8 (At1g49240) cDNA fragments (simultaneous detection by oligonucleotide primer pair). For gene-specific oligonucleotide primers, see Supplementary Table S2.

### In vitro pollen germination assays

*In vitro* pollen germination assays were performed according to (Boavida and McCormick, 2007). After harvesting, freshly opened flowers, not older than one day (see Boavida and McCormick, 2007), were gently dabbed onto 1 cm² big cellophane sheets placed on solid medium (10 % sucrose; 0.01 % H_3_BO_3_; 1 mM MgSO4; 5 mM CaCl_2_; 5 mM KCl; 1.5 % low melting point agar, pH 7.5) to release pollen grains. Pollen tube germination occurred at maximum humidity in closed containers at 22°C for 16 h in darkness. After incubation, germinated pollen tubes as well as the total amount of pollen grains were determined by light microscopy. In total, pollen grains harvested from 6 flowers of different individual plants from each genotype were analyzed on three consecutive days, - i.e. overall pollen from 18 different flowers for each genotype.

### Isolation of Pisum sativum FAX2 and FAX3 cDNA

For determination of FAX2 and FAX3 coding sequences from *Pisum sativum* (garden pea), RT-PCR was performed using pea seedling cDNA as template and oligonucleotide primers designed according to a pea EST contig sequence (Franssen et al., 2011). The corresponding mRNA molecule for Ps-FAX2 was 1,056 bp long with 138 bp 5’UTR and a 918 bp coding region. For Ps-FAX3 a coding region of 678 bp was amplified. For GenBank, GenPept accessions and protein features see Supplementary Table S1 and Fig. S7, for oligonucleotide primers Supplementary Table S2.

### Immunoblot analysis and proteolysis

Pea chloroplasts were isolated and sub-fractioned into OE-, IE-membrane vesicles, stroma and thylakoids as described (Waegemann et al., 1992). Proteolysis of IE vesicles (IEVs) by thermolysin (0.1 µg per 1 µg of IE proteins) occurred in buffer (330 mM sorbitol; 50 mM HEPES-KOH pH 7.6; 5 mM CaCl_2_) on ice for 0-15 min. The reaction was stopped by adding EDTA to a final concentration of 5 mM and subsequently samples were subjected to immunoblot analysis.

For immunoblot analysis, appropriate amounts of proteins were separated by SDS-PAGE and transferred to PVDF membranes. Primary antisera were used in 1:500 to 1:1000 dilutions in TTBS buffer (100 mM Tris-HCl pH 7.5; 150 mM NaCl; 0.2 % Tween-20; 0.1 % BSA). Non-specific signals were blocked by 3 % skim milk powder and 0.1 % BSA. Secondary anti-rabbit IgG alkaline phosphatase antibodies (Sigma-Aldrich) were diluted 1:6,000. Blots were stained for alkaline phosphatase reaction with 66 µg/ml nitroblue tetrazolium (NBT) and 33 µg/ml bromochloroindolyl phosphate (BCIP) in AP buffer (100 mM Tris-HCl pH 9.5; 100 mM NaCl; 5 mM MgCl_2_).

Ps-FAX2 and Ps-FAX3 antisera were raised in rabbit (Pineda Antibody Service, Berlin, Germany) against N-terminal peptide sequences of both proteins (see Supplementary Fig. 7). Antisera for Ps-FAX1 and marker proteins were produced as described previously (Küchler et al., 2002, Duy et al., 2007, Philippar et al., 2007, Li et al., 2015). Α-FBPase against *Arabidopsis* fructose-1,6-bisphosphatase was a gift from B. Bölter (LMU Munich).

### In vivo GFP, YFP targeting

C-terminal fusions of GFP and YFP to the preproteins of At-FAX1, At-FAX2, and At-FAX3 were constructed by subcloning PCR-amplified cDNA into pENTR/D/TOPO and further into pK7FWG2 (GFP) or pB7YWG2 (YFP) plasmid vectors (Karimi et al., 2002) using the Gateway cloning system (Invitrogen). Subsequently, *Agrobacterium tumefaciens* AGL1 cells were transformed with the respective plasmids.

For transient transfection of tobacco (*Nicotiana benthamiana*) leaves, exponentially growing liquid *Agrobacterium* cultures of the respective FAX constructs and of a helper plasmid with the p19 protein of tomato bushy stunt virus (Voinnet et al., 2003) were harvested and further incubated for 2 hours at 28°C, 75 rpm in induction medium (10 mM MES, pH 6.0; 10 mM MgCl_2_; 0.2 mM acetosyringone). Next, cells were harvested and resuspended with an OD_600_ of 0.5-0.7 in 5 % sucrose, 0.2 mM acetosyringone. Suspensions of p19 and the respective FAX constructs were mixed 1:1 and were infiltrated into mature tobacco leaves. After two to five days of incubation of intact, infiltrated tobacco plants, leaf mesophyll cell walls were enzymatically digested and protoplasts were isolated for fluorescence microscopy. Therefore, tobacco leaves were cut into thin 5-10 mm pieces and incubated for 90 min at 90 rpm, 25°C in the dark with 1 % (w/v) cellulase R10 and 0.3 % (w/v) macerozyme R10 in F-PIN solution (1.12 % KNO_3_; 0.44 % CaCl_2_; 0.37 % MgSO_4_; 0.17 % KH_2_PO_4_; 3.9 % MES; 20 mM NH_4_-Succinate; 0.1 % BSA; adjusted to 550 mOsm with sucrose). Afterwards, the suspension was filtrated through a 100 µm nylon net into a 15 ml glass tube, overlaid with 2 ml F-PCN (1.12 % KNO_3_; 0.44 % CaCl_2_; 0.37 % MgSO_4_; 0.17 % KH_2_PO_4_; 3.9 % MES; adjusted to 550 mOsm with glucose), and centrifuged for 10 min at 70 g. Intact protoplasts, which accumulated between the enzyme F-PIN solution and the F-PCN overlay, were carefully removed and washed with W5 buffer (18.38 % CaCl_2_; 0.37 % KCl; 0.39 % MES, pH 5.7; adjusted with NaCl to 550-580 mOsm) for 10 min at 50 g. Finally, the protoplasts pellet was gently resuspended in 500 μl W5 buffer and used for fluorescence analysis.

Confocal images were recorded with a confocal laser scanning microscope (Leica, TCS SP5). Fluorescence signals in protoplasts were examined by a 63×1.3 glycerine-immersion objective using an argon laser for excitation (395/475 nm for GFP, 514 nm for YFP). The emitted light of GFP and YFP was detected at 509 nm and 527 nm, respectively. Chlorophyll autofluorescence was exited at 465/665 nm and monitored at 673-726 nm. Images were processed with Leica LA SAF Lite (Leica).

### Drop dilution growth assays in yeast

For growth assays in yeast, the coding sequence of the mature At-FAX2, At-FAX3, and At-FAX4 proteins were subcloned into the yeast expression plasmid pDR195 (XhoI/BamHI). Therefore, we fused the cDNA of the respective predicted mature proteins between an XhoI restriction site, followed by a start codon “ATG” and the native stop codon, followed by a BamHI restriction site with PCR amplification. At-FAX1 and At-PIC1 in pDR195 were produced previously (Duy et al., 2007, Li et al., 2015). The yeast mutant strains Δ *fat1* (LS2020-YB332) and Δ *faa1/faa4* (LS1849-YB525) are specified in (Zou et al., 2003). Both strains were transformed with the respective constructs (see above) or the empty vector control pDR195 as described (Li et al., 2015). Liquid cultures of yeast cells were grown to exponential phase in synthetic defined medium (SD-ura), containing 0.1% (w/v) glucose, 0.7% (w/v) yeast nitrogen base without amino acids, and necessary auxotrophic amino acids without uracil. For drop-dilution tests, 2 μl of the cultures were spotted in different dilutions onto SD-ura plates (2% agar), supplemented with 3.6 mM α-linolenic acid (0.1%, w/v in ethanol), and 1% tergitol (to increase α-linolenic acid solubility). For control plates, an equal amount of the solvent ethanol was added instead of α-linolenic acid. Growth of yeast cells was documented between 2 to 6 days at 30°C.

### Digital analysis of proteins and gene expression

*In silico* structural analysis and predictions for FAX proteins were performed by Phyre^2^ (Kelley et al., 2015), consensus motifs were analyzed by MEME suite (Bailey et al., 2015). Lengths of mature proteins were estimated according to stromal peptidase cleavage site predictions by ChloroP (Emanuelsson et al., 1999) and TargetP 2.0 (Almagro Armenteros et al., 2019). Α-helices of the Tmemb_14 domain were positioned according to the Aramemnon consensus prediction AramTmCon (Schwacke et al., 2003) of *Arabidopsis* proteins. Helical wheel analysis was conducted by heliquest (Gautier et al., 2008) and NetWheels (Mól et al., 2018) software. All heatmaps for gene expression were generated in R (R-Core-Team, 2019) using the heatmap.2 function from the gplots package (Warnes et al., 2020). Euclidean distance and average linkage were used for heatmap data clustering.

## Results

### The plant FAX-protein family in plastids

In *Arabidopsis*, seven proteins belong to the FAX family (Supplementary Fig. S1, Table S1; Li et al. [2015]), which can be divided into 6 subfamilies (FAX1, FAX2, FAX3, FAX4, FAX5/6, FAX7; Fig.1), and groups into the TMEM14 class of proteins (InterPro entry IPR005349) found in nearly all eukaryotic organisms and some bacteria (see PANTHER “transmembrane protein 14” subfamily, PTHR12668). While FAX1 unequivocally has been shown to insert into the chloroplast IE membrane (Li et al., 2015), predictions for intracellular localization (Table S1) and recent publications based on GFP-targeting of At-FAX2, At-FAX4 (Tian et al., 2019, Li et al., 2020) as well point to chloroplast envelopes for FAX2, FAX3 and FAX4 proteins. At-FAX5, At-FAX6 and At-FAX7 in contrast are expected to insert into membranes of the secretory pathway (Table S1). Here the first evidence for ER membrane insertion was provided only recently for FAX5 in the green microalga *Chlamydomonas reinhardtii* (Peter et al., 2022). Common to all FAX/TMEM14 proteins are four conserved α-helices (Fig. 1), which build the so-called Tmemb_14 domain (Pfam entry PF03647). In addition, mature, plastid-localized FAX proteins have an extended and variable N-terminal region (see below). FAX5, FAX6 and FAX7 instead essentially consist of the Tmemb_14 domain and thereby resemble their TMEM14 protein relatives in vertebrates and yeast (InterPro entry IPR005349; (Li et al., 2015). At-FAX5 and At-FAX6 are very similar in sequence (81% identical, Supplementary Fig. S1B) and not in every plant species are represented by two gene copies (Könnel et al., 2019). Thus, they most likely represent paralogs with redundant functions and were grouped into one FAX5/6 subfamily (Fig. 1).

**Fig. 1.**
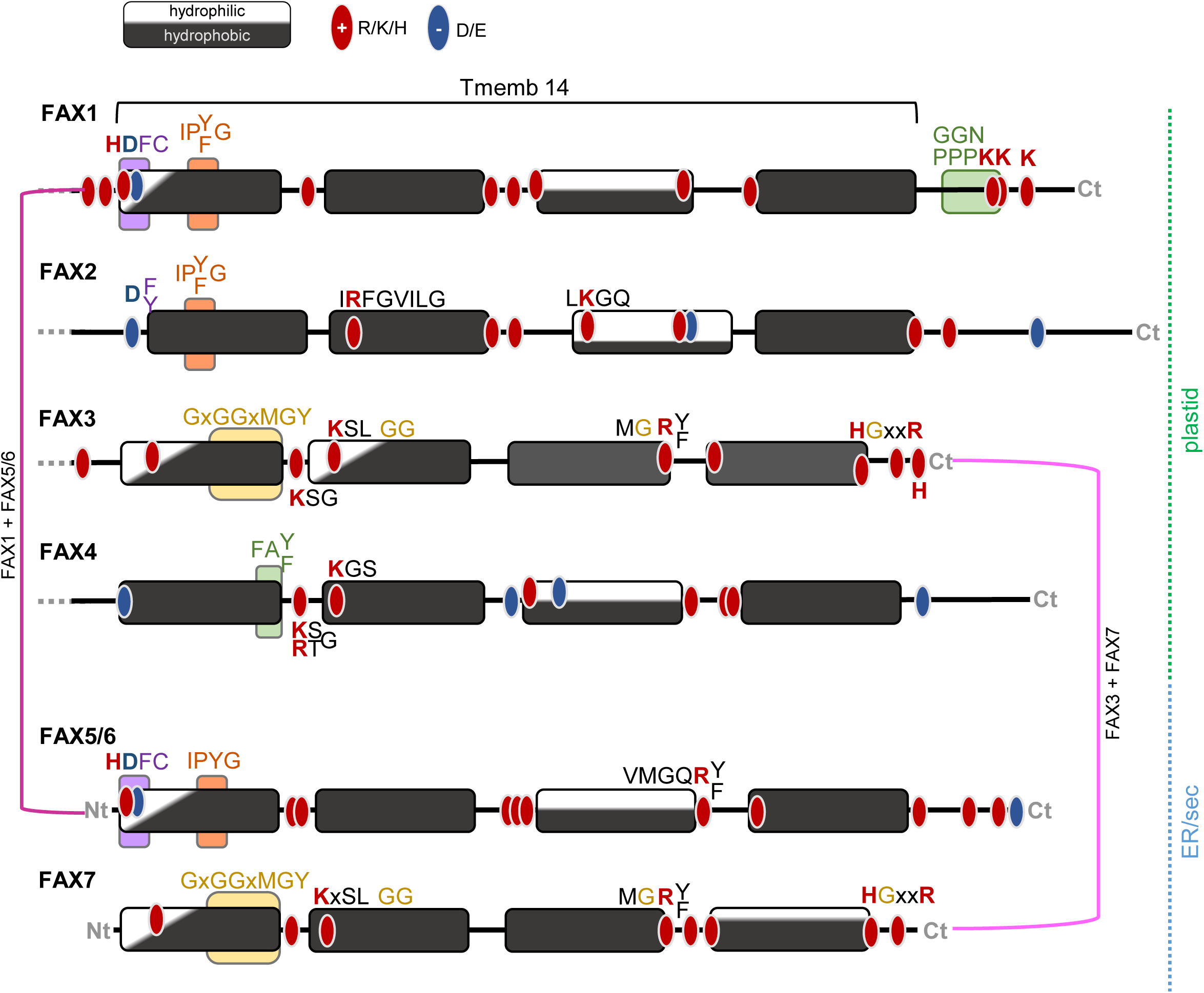
Tmemb14 domains of plant FAX proteins. Plant FAX proteins can be grouped into six subfamilies (FAX1-FAX4, FAX5/6, FAX7) and contain a Tmemb_14 domain (Pfam entry PF03647) at their C-terminal end. The Tmemb_14 domain consists of four α-helices (rectangles) with hydrophilic (white) and/or hydrophobic (black) faces. Within each FAX subfamily, the amphiphilic or hydrophobic character of these α-helices as well as the distribution of positively (R, K, H; red dots) and negatively (D, E; blue dots) charged amino acid residues are highly conserved among different plant species (compare Supplementary Figs. S2-S5, S7, Könnel et al. [2019]). Further, several conserved sequence motifs are indicated by colored boxes and consensus sequences. FAX5 and FAX6 with 81% identity in their amino acid sequences show an identical distribution of conserved amino acids and amphiphilic helices and thus are grouped into one subfamily. As indicated by purple brackets, conserved sequence motifs of FAX1 + FAX5/6 as well as FAX3 + FAX7 are highly similar. Thus, FAX1 and FAX5/6 as well as FAX3 and FAX7 most likely represent two sets of basic FAX proteins in plastids (FAX1, FAX3) and ER/secretory pathway membranes (FAX5/6, FAX7). Please note that the mature plastid proteins FAX1, FAX2, FAX3 and FAX4 contain an extended and variable N-terminal region (see Fig. 2A) as indicated by grey dots, whereas ER/secretory pathway predicted FAX5/6 and FAX7 essentially consist of the Tmemb14 domain. The lengths of polypeptide chains (black lines) are drawn to scale, amino acids are depicted in single letter code.

To identify conserved domains and sequence motifs characteristic for plastid-intrinsic FAX proteins, we searched for relatives of FAX1-FAX4 within the plant kingdom. Please note that in all original publications, i.e. in (Li et al., 2015, Tian et al., 2019, Li et al., 2020) and the Aramemnon plant membrane protein database (Schwacke et al., 2003), FAX2 relatives are similar to the protein with the *Arabidopsis* At2g38550 (At-FAX2), and FAX3 proteins correspond to At3g43520 (At-FAX3). Therefore, we keep this annotation in the following and define the plant FAX protein family as depicted in Fig. 1, Supplementary Fig. S1 and Table S1. FAX1 and FAX3 can be found in angiosperm dicots and monocots, in moss (*Physcomitrium patens*) as well as in the green microalga *Chlamydomonas reinhardtii* (Supplementary Figs. S2, S3; Peter et al. [2022]). In contrast, and well in line with the reported seed-specific function (Tian et al., 2019, Li et al., 2020) plastid expected FAX2 proteins appear to be specific only for seed plants (Supplementary Fig. S4). Relatives to At-FAX4 could be detected in dicots, monocots, *Selaginella moellendorfii* (spikemoss) and *Physcomitrium*, but not in *Chlamydomonas* (Supplementary Fig. S5; Peter et al. [2022]).

Within their Tmemb_14 domain, all FAX proteins contain at least one α-helix with an amphiphilic and at least two α-helices with a hydrophobic character (Fig. 1, Supplementary Fig. S6; Li et al. [2015], Könnel et al. [2019]). In general, it can be assumed that the Tmemb_14 domain of FAX/TMEM14 proteins inserts into lipid bilayer membranes (Klammt et al., 2012, Li et al., 2015). The amphiphilic α-helices as well as the asymmetrically distributed and well conserved, charged amino acid residues in this region thereby might contribute to membrane shaping and/or stabilization (Könnel et al., 2019). The sequence of the Tmemb_14 domain of At-FAX1 is most similar to At-FAX2 and At-FAX5/At-FAX6, while At-FAX3 Tmemb_14 resembles that of At-FAX7 (Supplementary Fig. S1B). Interestingly, sequence motifs and distribution of charged amino acids within the Tmemb_14 domain, in particular in the first α-helices of FAX1 and FAX5/6 as well as of FAX3 and FAX7 are almost identical. Thus, FAX1 together with FAX5/6 (see also Könnel et al. [2019]) as well as FAX3 and FAX7 seem to form two subsets with slightly different features, each containing plastid and ER/secretory pathway-predicted FAX proteins (Fig. 1). This observation is also supported by the finding that these two plastid-ER/secretory pathway sets – i.e. FAX1 + FAX5/6 and FAX3 + FAX7 – are conserved in *Chlamydomonas* (Supplementary Figs. S2, S3, S7; Peter et al. [2022]). In the green microalga we further could show that Cr-FAX1 in the chloroplast envelope and Cr-FAX5 in the ER membrane indeed function in the same metabolic pathway – i.e. in production of the neutral storage lipid TAG (Peter et al., 2022). Further, all FAX1 proteins in plants are characterized by a prominent stretch of two glycines, one asparagine, followed by two to three prolines and two to three lysine residues (GGNPPKK-motif) after helix four (Fig. 1, Supplementary Fig. S2A).

Mature plastid-predicted FAX1-FAX4 contain a highly variable N-terminal region (Fig. 2A, Supplementary Figs. S2-S5). Whereas for dicot and monocot FAX1 proteins we found one additional, non-hydrophobic α-helix at the N-terminus (Supplementary Fig. S2), FAX2 polypeptides with four predicted extra α-helices have the longest N-terminal region (Fig. 2A; Supplementary Fig. S4; Li et al. [2015]). Remarkably, structural modelling of the FAX2 N-terminus revealed that three of these helices (h1, h2, h3) form a so-called helical up-and-down bundle with strong similarity to animal apolipoproteins/apo-lipophorins (Fig. 2B). The mammalian apolipoproteins apoE and apoA-I as well as insect apolipophorin III are described to contain four to five α-helices with amphiphilic character that form helical bundles, which function in solubilization of lipids and/or lipid membranes during biogenesis of high density lipoprotein (HDL) particles (Narayanaswami et al., 2010, Huebbe and Rimbach, 2017). Similar to apolipoprotein/apolipophorin helices (Phillips, 2013) also h1, h2, h3 of FAX2 proteins display a hydrophobic face that separates from patterned charged amino acid residues on the hydrophilic side of the helix (Fig. 2C, D). Although FAX2-like proteins appear to be present in seed plants only, in *Physcomitrium patens* and *Selaginella moellendorffii*, however, we found polypeptides that contain the FAX2-like apolipoprotein N-terminus but have a Tmemb_14 domain and the C-terminal GGNPPKK-motif specific for the FAX1 subfamily (Supplementary Fig. S8). For FAX3 proteins no additional helices are predicted in the N-terminal region, however, the presence of a poly-glycine stretch (polyG) with distinct conserved motifs is quite evident (Fig. 2E, Supplementary Fig. S3A, C). The N-terminus of mature predicted FAX4 proteins is rather short (20 aa in At-FAX4) with no conserved features (Fig. 2A, Supplementary Fig. S5).

**Fig. 2.**
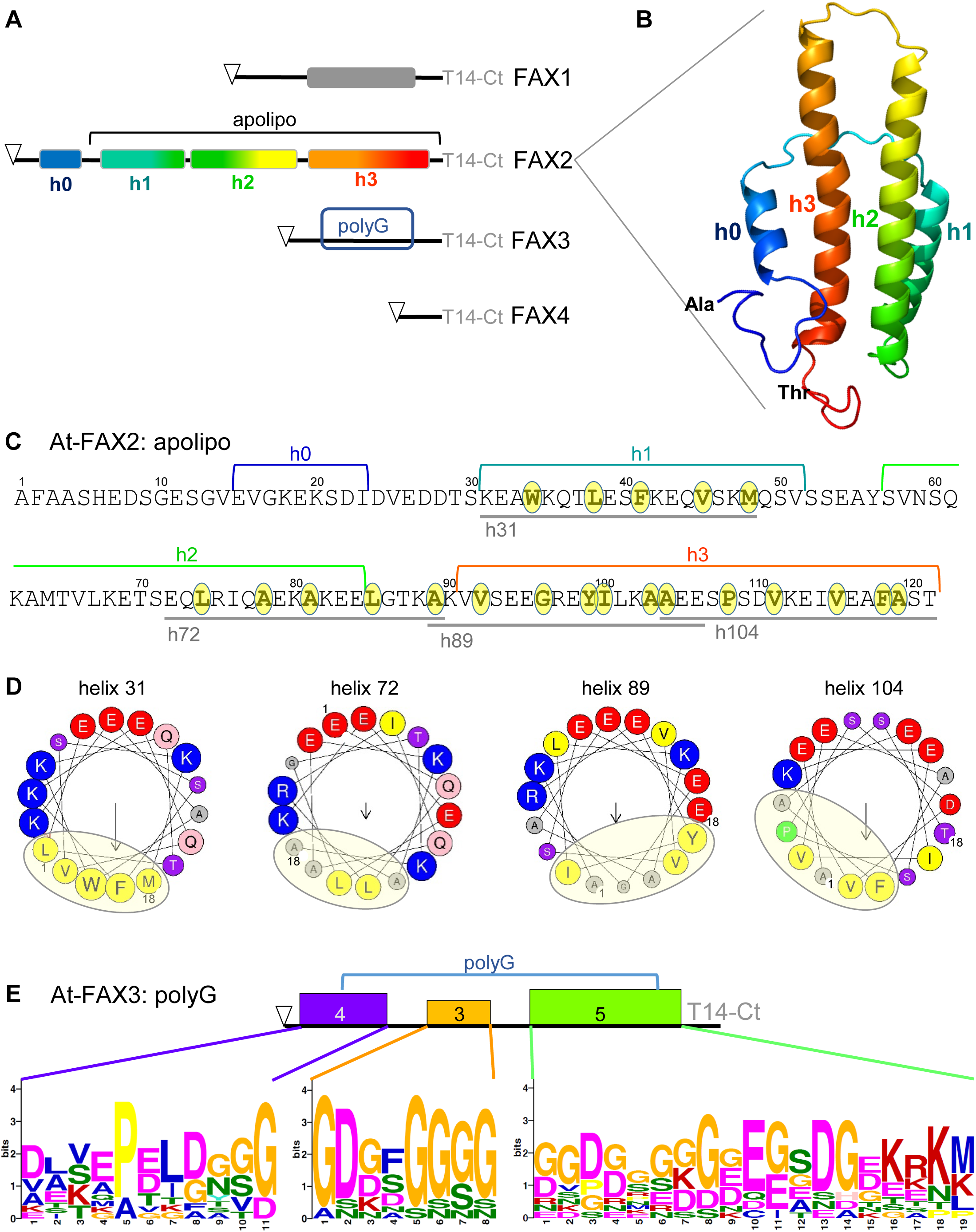
N-terminal domains of plastid-predicted FAX1, FAX2, FAX3, and FAX4 proteins. (A) The variable N-terminal regions of plastid-predicted mature FAX proteins are highly conserved among subfamilies (compare Supplementary Figs. S2-S5, S8). N-terminal of the Tmemb_14 domain (T14-Ct, see Fig. 1), dicot and monocot FAX1 proteins are predicted to contain a non-hydrophobic α-helix (grey rectangle, Phyre^2^ prediction [Kelley et al., 2015]). Mature FAX2 proteins have the longest N-terminal region, which according to *in silico* structural analysis is characterized by four additional α-helices (h0-h3, colored rectangles). Among these, h1-h3 built the so-called apolipoprotein domain (see [B]-[D]). Specific for all FAX3 proteins is an N-terminal poly-glycine stretch (polyG, see [E]), while the short amino terminus of FAX4 is rather unobtrusive. Lengths of polypeptide chains (black lines) are drawn to scale and predicted N-terminal cleavage sites for plastid stromal peptidases are indicated by triangles. Please note that for At-FAX1, Ps-FAX1, Ps-FAX2 and Ps-FAX3 length of predicted chloroplast targeting peptides could be confirmed by peptide sequencing (see Supplementary Fig. S9; Peter et al. [2022]). (B) The secondary structure of the At-FAX2 N-terminus (h1-h3 modelled at 49.5% confidence) was predicted by Phyre^2^ (Kelley et al., 2015) in comparison to the structure of a four-helical up-and-down bundle of a human apolipoprotein (template d1gs9a: 22k domain of hm apolipoprotein e4, PDB entry 1GS9). The expected α-helices h0-h3 are depicted in rainbow color code. N- and C-terminal amino acids of the 121 amino acid (aa) long At-FAX2 N-terminus (alanine and threonine, respectively) are shown. (C) Amino acid sequence of the At-FAX2 N-terminus used for structural prediction in (B). The predicted helices h0-h3 are indicated. Helical wheel analysis (see [D]) reveals a hydrophobic face for four 18 aa-long α-helical turns (h31, h72, h89, h104, grey lines). Amino acids forming these hydrophobic faces are highlighted in yellow. (D) Helical wheel analysis of the 121 aa N-terminus of At-FAX2 (heliquest [Gautier et al., 2008]) shows hydrophobic faces in helices h31, h72, h89, and h104 (18 aa each, see [C]). These hydrophobic, non-charged regions (light yellow circles) as well as a distinct pattern of positively (blue) and negatively (red) charged amino acids along the helix axis are characteristic for amphiphilic α-helices of apolipoproteins (Phillips, 2013). (E) In the polyG stretch at the N-terminus of plant FAX3 proteins, three consensus motifs could be identified by the MEME suite software (Bailey et al., 2015). The positions of the motifs 4 (blue), 3 (orange), and 5 (green; see Supplementary Fig. 3A and C) are depicted on the N-terminus of mature At-FAX3 protein. Please note that the conserved glycine repeats start after the conserved proline in motif 4 and are accompanied by conserved negatively charged amino acid residues, namely aspartate and glutamate. Amino acids in (C)-(E) are given in single letter code.

### FAX1, FAX2 and FAX3 integrate into the inner envelope of chloroplasts

Besides At-FAX1, proteomic analysis of chloroplast membranes also identified At-FAX2 and At-FAX3 as potential chloroplast IE proteins (Table S1; Ferro et al. [2010], Bouchnak et al. [2019]). Further, At-FAX2 together with At-FAX4 was described to play a role in FA-export across the plastid envelope of developing seeds (Tian et al., 2019, Li et al., 2020). To clarify the subcellular localization of FAX2 and FAX3 proteins, we generated antisera against peptide stretches of *Pisum sativum* (pea) FAX2 and FAX3 orthologs, for which we isolated the corresponding cDNA from pea seedling RNA beforehand (Table S1, Supplementary Fig. S9). Subsequent immunoblot analysis on sub-fractionated pea chloroplast membranes showed a localization in the IE membrane of chloroplasts for Ps-FAX2 and Ps-FAX3 (Fig. 3A). The antiserum against the Ps-FAX2 peptide detected two distinct bands of around 31-32 kDa in close proximity, which might be due to alternative processing of the chloroplast targeting peptide and/or structural variances in the N-terminal apolipoprotein domain. α-Ps-FAX3 stained a signal of about 21 kDa. With both antisera, however, signals appeared exclusively in the IE membrane and not in outer envelopes (OE), thylakoids or stroma. As for Ps-FAX1 (band at 25 kDa, predicted mature protein at 20.6 kDa, see Fig. 3A; Li et al. [2015]), the apparent molecular weight of Ps-FAX2 and FAX3 proteins on SDS gels thereby was about 3-4 kDa bigger than expected for the predicted mature polypeptides (Ps-FAX2 at 27 kDa, Ps-FAX3 at 16.6 kDa; compare Table S1). Proper prediction of processing sites for stromal peptidase in Ps-FAX1, Ps-FAX2 and Ps-FAX3 preproteins was supported by trypsin proteolysis, followed by peptide sequencing of the corresponding bands from SDS-separated pea IE proteins (Supplementary Fig. S9). Please note that due to the absence of FAX4-like expressed sequences tags in pea (database in Franssen et al. [2011]), immunoblot analysis on purified pea chloroplast membranes could not be performed for FAX4.

**Fig. 3.**
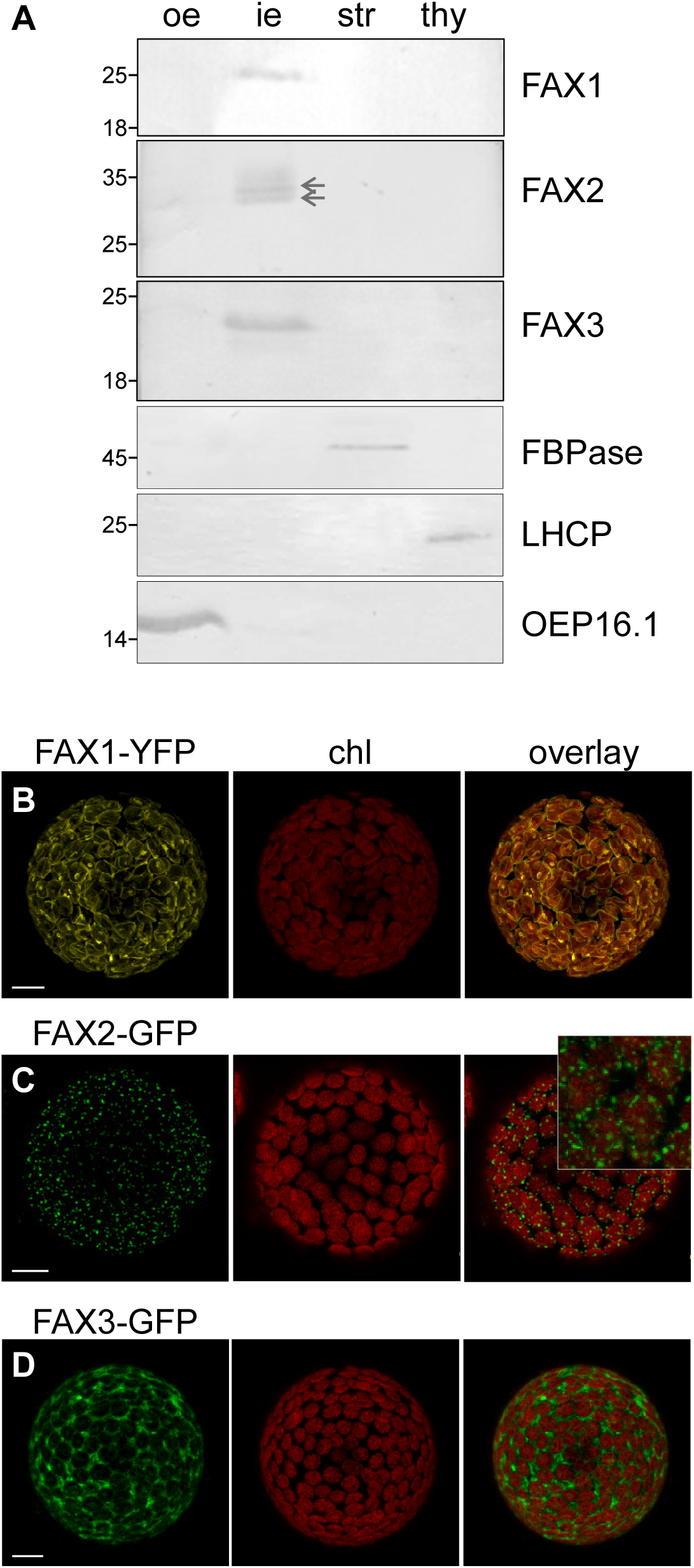
FAX1, FAX2, and FAX3 localize to the chloroplast IE membrane. (A) Immunoblot analysis of FAX1, FAX2 and FAX3 in fractionated pea chloroplasts. Pea chloroplasts were isolated and fractionated into outer (oe) and inner (ie) envelope membranes, stroma (str) as well as thylakoid membranes (thy). 10 µg of proteins were separated by SDS-PAGE and subjected to immunoblot analysis using antisera directed against Ps-FAX1, Ps-FAX2 and Ps-FAX3. The chloroplast outer envelope protein OEP16.1, the stromal enzyme fructose-1,6-bisphosphatase (FBPase) and thylakoid light harvesting complex proteins (LHCP) served as controls. For OEP16.1, FBPase and LHCP detection, 5 and 0.5 µg of proteins were used in each lane, respectively. Please note that Ps-FAX2 signals appear as double band at 31-32 kDa (arrows). Numbers indicate molecular mass of proteins in kDa. (B-D) *In vivo* GFP-targeting of At-FAX1, At-FAX2, and At-FAX3. *Nicotiana benthamiana* leaf protoplasts were transiently transformed with constructs for At-FAX1-YFP (B), At-FAX2-GFP (C), and At-FAX3-GFP (D). Images show GFP/YFP-signals (left), chlorophyll fluorescence (middle), as well as an overlay of both (right). The inset in (C) magnifies an overlay of At-FAX2-GFP and chlorophyll signals. Scale bars = 10 μm.

*In vivo* GFP-targeting of *Arabidopsis* FAX1, FAX2 and FAX3 proteins in tobacco (*Nicotiana benthamiana*) protoplasts confirmed a chloroplast envelope localization (Fig. 3B-D). FAX1-YFP (Fig. 3B) and FAX3-GFP (Fig. 3D) constructs showed ring-like signals around chloroplasts well-known for envelope-intrinsic membrane proteins (Breuers et al., 2012, Li et al., 2015, Voith von Voithenberg et al., 2019). The fluorescence of FAX2-GFP (Fig. 3C) in contrast, appeared to concentrate in small, irregular spots at the chloroplast periphery, a pattern, which has been previously shown for At-FAX2 in *Arabidopsis* by Tian and co-workers (Tian et al., 2019). Please note that in our hands *in vivo* targeting of At-FAX-GFP failed in tobacco, most likely because FAX4-like genes could not be in the genome of *Nicotiana benthamiana*. At-FAX4-GFP in *Arabidopsis* protoplasts as shown by Li et al. (2020), however, gives ring-like signals around chloroplasts, pointing to a chloroplast envelope localization of At-FAX4 as well.

To further determine topology and orientation of FAX1, FAX2, FAX3 proteins within the IE membrane, a protease treatment of IE vesicles (IEV) from pea chloroplasts, followed by immunoblot analysis was performed (Fig. 4A). In general, it has been shown that purified envelope membranes from isolated pea chloroplasts form vesicles with predominantly right-side-out orientation (Keegstra and Yousif, 1986). Thus, the endoprotease thermolysin, which cleaves polypeptides N-terminal of bulky, hydrophobic amino acid residues, can access only those peptide stretches of membrane-intrinsic proteins that are exposed to the outside of the IEVs. *In vivo*, the latter region corresponds to the intermembrane space (IMS) between OE and IE envelope membranes. For immunoblot analysis of pea FAX1, FAX2 and FAX3 proteins after proteolysis of IEVs, we used antisera directed against peptide regions at the N-terminus of the respective protein (see Supplementary Fig. S9). Thus, only protein fragments containing the intact, non-degraded N-terminus could be detected. The protein TIC62, which is attached to the IE at the stroma site (Küchler et al., 2002, Stengel et al., 2008), was used as a control and confirmed vesicle orientation (Fig. 4A). Thermolysin treatment of Ps-FAX1 and Ps-FAX3 proteins after 15 min led to stable, IE-membrane protected polypeptide fragments each of about 15 kDa, which both were detected by the respective antisera (Fig. 4A). This indicates that the N-termini of both proteins point to the inner face of the vesicles, which means *in vivo* to the stroma site (Fig. 4B). Signals of Ps-FAX2 protein fragments instead disappeared after 1 min of thermolysin treatment (Fig. 4A). Hence, the N-terminus of FAX2 rather points to the outer face of the IEVs and thus *in vivo* to the IMS (Fig. 4B). Based on the organization of the four α-helices in their Tmemb_14 domains (compare Fig. 1), we interpret the proteolytic patterns and the membrane topology (Fig. 4B) as follows: (i) Intact, mature Ps-FAX1 proteins appear at 25 kDa on SDS-PAGE (see Figs. 3A, 4A; Li et al. [2015]). Thermolysin treatment of Ps-FAX1 produced one major fragment of about 15 kDa (Fig. 4A), which can be assigned to the N-terminus after thermolysin cleavage between the first and second α-helical transmembrane domain. The predicted size of the cleaved and in the immunoblot non-detectable C-terminus after helix 1 is 10.8 kDa. Two weaker signals of about 20 and 18 kDa most likely correspond to intermediate fragments and/or proteolysis of the short soluble C-terminal end of Ps-FAX1 (1.9 kDa) pointing to the IMS (Fig. 4B). (ii) For Ps-FAX2 (mature protein at 31-32 kDa), early during proteolysis a band at 18 kDa, which quickly disappeared after 30 sec, was detected. Considering the peptide-epitope site at the N-terminus and the 13 kDa-sized C-terminal Tmemb_14 domain of Ps-FAX2, this polypeptide fragment might be assigned to the N-terminal apolipoprotein helix-bundle of FAX2 (compare Figs. 2, 4B). Thus, the apolipoprotein-like N-terminus of FAX2 most likely is located in the IMS, and the corresponding unprotected polypeptide was rapidly degraded by thermolysin. (iii) The fragmentation pattern of Ps-FAX3 confirms four transmembrane α-helices in the IE (compare Fig. 1). Right after 30 sec of proteolysis, two protein fragments appeared at ∼18 and ∼15 kDa (Fig. 4A). During prolonged thermolysin incubation, the upper band disappeared whereas the lower 15 kDa band was stable and increased in strength. This specific proteolysis pattern can be explained by two thermolysin cleavage sites inside Ps-FAX3 (apparent molecular weight at 21 kDa), which are exposed to the outside of the IEVs and are located in the short peptide loops between α-helices 1-2 and 3-4, respectively (see Fig. 4B). The 18 kDa fragment most likely originates when the C-terminus with α-helix 4 (3.6 kDa) of Ps-FAX3 is separated from the rest of the protein. The stable 15 kDa fragment therefore apparently represents the N-terminus with the first transmembrane α-helix of Ps-FAX3, thus the second thermolysin cleavage must occur between α-helix 1 and 2.

**Fig. 4.**
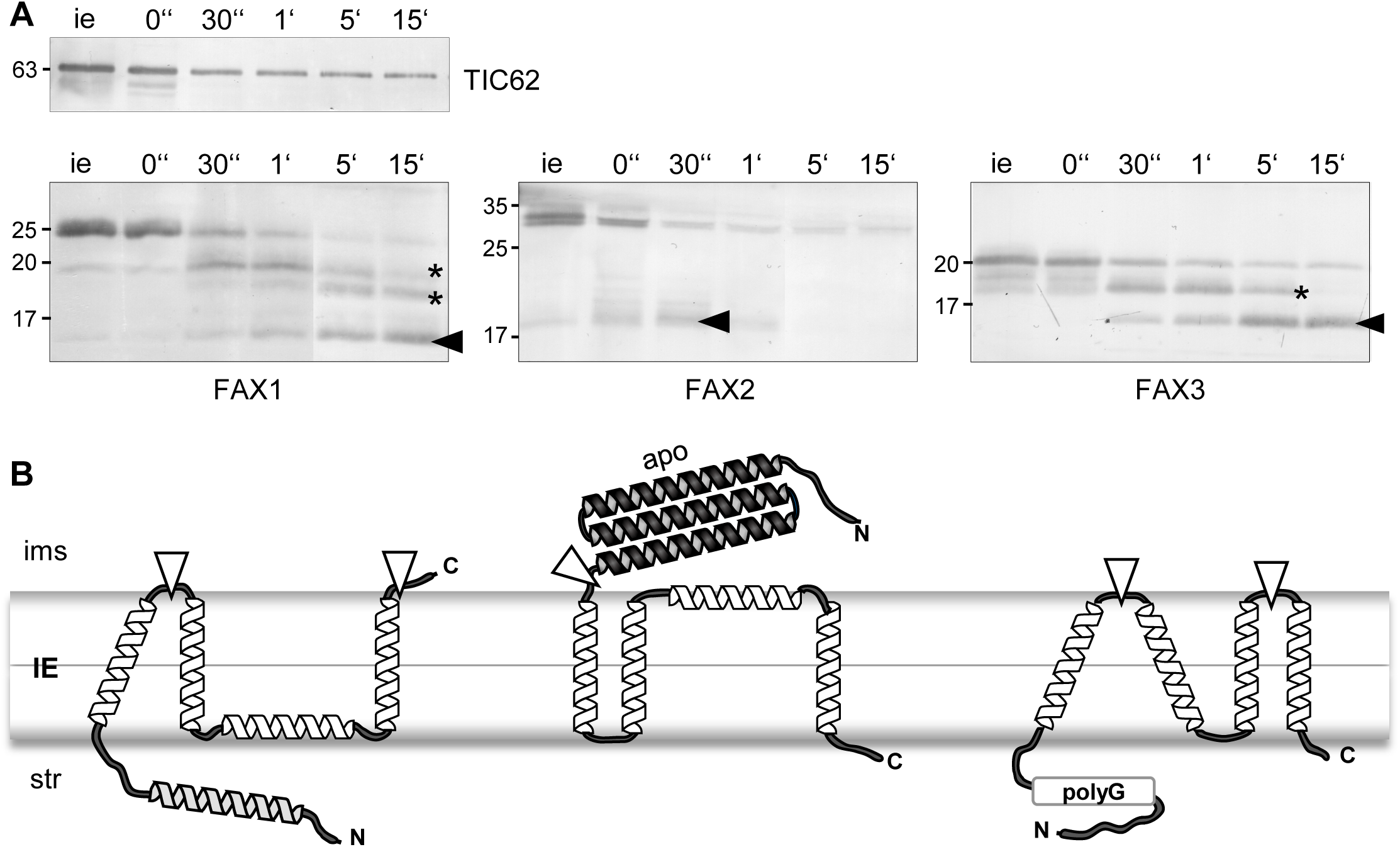
Membrane topology of FAX1, FAX2, and FAX3 proteins. (A) Proteolysis of *Ps*-FAX proteins. Inner envelope membrane vesicles (IEVs) from pea chloroplasts were protease treated with thermolysin (0.1 µg per 1 µg of IE protein) for 0, 30 sec, 1 min, 5 min, and 15 min. Subsequently, 15 µg of protein loaded equally in each lane were separated by SDS-PAGE and subjected to immunoblot analysis with antisera directed against peptide stretches within the N-termini of Ps-FAX1 (left), Ps-FAX2 (middle) or Ps-FAX3 (right; compare Supplementary Fig. S7). Antiserum against the stroma-localized and IE-attached TIC62 was used as control (upper panel). The positions of apparent major thermolysin degradation products of FAX1 (∼15 kDa), FAX2 (∼18 kDa) and of FAX3 (∼15 kDa) are indicated by arrowheads, intermediate fragments are highlighted by asterisks. IEVs without thermolysin addition (ie) were used as control for signals of intact proteins (compare Fig. 3A). Numbers indicate molecular mass of proteins in kDa. (B) Topology models of FAX1, FAX2, and FAX3 proteins in the IE membrane of chloroplasts. Membrane-embedded hydrophobic and amphiphilic α-helices (white) as well as the N-terminal α-helix of FAX1 (grey), the apolipoprotein helix-bundle of FAX2 (apo, dark grey), and the poly-glycine stretch (polyG) of FAX3 are depicted according to results in Fig.1 and Fig.2. White triangles indicate exposed intermembrane-space cleavage sites for thermolysin proteolysis in (A). ims, intermembrane space; str, stroma

In summary, our data demonstrate that FAX1, FAX2 and FAX3 insert into the chloroplast IE membrane. Interpretation of proteolytic patterns of FAX1, FAX2 and FAX3 proteins in pea IEVs and the respective features of the four membrane-embedded α-helices within the Tmemb_14 domain (compare Fig. 1) lead to the topology models as depicted in Fig. 4B. The most likely non-membrane associated N-terminus of FAX1 is located in the plastid stroma, while the short, positively charged C-terminus points to the IMS. In contrast, the N-terminal apolipoprotein-domain of FAX2 presumably can be found in the IMS and the C-terminus in the stroma. For FAX3, both, the N-terminus with the polyG stretch and the C-terminus are predicted in the stroma. In this model the first helix of the Tmemb_14 domain of FAX1 is vertically tilted in the membrane and the third amphiphilic helices of FAX1 and FAX2 immerge into the IE lipid bilayer from the stroma and IMS side, respectively. The first two helices of the Tmemb_14 region of FAX3, however, would capture a hairpin-like topology in the IE membrane.

### FAX gene expression in rosette leaves of plastid FAX knockout mutants

Previous studies revealed that upon loss of FAX1 in *fax1-1* and *fax1-2* ko lines, transcripts of *FAX2* and *FAX3* in *Arabidopsis* flower tissue were increased (Li et al., 2015), thereby most likely compensating for impaired plastid FA-export and leading to the rather mild overall phenotype of *fax1* ko. To follow regulation of *FAX* gene expression in response to *FAX1* deletion in the vegetative state of plant development, we analyzed the transcript content of all seven *Arabidopsis FAX* genes in mature rosette leaves of the *fax1-2* ko mutant (Fig. 5A). Here, transcripts of all chloroplast-localized proteins – i.e. of FAX2, FAX3, FAX4 –, and of the secretory-pathway predicted FAX5 increased significantly (between 2.6- to 3.5-fold) when compared to wildtype. Further, the mRNA level of *FAX6* was 1.8-fold higher in *fax1* than in the corresponding wild-type rosette leaves. Based on available public transcriptomic data, gene expression of *FAX1* and *FAX3* is highest and most closely correlated throughout entire leaf development and is accompanied by that of *FAX5* until fully expanded rosette leaf stages (Fig. 5B, C). In contrast, transcript levels of *FAX2*, *FAX6*, and *FAX7* are lower during leaf development but their expression profiles cluster together. Minimal *FAX4* gene expression, however, clearly outgroups. In summary, in mature rosette leaf tissue of *fax1* ko, gene expression of all chloroplast-intrinsic FAX proteins appears to be increased to compensate for the loss of FAX1 FA-export function. Most likely, FAX3, which is co-expressed during wild-type leaf development, is the most important substitute for FAX1, whereas FAX4 might be dispensable during vegetative growth of *Arabidopsis*. In addition, up-regulation of *FAX5* and *FAX6* expression in *fax1* ko plants points to a potentially enhanced flow of FAs into the ER and/or secretory pathway organelles.

**Fig. 5.**
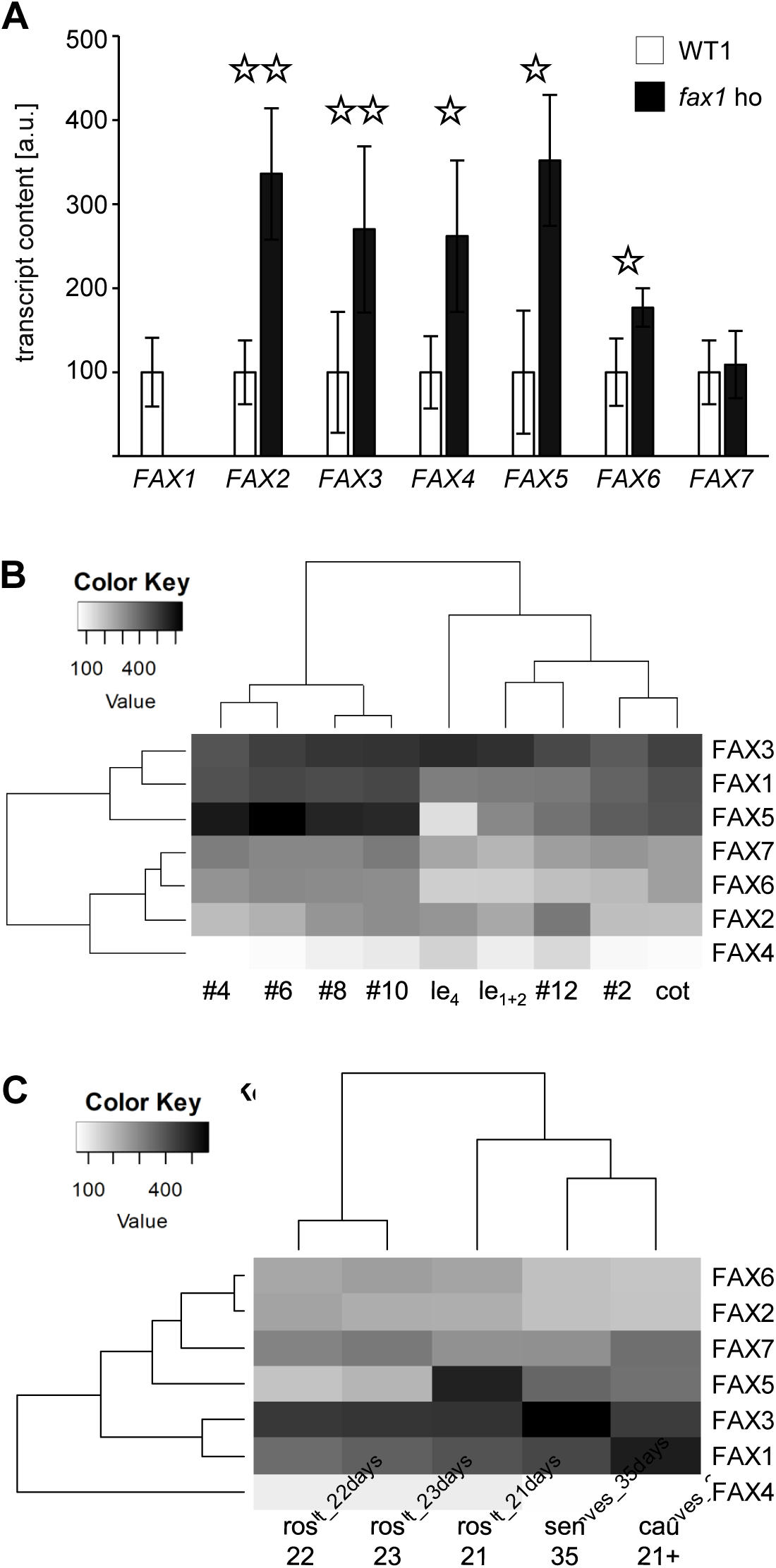
Gene expression of *FAX1* and *FAX3* correlates during *Arabidopsis* leaf development. (A) Expression of *Arabidopsis FAX* genes in rosette leaves of a FAX1 loss-of-function mutant. The transcript content of *At-FAX1*, *At-FAX2*, *At-FAX3*, *At-FAX4*, *At-FAX5*, *At-FAX6*, and *At-FAX7* was analyzed by qRT-PCR in rosette leaves of 5-week-old homozygous *fax1-2* knockouts (*fax1* ho, black bars), and the corresponding segregated wild-type allele (WT1, white bars). The mRNA of each *Arabidopsis FAX* gene was quantified relative to that of actin 2/8 and normalized to the respective amount in WT1, which was set to 100 (arbitrary units). Data points represent mean values from three independent biological replicates (n = 3 ± SD). Please note that for each biological replicate, rosette tissue from three individual plants was pooled prior to RNA extraction. Asterisks indicate significantly different values when compared to WT1 (*: p < 0.05, **: p < 0.01, Student’s t-test). (B), (C) Correlation of gene expression of *Arabidopsis FAX* genes in rosette leaves during vegetative development (B) and after transition to flowering (C). Heatmaps depict gene expression intensities, displayed in grayscale as shown in the color key. The darker the shades of grey, the higher the transcript level in the corresponding developmental phase. *FAX* genes are in rows, leaf development stages are organized in columns. Gene expression data used to create heatmaps are based on DNA microarray analyses obtained from AtGenExpress developmental series (Schmid et al., 2005). Please note that in these analyses, all plants were grown in continuous light. (B) cot, cotyledons of 7-day-old seedlings; le_1+2_, leaves no. 1 and 2 of 7-day-old seedlings; le_4_, rosette leaf no. 4 (1 cm long) of 10-day-old plants; #2, #4, #6, #8, #10, #12, different rosette leaves of 17-day-old plants. (C) cau_21+, cauline leaves of at least 21-day-old plants; sen_35, senescing leaves of 35-day-old plants; ros_21, _22, _23, entire rosette after transition to flowering, but before bolting from 21-, 22-, 23-day-old plants, respectively.

To further follow the function of chloroplast IE intrinsic FAX proteins, we characterized T-DNA insertion lines of *At-FAX2*, *At-FAX3,* and *At-FAX4* (Supplementary Fig. S10). By RT-PCR, we could demonstrate that *fax2-1*, *fax3-1,* as well as *fax4-3* represent knockout lines for the respective *FAX* gene (Supplementary Fig. S10A-C). The lines *fax4-1* and *fax4-2* with T-DNA insertions in the 5’UTR, however, do not represent loss-of-function mutants for *FAX4*, since residual transcripts are present. Please note that this finding is in contrast to a recent publication by Li and coworkers, who showed by RT-PCR that *FAX4* mRNA was absent in a T-DNA insertion line in the 5’UTR of *FAX4* (Li et al., 2020). Although FAX2 and FAX4 recently have been described to contribute to plastid FA-export in developing *Arabidopsis* seeds (Tian et al., 2019, Li et al., 2020), neither *fax2-1*, *fax3-1,* nor *fax4-3* single ko lines displayed obvious phenotypes during vegetative growth and seed/silique development under standard conditions. Furthermore, progeny of heterozygous lines segregated in a Mendelian fashion (Supplementary Fig. S10D), indicating normal gametophyte and embryo development. When we probed for gene expression of all other *FAX* genes in the mutant lines mentioned above, we found that in *fax2-*ko rosette leaves, transcripts of *FAX4* - but not of *FAX1* or *FAX3* - significantly increased (about 2-fold; Supplementary Fig. S11A). This is in line with the complementary function of FAX2 and FAX4 in plastid FA-export during seed development recently described (Li et al., 2020). As observed for *fax1* ko (see Fig. 5A), also transcripts for the secretory pathway predicted *FAX5*, *FAX6*, and for *FAX7* increased in *fax2-1* (Supplementary Fig. S11A), again indicating requirement of enhanced flow of FAs to the ER/secretory pathway when FAX function in plastids is impaired. Further, expression profiles of *FAX2*, *FAX6*, and *FAX7* cluster together during *Arabidopsis* leaf development (see Fig. 5B, C). In contrast, the loss of FAX3 and FAX4 in rosette leaves did not lead to a change in transcript levels of other *FAX* genes (Supplementary Fig. S11B, C). Thus, the deficit in function of FAX3 in vegetative growth under standard conditions, in *Arabidopsis* appears to be compensated by the already available set of chloroplast FAX proteins, most likely by FAX1, which is co-expressed at rather high levels throughout leaf development (compare Fig. 5B, C). Due to its low expression in leaves (see, Fig. 5B, C), FAX4 function in vegetative growth most likely is dispensable.

### Plastid FAX proteins can functionally complement for FA-uptake in yeast

In order to test, whether the plastid localized FAX1, FAX2, FAX3, and FAX4 are able to functionally complement for each other, we analyzed their capability to transport FAs in heterologous yeast cells. In yeast, the import of long-chain FAs occurs by a process called vectorial acylation and is mediated by a protein complex consisting of Fat1, a membrane-spanning transport protein, and Faa1 or Faa4, acyl-CoA synthetases for intracellular FA activation (Black and DiRusso, 2003). However, when challenged with high external concentrations of toxic FAs such as polyunsaturated α-linolenic acid (C18:3), yeast cells that contain a functional FA-uptake and activation pathway die (von Berlepsch et al., 2012, Li et al., 2015). As demonstrated for At-FAX1 (Li et al., 2015) and At-FAX2 (Tian et al., 2019), we performed FA-transport complementation assays for the mature At-FAX2, At-FAX3, and At-FAX4 proteins in yeast *fat1* and *faa1/faa4* knockout mutants in the presence of high external α-linolenic acid supply (Fig. 6). Here Δ*fat1* cells, which expressed FAX1, FAX2, FAX3, or FAX4, died with 3.6 mM α-linolenic in the medium (Fig. 6A-D), thereby demonstrating that FA-uptake can be mediated via plastid FAX-proteins when the yeast endogenous Fat1 transporter is missing. In contrast, Δ*fat1* cells transformed with the mature chloroplast iron (Fe) permease At-PIC1 (Fig. 6E) or empty vector controls (Fig. 6A-E) survived α-linolenic treatment. Especially the result for At-PIC1, a chloroplast IE Fe-transport protein with four membrane-intrinsic α-helices, which can mediate Fe-uptake into yeast cells and chloroplasts (Duy et al., 2007, Duy et al., 2011), pinpoints the specificity of FAX proteins for FA-transport. Furthermore, yeast knockouts of the acyl-CoA synthetases for intracellular FA activation (Δ*faa1/faa4*) survived α-linolenic treatment under identical conditions (Fig. 6A-E). The latter finding indicates that transport of free FAs across lipid bilayer membranes via FAX proteins requires concurrent acylation of free FAs to CoA by acyl-CoA synthetases, similar to the so-called vectorial acylation function of the yeast Fat1 and Faa1/Faa4 proteins (see (Black and DiRusso, 2003, Zou et al., 2003, Li et al., 2016) for discussion).

**Fig. 6.**
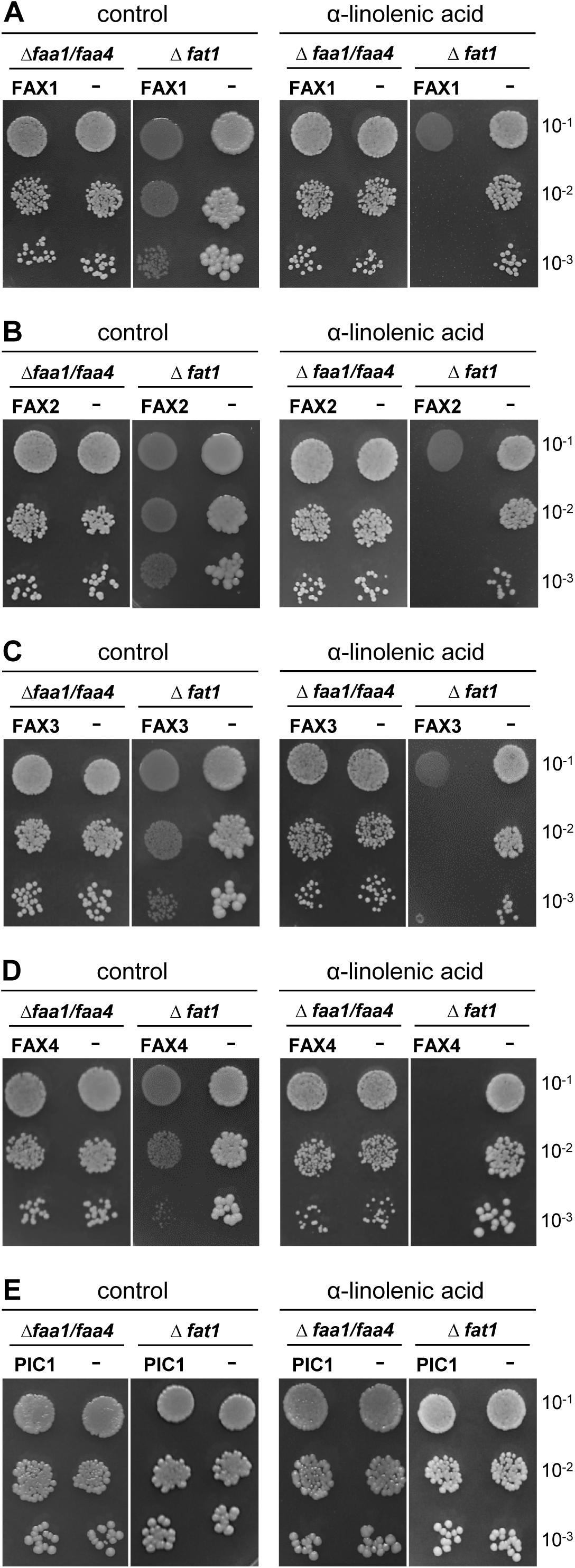
Plastid-localized FAX proteins can mediate fatty acid-transport in yeast. The empty plasmid pDR195 (-) and the cDNA of the mature proteins At-FAX1 (A), At-FAX2 (B), At-FAX3 (C), At-FAX4 (D), and of At-PIC1 (E) in pDR195 were introduced into Δ*faa1/faa4* and Δ *fat1* yeast mutant strains, respectively. Drop dilution assays were performed with 2 µl per droplet of serial dilutions (OD_600_ 10^−1^, 10^−2^, and 10^−3^) of exponentially growing yeast cells on SD-ura plates (0.1% glucose, 1% tergitol). Control plates (left) are shown in comparison to plates supplemented with 3.6 mM of the C_18:3_-fatty acid α-linolenic acid (right).

### The role of FAX1 and FAX3 in growth and development of Arabidopsis

FAX1 and FAX3 appear to represent the basic set of plastid-localized FAX proteins conserved throughout the green lineage ranging from green microalgae like *Chlamydomonas reinhardtii* (compare Peter et al. [2022]) to angiosperm plants (see Fig. 1 and Supplementary Figs. S2, S3). Further, while *At-FAX1* and *At-FAX3* are tightly co-expressed during vegetative leaf growth and development (see Fig. 5), a seed-specific function for At-FAX2 and At-FAX4 has been described (Tian et al., 2019, Li et al., 2020). Thus, among the plastid localized *At-FAX* genes differentially regulated in *fax1* ko – i.e. *FAX2, FAX3, FAX4* (see Fig. 5A) – *FAX3* is most likely to be able complement for FAX1 function. Therefore, we decided to further examine the *in planta* role of FAX1 and FAX3 in more detail, in particular in *Arabidopsis* pollen tube growth as well as rosette leaf development.

### FAX1 is essential for pollen development and tube growth

In our previous study, we already demonstrated that FAX1 is essential for pollen development and outer pollen cell wall formation (Li et al., 2015). These findings were complemented by the study of Zhu and co-workers (Zhu et al., 2020), who showed that *At-FAX1* is strongly expressed in tapetum cells in anthers and its loss-of-function impairs pollen wall formation as well as tapetum programmed cell death and transcriptional cascades. In consequence, the loss of FAX1 leads to male sterility and non-Mendelian segregation of the homozygous *fax1* ko-alleles (about 7 % and 4 % in *fax1-1* and *fax1-2* mutant lines, respectively; Li et al. [2015]). Accordingly, *FAX1* represents the plastid FAX protein with the highest gene expression in pollen development (Fig. 7A; Li et al. [2015], Zhu et al. [2020]), especially during maturation from uninucleate to tricellular microspores. During transition from bi-to tricellular pollen stages, a shift in expression of *FAX* genes occurs, because except for *FAX1*, transcript levels of all other plastid-localized FAX proteins drop down, whereas those of ER/secretory pathway-predicted FAX5 and FAX6 significantly increase (Fig. 7A). Moreover, FAX1 also was the only FAX protein that could be identified in proteomic analysis of mature *Arabidopsis* pollen grains (Grobei et al., 2009).

**Fig. 7.**
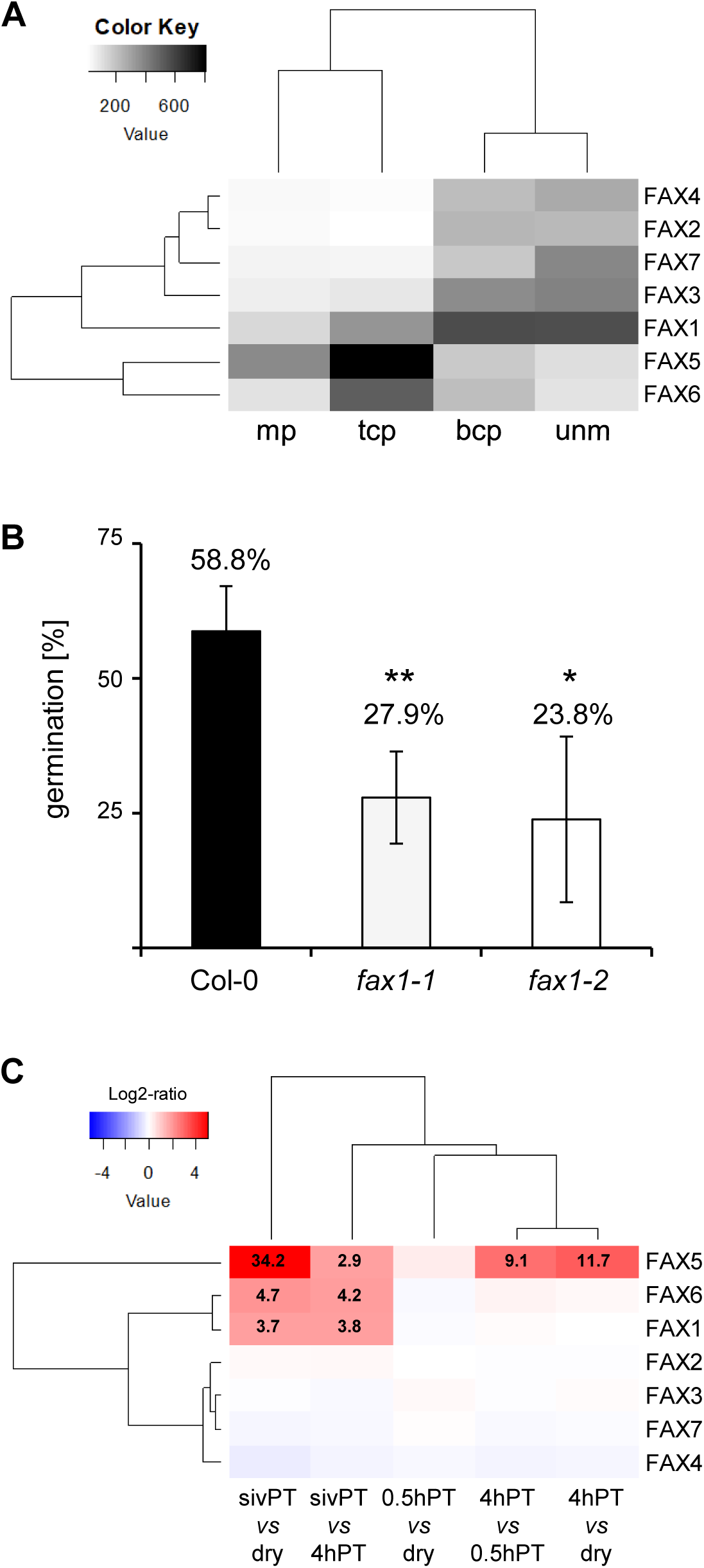
FAX1 is essential for pollen development and pollen tube growth. (A) Correlation of gene expression of *Arabidopsis FAX* genes during pollen development. The heatmap depicts gene expression intensities displayed in grayscale, as shown in the color key. The darker the shades of grey, the higher the transcript level. *FAX* genes are in rows, pollen developmental stages are organised in columns. Gene expression data used to create the heatmap is based on DNA microarray analyses of pollen development (Honys and Twell, 2003) and AtGenExpress developmental series (Schmid et al., 2005). unm, uninucleate microspore; bcp, bicellular pollen; tcp, tricellular pollen; mp, mature pollen grain. (B) *In vitro* pollen tube germination of pollen grains from Col-0 wildtype and heterozygous *fax1-1*, *fax1-2* mutants. Mature pollen grains from freshly opened flowers were germinated in three independent *in vitro* assays. Data (amount of visible pollen tubes in %) represent mean values ± SD from flowers with a pollen germination rate > 10%. This corresponds to n = 16, n = 13 and n = 7 flowers as well as 1861, 1701, and 856 pollen grains analyzed for Col-0, *fax1-1* and *fax1-2*, respectively. Asterisks indicate significantly different values when compared to Col-0 (*: p < 0.005, **: p < 0.00005, Student’s t-test). Please note that two flowers of *fax1-1* and of *fax1-2* lines released pollen with less than 10% and 5% germinated tubes, respectively and thus these data are not included in the depicted graph. Further, pollen grains of 2, 3, and 9 flowers from Col-0, *fax1-1* and *fax1-2*, respectively, failed to produce any tubes. (C) Differential gene expression of *Arabidopsis FAX* genes during pollen tube growth. The heatmap depicts the log2-ratio of the fold change of *Arabidopsis FAX* genes expression between two pollen developmental stages. Red color indicates up-regulation and blue indicates down-regulation as shown in the color key. *FAX* genes are in rows, whereas pollen tube growth conditions are organised in columns. Expression fold-changes of significantly up-regulated *FAX* gene expression (p-value < 0.001) are given in the corresponding cell. Data was gathered from Genevestigator® based on published DNA microarray data of pollen tube growth (Qin et al., 2009). dry, dry mature pollen grains; 0.5hPT, 4hPT, pollen tubes grown for 0.5h and 4h *in vitro*; sivPT, “semi *in vivo”* conditions for pollen tubes growing through stigma/style tissue of native pistils.

In order to examine, whether further male gametophyte defects influence the reduced segregation of *fax1* ko alleles, we tested for the role of FAX1 in pollen tube germination. When we analyzed *in vitro* pollen tube germination of mature pollen grains from heterozygous *fax1-1* and *fax1-2* mutant lines, we found a germination capacity of less than 50 % compared to Col-0 wild-type pollen. Here 27.9 %, 23.8 %, and 58.8 % of pollen grains from *fax1-1*, *fax1-2* and Col-0 were able to germinate pollen tubes *in vitro* (Fig. 7B). Thus, the loss of FAX1 in 50 % of the haploid mutant pollen grains appears to be sufficient to constrain *in vitro* pollen germination completely. Interestingly, we found that *FAX1* belongs to those genes, which are upregulated when pollen tubes are germinated under “semi *in vivo”* conditions (Qin et al., 2009). In the latter study, *FAX1* transcripts increased 3.7 and 3.8-fold when the transcriptome of “semi *in vivo”* pollen germination assays was compared to that of dry pollen grains and *in vitro*-grown pollen tubes, respectively (Fig. 7C). Again, *FAX1* represents the only plastid-localized *FAX* gene differentially expressed in pollen tube assays, and furthermore closely correlates with the expression profile of *FAX6*. The most strongly regulated *FAX* gene in germinating pollen tubes, however, is *FAX5*, which shows highly increased transcript content in “semi *in vivo”*, but also under *in vitro* pollen tube germination conditions (Fig. 7C). Thus, gene expression of *FAX1* and *FAX6* appears to be regulated by pistil–pollen tube specific interactions, which are possible in “semi *in vivo”* assays, while that of *FAX5* in general seems to be important in growing pollen tubes.

### FAX3 complements for the loss of FAX1 in rosette leaf and embryo/seed development

A potential complementary role of FAX1 and FAX3 in rosette leaf development (see above) in the following was investigated by generating *fax1-2/fax3-1* double mutant lines (*fax1/3* dm). Although analyzed by numerous independent approaches and in different filial generations we, however, were not able to raise adult plants homozygous (ho) for both T-DNA insertions *fax1-2* and *fax3-1* (*fax1/3* dm_ho/ho). Thus, in the following we examined segregation of the heterozygous (he) *fax1-2* allele in the background of the homozygous *fax3-1* mutation in two independent lines (*fax1/3* dm_he/ho, lines #4y10 and #5r10). For control, we compared segregation of *fax1-2* in the wild-type background, i.e. in the heterozygous *fax1-2* single mutant line. Further, all mutant lines were germinated and grown on two different media, specifically on agar and soil. As published previously and due to the male sterility described (Li et al., 2015), Mendelian segregation of the single *fax1-2* allele with only 4.2% and 5.4% of homozygous progeny on agar and on soil, respectively, was severely impaired (Table 1). Transmission of *fax1-2* in the background of the *fax3* knockout, however, was even less successful. While we failed to detect any *fax1/3* dm_ho/ho plantlets in line #5r10, we could identify 2.8% double homozygous seedlings on agar medium in line #4y10 (Table 1). Interestingly, this rate dropped to 0.6% - explicitly only one ho/ho plant identified in 170 analyzed - when plantlets were germinated on soil.

**Table 1.** Segregation analysis of the fax1-2 mutant allele in the background of Col-0 wildtype and the homozygous fax3-1 knockout mutation.

|  | n.p. [%] | no. | ho [%] | he [%] | wt [%] |
| --- | --- | --- | --- | --- | --- |
| **agar** |  |  |  |  |  |
| *fax1-2_he* | 5.6 | 120 | 4.2 | 56.7 | 39.2 |
| *fax1/3 dm_he/ho* #4y10 | 3.4 | 141 | 2.8 | 49.6 | 47.5 |
| *fax1/3 dm_he/ho* #5r10 | 8.2 | 132 | 0 | 59.8 | 40.2 |
| **soil** |  |  |  |  |  |
| *fax1-2_he* | 8.7 | 112 | 5.4 | 60.7 | 33.9 |
| *fax1/3 dm_he/ho* #4y10 | 5.5 | 170 | 0.6 | 59.4 | 40.0 |
| *fax1/3 dm_he/ho* #5r10 | 4.7 | 127 | 0 | 52.8 | 47.2 |

The progeny of heterozygous (he) single *fax1-2* mutants and of *fax1-2* (he)/*fax3-1* (ho) double mutants (*fax1/3 dm*) was germinated on agar medium or on soil and analyzed after three weeks of growth by PCR-genotyping. For *fax1/3 dm\_he/ho* two independent lines #4y10 and #5r10 were tested. The genotype of homozygous (ho), heterozygous (he), and wild-type (wt) *fax1-2* mutant alleles is given in percent. n.p. [%], percentage of seeds that did not germinate or develop into seedlings ("no plantlet"); no., number of plantlets analyzed by PCR-genotyping.

As described earlier (Li et al., 2015), *fax1-2* single mutant plants were smaller than wildtype but appeared not to be delayed in development. The phenotype of *fax1/3* dm_ho/ho grown on soil, however, showed an arrest in growth and development of rosette leaves. Here, the 3-week-old plant of *fax1/3* double ko mutant line #4y10 had unfolded green cotyledons but failed to grow any rosette leaves (Fig. 8A), while the corresponding wild-type plants contained four fully developed rosette leaves and were about seven times bigger (Fig. 8C). When grown on agar medium, the *fax1/3* dm_ho/ho seedlings managed to develop first rosette leaves (Fig. 8B), but still were about two to threefold smaller than wild-type plantlets (Fig. 8D). Transfer of these *fax1/3* dm_ho/ho seedlings to soil - after germination of seeds on agar medium -, however, did not result in any surviving double ko line, in contrast to plants with heterozygous or wild-type genotypes for *fax1-2*. Therefore, we can conclude that *fax1/3* double ko plants in general are not able to develop into mature rosette stages and are seedling lethal when grown on soil.

**Fig. 8.**
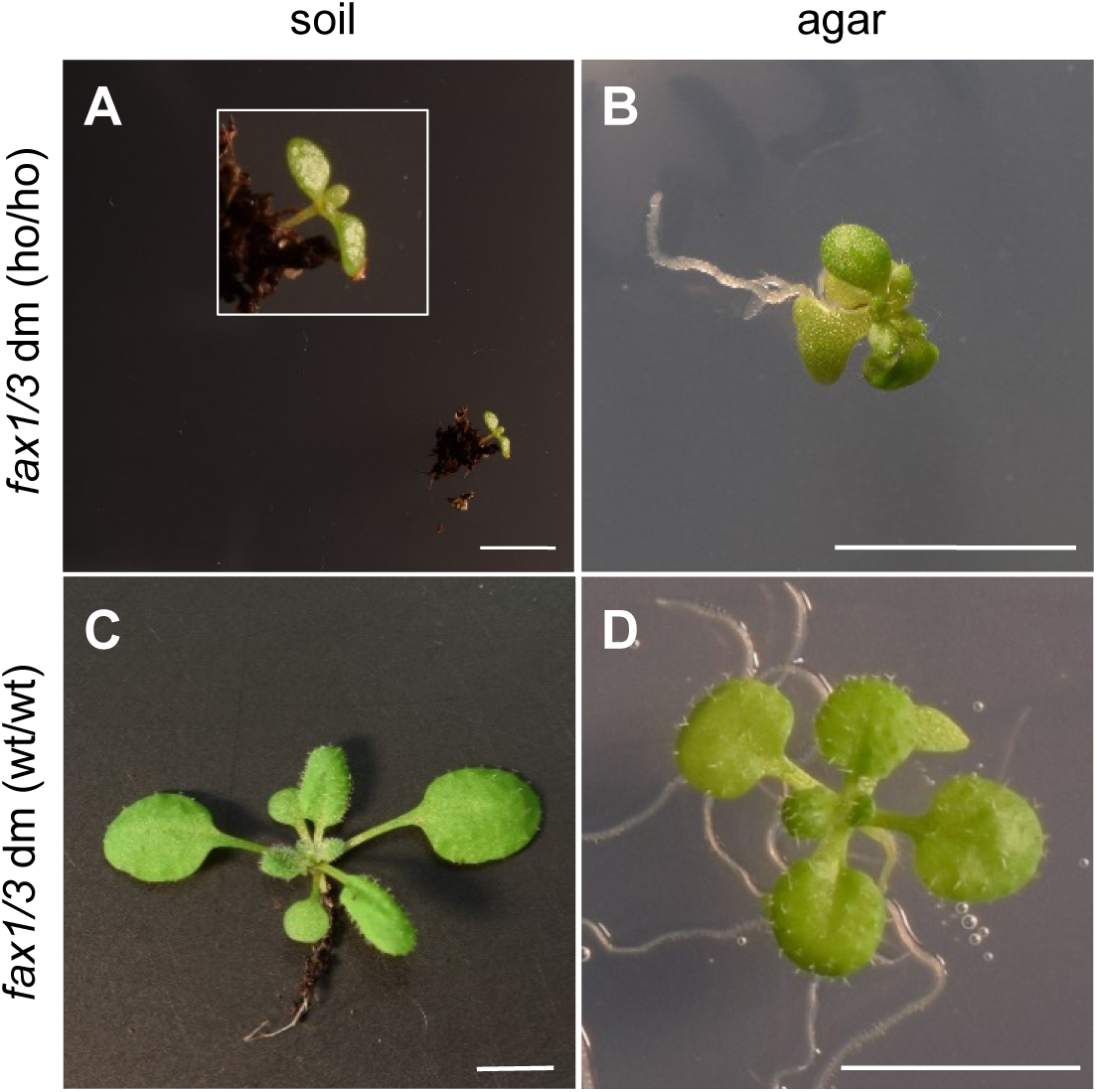
The double knockout of FAX1 and FAX3 leads to an arrest in rosette leaf growth. 3-week-old plantlets of the *fax1-2*, *fax3-1* double mutant (*fax1/3* dm), line #4y10 grown on soil (A, C) and agar medium (B, D). (A), (B) Mutants, homozygous for both *fax1-2* and *fax3-1* alleles (*fax1/3* dm_ho/ho). (C), (D) Plants with corresponding segregated double wild-type alleles for *fax1-2* and *fax3-1* (*fax1/3* dm_wt/wt). Please note that the width of the cotyledons of *fax1/3* dm_ho/ho on soil in (A) is around 0.37 cm, while the rosette leaf diameter of the corresponding wildtype is about 2.7 cm. Inset in (A), magnification of double knockout seedling. Scale bar = 5 mm.

Further, from the data presented in Table 1 we can assume the following for gametophyte and embryo/seed development of *fax1/3* double ko plants: (i) Line #5r10 (*fax1/3* dm_he/ho) did neither on agar nor on soil produce any double homozygous progeny and therefore displays a strong, lethal phenotype in most likely either gametophyte or embryo/seed developmental stages. (ii) Line #4y10 (*fax1/3* dm_he/ho) generated only very few double homozygous seedlings on agar (2.8%) and on soil (0.6%). (iii) Thus, on agar medium with sucrose supplementation most likely seed germination and seedling development until an early rosette stage of *fax1/3* dm_ho/ho is not impaired. In conclusion we can assume that around 2.8% embryos/seeds for *fax1/3* dm are double homozygous knockouts for *FAX1* and *FAX3*. (iv) On soil, approximately 2.2% of these *fax1/3* dm_ho/ho seeds do not germinate or develop into seedlings and only 0.6% of produce seedlings, which are arrested in the cotyledon stage (see above and Fig. 8A).

In comparison to the segregation of ho *fax1-2* in Col-0 background - i.e. about 4.2 to 5.4% (compare Table 1 and Li et al. [2015]) - the above extrapolated 2.8% occurrence of double homozygous *fax1/3* ko seeds is significantly reduced. In consequence, the missing 1.4 to 2.6% of *fax1/3* dm_ho/ho progeny when compared to *fax1* single mutants most likely are due to the loss of FAX3 function either in gametophyte or in embryo/seed development. A normal Mendelian segregation of *fax3-1* single ko mutants (Supplementary Fig. S10D) and rather low gene expression of *FAX3* in pollen stages (see Fig. 7) as well as the predominant role of FAX1 in pollen development (Li et al., 2015, Zhu et al., 2020) and pollen tube growth (see Fig. 7B,C), argue against a crucial function of FAX3 in gametophyte development. In contrast, *FAX3* transcript levels are high in embryo and seed development (Fig. 9A). Here *FAX3* expression is strongest in all stages, in particular in late seed development and dry seeds and does not cluster with that of any other *FAX* gene. Only when seed germination is initiated by imbibition, FAX3 appears to be supported by increased gene expression of *FAX1, FAX2,* and *FAX4*. Therefore, in combination with the knockout of *FAX1*, FAX3 loss-of-function might impair normal maturation of embryos and/or seeds in *fax1/3* dm. To test the latter hypothesis, we examined development of seed grains in mature green siliques of *fax1/3* dm plants from line #4y10 (Fig. 9B). We determined the total number of seeds per silique as well as the amount of empty white seed coats and gaps in siliques of *fax1/3* dm_he/ho and *fax1/3* dm_wt/ho plants (Fig. 9C). Here, siliques from *fax1/3* dm_he/ho (mean of 40 seeds, 1.80 gap per silique) contained about two seeds less than those from *fax1/3* dm_wt/ho plants (mean of 41.8 seeds, 1.07 gap per silique). Further, *fax1/3* dm_he/ho siliques also produced about two times more empty white seed coats (mean of 0.93) without fully developed embryos than *fax1/3* dm_wt/ho plants (mean of 0.43), which represent the *fax3* single mutant background. Thus, it is very likely that the knockout of both *FAX1* and *FAX3* results in defects in embryo and/or seed development.

**Fig. 9.**
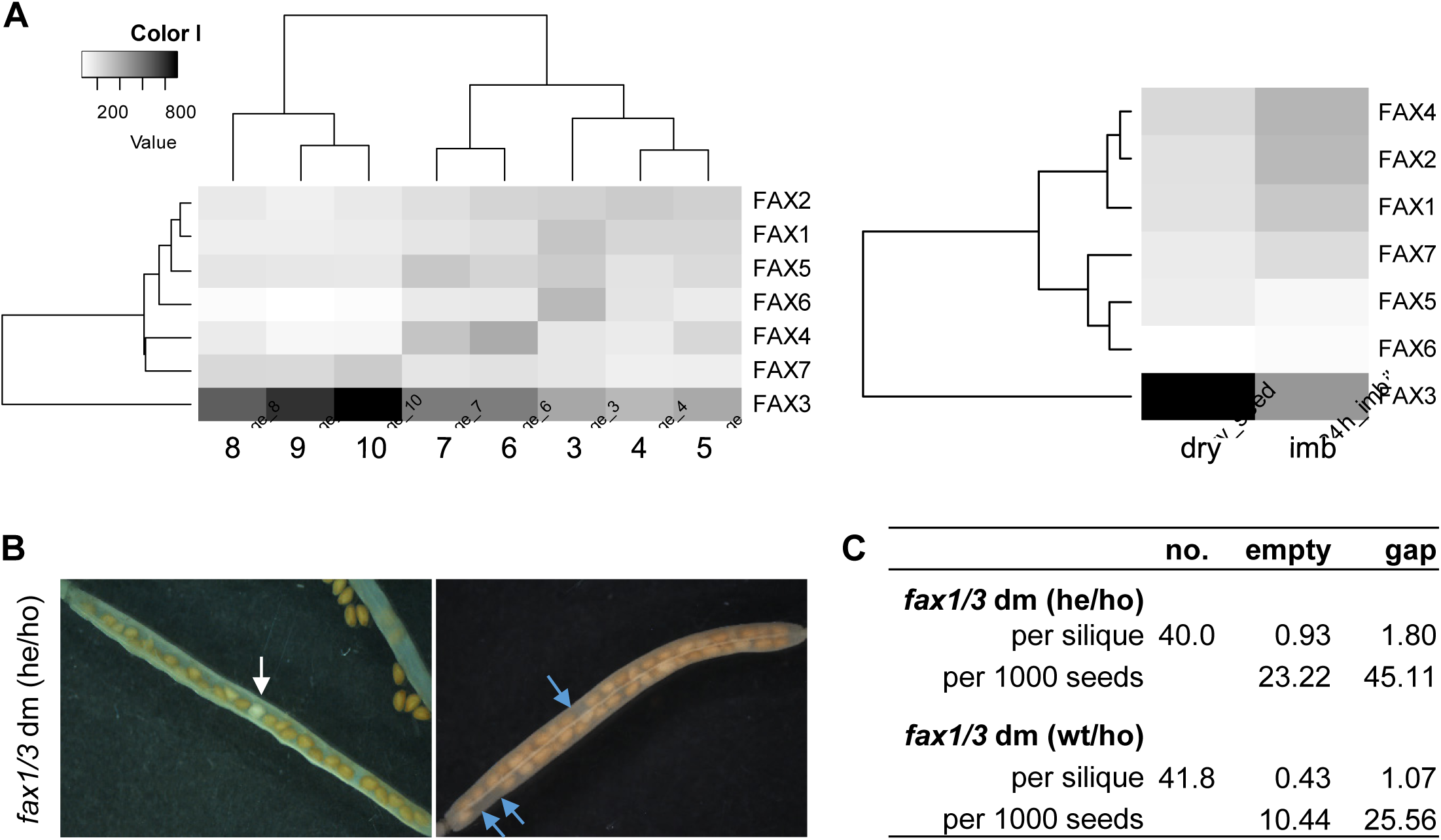
The double knockout of FAX1 and FAX3 leads to impaired seed/embryo development. (A) Correlation of gene expression of *Arabidopsis FAX* genes during seed development. Heatmaps depict gene expression during seed development (left) as well as dry and imbibed (24 h in water to induce germination) seeds (right). Gene expression intensities are displayed in grayscale as shown in the color key (the darker the shades of grey, the higher the gene expression). *FAX* genes are in rows, whereas seed stages are organised in columns. Gene expression data used to create heatmaps are based on DNA microarray analyses obtained from AtGenExpress developmental series (Schmid et al., 2005). Left, seed stages 3–10 are defined according to embryo development as follows: 3: mid-globular to early heart; 4: early heart to late heart; 5: late heart to mid-torpedo; 6: mid-torpedo to late torpedo; 7: late torpedo to early walking stick; 8: walking stick to early curled cotyledons; 9: curled cotyledons to early green cotyledons; 10: green cotyledons. Note that stages 3–5 include silique tissue. (B) Representative pictures of mature green siliques from the *fax1-2*, *fax3-1* double mutant (*fax1/3* dm, line #4y10), heterozygous for *fax1-2* and homozygous for *fax3-1* alleles (*fax1/3* dm_he/ho). Left, half of opened green silique to visualize white seed coats. Right, destained intact mature silique to detect gaps without seeds. White empty seed coats with undeveloped embryos are indicated by white arrows, gaps without any seeds are highlighted by blue arrows. (C) Quantification of the total seed number per silique (no.), white empty seed coats with undeveloped embryos (empty), and gaps without any seeds in mature siliques of *fax1/3* dm_he/ho (see [B]), and of the corresponding *fax1-2* wild-type allele *fax1/3* dm_wt/ho. Data represent mean values of 56 siliques from 6 individual plants of *fax1/3* dm_he/ho, and of 44 siliques from 5 different plants of *fax1/3* dm_wt/ho.

In summary, we could show that the simultaneous loss of both FAX1 and FAX3 proteins leads (i) to defects in seed and/or embryo development, (ii) to reduced seed germination and/or seedling growth on soil as well as (iii) to an arrest in rosette leaf growth. In these developmental stages, *fax1* single ko mutants, however, only showed defects in seed germination/seedling growth on soil in comparison to sugar-supplemented agar (8.7% and 5.6% without germinated seeds/developed seedling, respectively, see Table1). Thus, we can conclude that in embryo/seed development as well as in rosette leaf growth, function of FAX3 most likely complements for that of FAX1.

## Discussion

### The plant FAX protein family

Based on *Arabidopsis* proteins and on comparison with various plant species we here define the plant FAX-protein family, which contains six subfamilies consisting of FAX1-, FAX2-, FAX3-, FAX4-, FAX5/6-, and FAX7-like proteins. With only little sequence similarity between these subgroups (< 39% aa identity), At-FAX5 and At-FAX6 with 81% identical amino acids appear to be recently developed paralogs and were grouped in one subfamily. Also, identical gene organization in exon/intron regions of At-FAX5 and At-FAX6 as well as the fact that only in few plant species two gene copies for FAX5 and FAX6 are present (see Könnel et al. [2019], Peter et al. [2022]), points to a recent gene duplication event for *At-FAX5* and *At-FAX6*. Genes coding for FAX1, FAX3, FAX5/6 and FAX7 proteins can be found in the clade of *Viridiplantae* ranging from green microalgae (i.e. *Chlamydomonas reinhardtii*, see Peter et al. [2022]) to angiosperm plants. FAX1 proteins with a FAX2-like apolipoprotein N-terminus are present in moss and spikemoss (see below), but a separate FAX2-subfamily appears to be present in seed plants only. FAX4-like proteins can be found in *Selaginella moellendorfii* and *Physcomitrium patens*, but not in *Chlamydomonas*. Further, FAX4 proteins are present in monocots like rice and maize but not in several dicot species where we identified relatives of all other plastid-localized FAX proteins (compare Supplementary Fig. S5). A clear statement on the origin and evolution of FAX proteins in plants, however, is only possible with a distinct phylogenetic analysis, which is not part of the current study that is based on conserved sequences and structural motifs. Common to all plant FAX proteins and to their vertebrate TMEM14 relatives is the membrane-intrinsic Tmemb_14 domain of four α-helices of which at least two are hydrophobic and no less than one is amphiphilic (Fig.1). Within the Tmemb_14 domain we were able to identify several motifs, which are conserved and are specific for each FAX subfamily. Here FAX1, FAX2, FAX4, and FAX5/6 as well as FAX3 and FAX7 form two subgroups with similar distribution of polar and hydrophobic helix faces, charged amino acid residues and conserved sequence motifs. In particular, the first α-helix of plastid-localized FAX1 and FAX3 is almost identical to the ER/secretory pathway predicted FAX5/6 and FAX7, respectively (see also (Könnel et al., 2019, Peter et al., 2022) for FAX1 and FAX5/6). Specific for FAX1 proteins is the highly conserved GGNPPKK-motif, C-terminal of the Tmemb_14 domain. According to our topology model (compare Fig. 4B), this motif points to the IMS. In an evolutionary perspective and based on the conserved sequence motifs, we thus can assume that the basic set of plant FAX proteins consists of FAX1 and FAX3 in plastids as well as of FAX5/6 and FAX7 in ER/secretory pathway membranes.

### Structural features of the plastid IE-intrinsic FAX1, FAX2 and FAX3 proteins

In addition to FAX1 (Li et al., 2015), we here for the first time could unequivocally localize FAX2 and FAX3 proteins by *in vivo* GFP-targeting and immunoblot analysis to the IE membrane of chloroplasts. While GFP-targeting was performed with *Arabidopsis* FAX proteins in tobacco protoplasts, the isolation of the coding sequences of FAX2 and FAX3 from pea mRNA, allowed immunoblot analysis on purified membranes from pea chloroplasts. Our results are supported by predicted N-terminal chloroplast targeting peptides, the presence in proteomic analysis of chloroplast membranes (Ferro et al., 2010, Bouchnak et al., 2019), and the targeting of FAX2-GFP to chloroplast envelopes in transiently transformed *Arabidopsis* protoplasts (Tian et al., 2019, Li et al., 2020). By proteolysis of IE membrane vesicles from pea chloroplasts, followed by immunoblot analysis, we further could develop a topology model for FAX1, FAX2, and FAX3 in the IE lipid bilayer membrane (see Fig. 4B). Because FAX4 relatives in pea are absent, immunoblotting and topology analysis on pea chloroplast membranes is impossible. For FAX4 localization we thus have to rely on *in vivo* GFP-targeting in *Arabidopsis* protoplasts, which points to a chloroplast envelope localization as well (Li et al., 2020). Since for all FAX4 proteins identified, a classical NT-terminal and cleavable chloroplast transit peptide is very likely, targeting to the IE membrane is to be expected for FAX4 as well. As described in (Könnel et al., 2019) for FAX1, we propose that the amphiphilic helices in the Tmemb_14 domain of plastid IE-intrinsic FAX1, FAX2, FAX3, and FAX4 proteins in combination with the conserved positively and negatively charged amino acid residues at the helix rims, can create a certain asymmetry to the IE lipid bilayer membrane. Helix 3 of FAX1 and FAX2 according to (Gkeka and Sarkisov, 2010) is a type II amphiphilic helix with polar and hydrophobic faces of either similar size or a larger polar face along the vertical helix axis (see Fig.1). These so-called “classical” amphiphilic helices are known to bind lipids and to lipid membranes by plunging into the bilayer parallel to the membrane surface. Helix 1 of FAX1 and helices 1 and 2 of FAX3 instead represent type III amphiphilic helices where hydrophobic residues are distributed in a conical fashion, i.e. the hydrophilic residues are clustered only one side of the vertical helix axis in this case at the N-terminal end (compare Fig. 1). This generates membrane-spanning helices that are tilted vertically inside the membrane. For FAX3, therefore, a hairpin like arrangement of helices 1 and 2 is proposed. In consequence this membrane topology of FAX1, FAX2, and FAX3 (see Fig. 4B), most likely stabilizes and/or shapes the IE membrane and thereby might contribute to shuttling fatty acids across the IE (for further discussion see Könnel et al. [2019]). Compact insertion within the lipid bilayer membrane and strong binding of FAX proteins to lipids further could explain why the apparent molecular weight of FAX1, FAX2, FAX3 on SDS-PAGE is about 5 kDa larger than the calculated mass. Although a topology prediction for FAX4 was not possible, distribution of conserved motifs in the Tmemb14 domain suggest an integration into the lipid bilayer membrane similar to FAX1 and FAX2.

Remarkably the mature, plastid IE-localised proteins FAX1, FAX2 and FAX3 contain extended polypeptide stretches N-terminal of the Tmemb_14 domain with motives and features specific for the respective FAX subfamily (see Fig. 2A). The N-terminal region of <u>FAX1</u> proteins appears to be in the plastid stroma and seems to fold into an additional non membrane-associated α-helix (compare Li et al. [2015]) with a function that has yet to be determined. However, this N-terminal helix in the stroma and the C-terminal GGNPPK-motif of FAX1 in the IMS for example could be involved in mediating interaction with still unknown partner proteins. The most prominent N-terminal domain of FAX proteins, however, is the apolipoprotein-like α-helical bundle of <u>FAX2</u> proteins. Similar to the helical bundles in the mammalian apolipoproteins apoE and apoA-I as well as in the insect apolipophorin III (Narayanaswami et al., 2010, Huebbe and Rimbach, 2017), we could determine four amphiphilic α-helical cylinders in the N-terminus of FAX2, which display a hydrophobic face along the vertical axis that separates from patterned charged amino acid residues on the hydrophilic side of the helix (Fig. 2C, D). In addition, structural modelling of the entire N-terminal region of At-FAX2 reveals structural similarity with the helical bundle of human apolipoprotein e4 (Fig. 2B). The hydrophobic faces of mammalian apolipoprotein helices form a hydrophobic core in the helical bundle formation that reversibly unfolds and refolds in aqueous solution. In contact with lipid bilayers or liposomes, these helical bundles open up and thereby manage lipid solubilization during HDL biogenesis by producing disc-shaped HDL particles of around 10nm diameter, which enclose a phospholipid bilayer segment (Phillips, 2013). In plants, however, this class of apolipoproteins with α-helical bundles has not been described so far. Further, to our knowledge FAX2, with its C-terminal Tmemb_14 domain in the IE membrane, represents a novel and unique class of membrane-intrinsic proteins with an attached apolipoprotein-like α-helix bundle. Due to our model (Fig. 4B), this N-terminal α-helical bundle of FAX2 points to the IMS. Therefore, we propose that this domain might be able to mediate membrane contacts between OE and IE by solubilizing lipid bilayer membranes and/or forming lipid nanodisc-like structures similar to mammalian apolipoproteins during HDL biogenesis. Thereby FAX2 proteins would be able to facilitate fatty acid/lipid transfer across the two plastid envelope membranes. Moreover, the punctuate structure of FAX2-GFP signals (see Fig. 3C and Tian et al. [2019]) could be explained by the formation of larger complexes containing lipid nanodisc-like structures. A similar punctuate GFP-pattern in chloroplasts for example is observed for VIPP1 (Aseeva et al., 2004), which forms large ring- or rod-shaped homo-oligomeric complexes that in their lumen bind lipids or liposomes via an N-terminal amphiphilic α-helix (Gupta et al., 2021). Further, the punctuate FAX2-GFP signals could also point to membrane contacts for trafficking of lipid compounds between the chloroplast envelope and the ER (Block and Jouhet, 2015). Remarkably, FAX2 relatives are present in dicot and monocot angiosperm plants but are absent in the green microalga *Chlamydomonas reinhardtii*. In moss and spikemoss (*Physcomitrium patens* and *Selaginella moellendorffii*), however, proteins with a FAX1-like Tmemb_14 domain and the C-terminal GGNPPKK-motif contain the FAX2-like apolipoprotein N-terminus. Thus, most likely FAX2 proteins evolved together with seed plants, while during the transfer from water (green microalgae) to early land plants (mosses, spikemoss), the apolipoprotein α-helical bundle appeared and was linked to a FAX1-like, membrane-intrinsic Tmemb_14 C-terminus. All <u>FAX3</u> proteins in their N-terminus, however, contain a remarkable region (polyG stretch) comprising three conserved motives with glycine repeats (Fig 2E), which most likely is localized in the plastid stroma (see Fig. 4B). Although in plants several glycine repeat motifs are described in glycine-rich proteins (Czolpinska and Rurek, 2018), the FAX3-specific polyG repeats are unique since they can contain up to seven consecutive glycine residues (e.g. in At-FAX3) and are accompanied by also conserved patterns of negatively charged amino acids. Among the few polyG motifs with demonstrated function, one is found in the targeting signal for the chloroplast OE protein translocon channel Toc75. Here this motif - also five glycines in a row - is necessary for proper envelope sorting during chloroplast import of the preprotein Toc75 (Endow et al., 2016). However, the Toc75 polyG domain, which according to Endow and co-workers is proposed to act as a rejection signal for Toc75 at the chloroplast IE, after import and sorting is split off and thus is not present in the mature, functional Toc75 ß-barrel pore. For FAX3 in contrast, we could detect the polyG repeats at the N-terminus of the mature protein in the IE membrane. Since glycine-repeats in proteins often are involved in mediating protein-protein interactions (Mangeon et al., 2010, Czolpinska and Rurek, 2018), the small glycine residues in FAX3 polyG could simply function as intrinsically disordered, flexible linker region for mediating interactions of the FAX3 N-terminus, e.g. via the conserved and negatively charged amino acid residues. Only recently, a study with glycine-rich peptides, which form intrinsically disordered structures and include poly-Gly motifs, showed that 10 consecutive glycine residues are necessary for self-assembly of the peptides into fibrillar structures (Kar et al., 2021). Further, Kar and co-workers conclude that the transition from a soluble to a strongly self-associating system occurs between polyG length between 6-10 residues. Thus, it is tempting to speculate that the polyG motif at the N-terminus of FAX3 proteins can mediate self-assembly of FAX3 proteins and/or interaction with protein partners in the plastid stroma.

In membrane-spanning α-helices, glycine-repeat motifs as well are described to mediate protein oligomerization via helix-helix interactions and organization of membrane topology (Teese and Langosch, 2015). Specifically, the Mic10 protein in the inner membrane of mitochondria has been shown to oligomerize via its GxxxG motifs in its two transmembrane helices (Barbot et al., 2015). Due to the presence of three conserved positively charged amino acids in the short loop between the two α-helices of Mic10, the protein captures a hairpin-like formation in the lipid bilayer. In consequence, this topology and the homo-oligomerization of Mic10 are prerequisites for membrane curvature in the cristae junctions of inner mitochondrial membranes (Barbot and Meinecke, 2016, Könnel et al., 2019). Further, the two predicted IE transmembrane domains of the plastid IMS-localized TGD5 protein also contain glycine-rich repeats (Fan et al., 2015). Here Fan and co-workers speculate that these motifs contribute to TGD5 homo-oligomerization and thereby to assembly of large TGD protein complexes by bridging the TGD1-3 ABC transporter at the plastid IE with the TGD4 ß-barrel at the OE for import of eukaryotic lipids into chloroplasts (LaBrant et al., 2018). Interestingly, the first transmembrane α-helices in the Tmemb_14 domain of plastid IE intrinsic FAX1, FAX2 and FAX3 proteins as well contain conserved glycine-repeat motifs (see Supplementary Figs. 2-4). Thus, it is tempting to speculate that these glycine repeats contribute to homo- and/or hetero-oligomerization of FAX1, FAX2, and FAX3. With regard to the proposed hairpin- and Mic10-like arrangement of helix 1 and helix 2 of the Tmemb_14 domain of FAX3 (see Fig. 4B), a function in membrane-shaping and stability is possible as well.

In summary, we conclude that the specific sequence and structural features of plastid-localized FAX1, FAX2, and FAX3 proteins in their C-terminal Tmemb_14 domains and in their particular N-terminal stretches could contribute to protein-protein interactions as well as IE membrane shaping and/or stability. Most likely thereby plastid FAX proteins might be able to mediate membrane contacts between IE and OE and to shuttle fatty acids across the plastid envelope via a vectorial acylation mechanism (see also Li et al. [2016], Li-Beisson et al. [2017], Könnel et al. [2019]). As demonstrated in our current study (see Fig. 6) as well as by (Li et al., 2015, Tian et al., 2019) for At-FAX1 and At-FAX2, respectively, the plastid localized *Arabidopsis* FAX1-FAX4 proteins by this mechanism are able to complement the function of the yeast FA-transport protein Fat1. Here most likely, vectorial acylation is enabled by interaction of FAX proteins with the yeast fatty acyl Co-A synthetases Faa1, Faa4. In chloroplasts, the preferred long-chain acyl CoA synthetase for interaction would be the OE-attached LACS9 as already proposed by (Tian et al., 2019) for At-FAX2.

### The function of plastid IE-intrinsic FAX1 and FAX3 in plant growth and development

In addition to the well-documented function of FAX1 in tapetum cell plastids during formation of the lipophilic outer pollen cell wall (Li et al., 2015, Zhu et al., 2020), we here could show that FAX1 function is also required for germination and growth of pollen tubes. Due to expression profiles (see Fig. 7), *At-FAX1* is the only plastid-FAX protein with strong gene expression in pollen development and pollen tubes. Also signals of FAX1 promoter::GUS lines in addition to tapetum cells point to expression in microspores (Zhu et al., 2020). Further, FAX1 peptides were identified in a proteomic analysis of mature *Arabidopsis* pollen grains (Grobei et al., 2009), indicating that FAX1 represents a relevant protein that is stored in mature pollen grains to be ready for rapid pollen germination and tube growth upon pollen grain hydration. Growing pollen tubes need lots of energy as well as phospholipid material for the expanding pollen tube plasma membrane. Therefore, it seems likely that part of these energy metabolites and FA/lipid compounds are delivered by pollen-intrinsic plastids, either already during pollen grain development or in the vegetative cell of growing pollen tubes (Selinski and Scheibe, 2014, Ischebeck, 2016). Further, very long chain fatty acids from the tapetum-made pollen coat have been described to function in pollen grain hydration (Zhan et al., 2018). We thus conclude, that besides its role in tapetum plastids, FAX1 is crucial for export of FAs from plastids in pollen grains and/or pollen tubes and in consequence for germination and growth of pollen tubes.

In *Arabidopsis* embryo and seed development FAX2 and FAX3 appear to be the dominant plastid FAX proteins expressed (Li et al., 2015, Tian et al., 2019). While the study by Tian and co-workers (2019) demonstrates the pivotal task of At-FAX2 in seed TAG production, we here checked for the role of FAX3 in seed and embryo development of *fax1/3* double ko in comparison to the respective single ko mutants. Although not very pronounced, we could show that the simultaneous loss of FAX1 and FAX3 function leads to impaired embryo and/or seed development. Due to the still possible functional complementation by plastid-localized FAX2 and FAX4, which are also expressed during embryo and seed development, triple our even quadruple mutants of *FAX1-4* genes might be required to unravel stronger effects. In vegetative growth, however, strong expression of *At-FAX1* and *At-FAX3* genes nicely correlates and *FAX3* transcripts increased in rosette leaves of *fax1* ko. In consequence, *fax1/3* double ko mutants display an arrest in rosette leaf growth that leads to a seedling lethal phenotype. Most likely, the simultaneous loss-of-function of chloroplast IE-intrinsic FAX1 and FAX3 proteins during leaf growth and development prevents the export of FAs from chloroplasts and thus leads to a lethal interruption of acyl lipid production for energy storage and lipid bilayer membranes. Although somewhat different in their structural features (see before), both FAX1 and FAX3 proteins demonstrated FA-transport function in yeast, and thus are likely to complement each other in FA-export from chloroplasts. If they act alone as separate FA-transport proteins, in protein complexes with yet unknown partners or even might form FAX hetero-oligomers by interaction for example via the glycine repat motifs in their membrane-spanning α-helices, still has to be clarified. Evidence for a potential function of FAX1 together with FAX3 in larger membrane protein complexes comes from co-migration analysis of peptides on blue-native protein gels. Thereby, we could spot FAX3 to run together with FAX1 in an approximately 230 kDa complex from *Arabidopsis* chloroplasts (Lundquist et al., 2017). Further, we found FAX1, FAX2, and FAX3 to co-migrate in isolated *Arabidopsis* chloroplast envelope membranes in the PCoM database (Takabayashi et al., 2017).

## Abbreviations

CoA: coenzyme A
ER: endoplasmic reticulum
FA: fatty acid
HDL: high density lipoprotein
ho: homozygous
he: heterozygous
IE: chloroplast inner envelope membrane
IEV: IE vesicles
IMS: intermembrane space
ko: knockout
OE: chloroplast outer envelope membrane
TAG: triacylglycerol

## Supplementary data

**Table S1.** Features of FAX family proteins in *Arabidopsis* and pea.

**Table S2.** List of oligonucleotides used in this study.

**Figure S1.** Amino acid sequence alignment of mature *Arabidopsis* FAX proteins.

**Figure S2.** Amino acid sequence alignment of mature plant FAX1 proteins.

**Figure S3.** Amino acid sequence alignment of mature plant FAX3 proteins.

**Figure S4.** Amino acid sequence alignment and motif consensus prediction of mature plant FAX2 proteins.

**Figure S5.** Amino acid sequence alignment of mature plant FAX4 proteins.

**Figure S6.** Helical wheel analysis of all four membrane-intrinsic α-helices predicted for FAX protein subfamilies.

**Figure S7.** Amino acid sequence alignment of plant FAX7 proteins.

**Figure S8.** Amino acid sequence alignment of mature FAX1 proteins in moss and spikemoss.

**Figure S9.** Amino acid sequences of mature FAX1, FAX2, and FAX3 proteins from *Arabidopsis* and pea.

**Figure S10.** Characterization of T-DNA insertion lines for *At-FAX2*, *At-FAX3*, and *At-FAX4*.

**Figure S11.** Expression of *Arabidopsis FAX* genes in rosette leaves of FAX2-, FAX3-, and FAX4 loss-of-function mutants.

## Acknowledgements

We gratefully acknowledge excellent technical assistance by Angelika Anna and we thank Eliane Klein for crossing *fax1, fax3* mutants as well as several student assistants for help in PCR-genotyping. Thanks to Frederik Sommer/Michael Schroda for peptide sequencing, to Louis Percifull for co-migration data mining and to Jens Neunzig for critical reading and discussion. Research was funded by the Deutsche Forschungsgemeinschaft (DFG) grants PH73/6-1, PH73/7-1, and in the framework of the International Research Training Group 1830 (IRTG1830) to KP.

## Author contributions

KP conceived the study and accomplished *in silico* protein motif and domain analysis. WB and AK performed protein sequence comparison, helical wheel analysis and qRT PCR. WB performed immunoblot analysis, characterized *fax2, fax3* single and *fax1/3* double mutants, conducted segregation analysis and phenotyped *fax1/3*dm. AK executed functional assays in yeast and together with JM characterized *fax4* T-DNA insertion lines. JP conducted proteolysis experiments and developed membrane topology models. AS made GFP-targeting experiments, VH performed pollen tube germination analysis and CS achieved seed analysis in *fax1/3*. ABC analyzed expression profiles of At-FAX genes. KP wrote the manuscript with the help of AK and JP.

## Data Availability

All data supporting the findings of this study are available within the paper and within its supplementary materials published online.

## References

Almagro Armenteros, J.J., Salvatore, M., Emanuelsson, O., Winther, O., von Heijne, G., Elofsson, A. and Nielsen, H. (2019) Detecting sequence signals in targeting peptides using deep learning. Life Science Alliance, 2, e201900429.

Ariizumi, T. and Toriyama, K. (2011) Genetic Regulation of Sporopollenin Synthesis and Pollen Exine Development. Annual Review of Plant Biology*, Vol* 62, 62, 437–460.

Aseeva, E., Ossenbuhl, F., Eichacker, L.A., Wanner, G., Soll, J. and Vothknecht, U.C. (2004) Complex formation of Vipp1 depends on its alpha-helical PspA-like domain. J Biol Chem, 279, 35535–35541.

Bailey, T.L., Johnson, J., Grant, C.E. and Noble, W.S. (2015) The MEME Suite. Nucleic Acids Res, 43, W39–49.

Baker, A., Carrier, D.J., Schaedler, T., Waterham, H.R., van Roermund, C.W. and Theodoulou, F.L. (2015) Peroxisomal ABC transporters: functions and mechanism. Biochem Soc Trans, 43, 959–965.

Barbot, M., Jans, D.C., Schulz, C., Denkert, N., Kroppen, B., Hoppert, M., Jakobs, S. and Meinecke, M. (2015) Mic10 oligomerizes to bend mitochondrial inner membranes at cristae junctions. Cell metabolism, 21, 756–763.

Barbot, M. and Meinecke, M. (2016) Reconstitutions of mitochondrial inner membrane remodeling. Journal of structural biology, 196, 20–28.

Bates, P.D. (2016) Understanding the control of acyl flux through the lipid metabolic network of plant oil biosynthesis. Biochim Biophys Acta, 1861, 1214–1225.

Beisson, F., Li-Beisson, Y. and Pollard, M. (2012) Solving the puzzles of cutin and suberin polymer biosynthesis. Curr Opin Plant Biol, 15, 329–337.

Black, P.N. and DiRusso, C.C. (2003) Transmembrane movement of exogenous long-chain fatty acids: proteins, enzymes, and vectorial esterification. Microbiol Mol Biol Rev, 67, 454–472.

Block, M.A. and Jouhet, J. (2015) Lipid trafficking at endoplasmic reticulum-chloroplast membrane contact sites. Current opinion in cell biology, 35, 21–29.

Bouchnak, I., Brugiere, S., Moyet, L., Le Gall, S., Salvi, D., Kuntz, M., Tardif, M. and Rolland, N. (2019) Unraveling Hidden Components of the Chloroplast Envelope Proteome: Opportunities and Limits of Better MS Sensitivity. Mol Cell Proteomics, 18, 1285–1306.

Breuers, F.K., Bräutigam, A., Geimer, S., Welzel, U.Y., Stefano, G., Renna, L., Brandizzi, F. and Weber, A.P. (2012) Dynamic Remodeling of the Plastid Envelope Membranes - A Tool for Chloroplast Envelope in vivo Localizations. Front Plant Sci, 3, 7.

Browse, J., Warwick, N., Somerville, C.R. and Slack, C.R. (1986) Fluxes through the Prokaryotic and Eukaryotic Pathways of Lipid-Synthesis in the 16-3 Plant Arabidopsis-Thaliana. Biochem J, 235, 25–31.

Carrier, D.J., van Roermund, C.W.T., Schaedler, T.A., Rong, H.L., L, I.J., Wanders, R.J.A., Baldwin, S.A., Waterham, H.R., Theodoulou, F.L. and Baker, A. (2019) Mutagenesis separates ATPase and thioesterase activities of the peroxisomal ABC transporter, Comatose. Scientific reports, 9, 10502.

Chen, R., Yamaoka, Y., Feng, Y., Chi, Z., Xue, S. and Kong, F. (2023) Co-Expression of Lipid Transporters Simultaneously Enhances Oil and Starch Accumulation in the Green Microalga Chlamydomonas reinhardtii under Nitrogen Starvation. Metabolites, 13.

Chen, R., Yang, M., Li, M., Zhang, H., Lu, H., Dou, X., Feng, S., Xue, S., Zhu, C., Chi, Z. and Kong, F. (2022) Enhanced accumulation of oil through co-expression of fatty acid and ABC transporters in Chlamydomonas under standard growth conditions. Biotechnol Biofuels Bioprod, 15, 54.

Claus, S., Jezierska, S. and Van Bogaert, I.N.A. (2019) Protein-facilitated transport of hydrophobic molecules across the yeast plasma membrane. Febs Lett, 593, 1508–1527.

Czolpinska, M. and Rurek, M. (2018) Plant Glycine-Rich Proteins in Stress Response: An Emerging, Still Prospective Story. Front Plant Sci, 9, 302.

Dhara, A. and Raichaudhuri, A. (2021) ABCG transporter proteins with beneficial activity on plants. Phytochemistry, 184, 112663.

Do, T.H.T., Martinoia, E. and Lee, Y. (2017) Functions of ABC transporters in plant growth and development. Curr Opin Plant Biol, 41, 32–38.

Duy, D., Stübe, R., Wanner, G. and Philippar, K. (2011) The Chloroplast Permease PIC1 Regulates Plant Growth and Development by Directing Homeostasis and Transport of Iron. Plant Physiol, 155, 1709–1722.

Duy, D., Wanner, G., Meda, A.R., von Wiren, N., Soll, J. and Philippar, K. (2007) PIC1, an ancient permease in Arabidopsis chloroplasts, mediates iron transport. Plant Cell, 19, 986–1006.

Emanuelsson, O., Nielsen, H. and Von Heijne, G. (1999) ChloroP, a neural network-based method for predicting chloroplast transit peptides and their cleavage sites. Protein Sci, 8, 978–984.

Endow, J.K., Rocha, A.G., Baldwin, A.J., Roston, R.L., Yamaguchi, T., Kamikubo, H. and Inoue, K. (2016) Polyglycine Acts as a Rejection Signal for Protein Transport at the Chloroplast Envelope. Plos One, 11, e0167802.

Fan, J., Zhai, Z., Yan, C. and Xu, C. (2015) Arabidopsis TRIGALACTOSYLDIACYLGLYCEROL5 Interacts with TGD1, TGD2, and TGD4 to Facilitate Lipid Transfer from the Endoplasmic Reticulum to Plastids. Plant Cell, 27, 2941–2955.

Ferro, M., Brugiere, S., Salvi, D., Seigneurin-Berny, D., Court, M., Moyet, L., Ramus, C., Miras, S., Mellal, M., Le Gall, S., Kieffer-Jaquinod, S., Bruley, C., Garin, J., Joyard, J., Masselon, C. and Rolland, N. (2010) AT_CHLORO, a comprehensive chloroplast proteome database with subplastidial localization and curated information on envelope proteins. Mol Cell Proteomics, 9, 1063–1084.

Fich, E.A., Segerson, N.A. and Rose, J.K. (2016) The Plant Polyester Cutin: Biosynthesis, Structure, and Biological Roles. Annual review of plant biology, 67, 207–233.

Franssen, S.U., Shrestha, R.P., Brautigam, A., Bornberg-Bauer, E. and Weber, A.P. (2011) Comprehensive transcriptome analysis of the highly complex Pisum sativum genome using next generation sequencing. BMC Genomics, 12, 227.

Gautier, R., Douguet, D., Antonny, B. and Drin, G. (2008) HELIQUEST: a web server to screen sequences with specific alpha-helical properties. Bioinformatics, 24, 2101–2102.

Gkeka, P. and Sarkisov, L. (2010) Interactions of phospholipid bilayers with several classes of amphiphilic alpha-helical peptides: insights from coarse-grained molecular dynamics simulations. J Phys Chem Biol, 114, 826–839.

Grafe, K. and Schmitt, L. (2021) The ABC transporter G subfamily in Arabidopsis thaliana. J Exp Bot, 72, 92–106.

Grobei, M.A., Qeli, E., Brunner, E., Rehrauer, H., Zhang, R., Roschitzki, B., Basler, K., Ahrens, C.H. and Grossniklaus, U. (2009) Deterministic protein inference for shotgun proteomics data provides new insights into Arabidopsis pollen development and function. Genome Res, 19, 1786–1800.

Gupta, T.K., Klumpe, S., Gries, K., Heinz, S., Wietrzynski, W., Ohnishi, N., Niemeyer, J., Spaniol, B., Schaffer, M., Rast, A., Ostermeier, M., Strauss, M., Plitzko, J.M., Baumeister, W., Rudack, T., Sakamoto, W., Nickelsen, J., Schuller, J.M., Schroda, M. and Engel, B.D. (2021) Structural basis for VIPP1 oligomerization and maintenance of thylakoid membrane integrity. Cell, 184, 3643–3659 e3623.

Honys, D. and Twell, D. (2003) Comparative analysis of the Arabidopsis pollen transcriptome. Plant Physiol, 132, 640–652.

Huebbe, P. and Rimbach, G. (2017) Evolution of human apolipoprotein E (APOE) isoforms: Gene structure, protein function and interaction with dietary factors. Ageing Res Rev, 37, 146–161.

Hurlock, A.K., Roston, R.L., Wang, K. and Benning, C. (2014) Lipid trafficking in plant cells. Traffic, 15, 915–932.

Ischebeck, T. (2016) Lipids in pollen - They are different. Biochim Biophys Acta, 1861, 1315–1328.

Jang, S., Kong, F., Lee, J., Choi, B.Y., Wang, P., Gao, P., Yamano, T., Fukuzawa, H., Kang, B.H. and Lee, Y. (2020) CrABCA2 Facilitates Triacylglycerol Accumulation in Chlamydomonas reinhardtii under Nitrogen Starvation. Mol Cells, 43, 48–57.

Kalinger, R.S., Pulsifer, I.P., Hepworth, S.R. and Rowland, O. (2020) Fatty Acyl Synthetases and Thioesterases in Plant Lipid Metabolism: Diverse Functions and Biotechnological Applications. Lipids, 55, 435–455.

Kar, M., Posey, A.E., Dar, F., Hyman, A.A. and Pappu, R.V. (2021) Glycine-Rich Peptides from FUS Have an Intrinsic Ability to Self-Assemble into Fibers and Networked Fibrils. Biochemistry, 60, 3213–3222.

Karimi, M., Inze, D. and Depicker, A. (2002) GATEWAY vectors for Agrobacterium-mediated plant transformation. Trends Plant Sci, 7, 193–195.

Keegstra, K. and Yousif, A.E. (1986) Isolation and characterization of chloroplast envelope membranes. Methods in Enzymology, 118-Plant Molecular Biology, 316–325.

Kelley, L.A., Mezulis, S., Yates, C.M., Wass, M.N. and Sternberg, M.J. (2015) The Phyre2 web portal for protein modeling, prediction and analysis. Nature protocols, 10, 845–858.

Kim, S., Yamaoka, Y., Ono, H., Kim, H., Shim, D., Maeshima, M., Martinoia, E., Cahoon, E.B., Nishida, I. and Lee, Y. (2013) AtABCA9 transporter supplies fatty acids for lipid synthesis to the endoplasmic reticulum. Proc Natl Acad Sci USA, 110, 773–778.

Klammt, C., Maslennikov, I., Bayrhuber, M., Eichmann, C., Vajpai, N., Chiu, E.J., Blain, K.Y., Esquivies, L., Kwon, J.H., Balana, B., Pieper, U., Sali, A., Slesinger, P.A., Kwiatkowski, W., Riek, R. and Choe, S. (2012) Facile backbone structure determination of human membrane proteins by NMR spectroscopy. Nature methods, 9, 834–839.

Könnel, A., Bugaeva, W., Guegel, I.L. and Philippar, K. (2019) BANFF: bending of bilayer membranes by amphiphilic alpha-helices is necessary for form and function of organelles. Biochem Cell Biol, 97, 243–256.

Küchler, M., Decker, S., Hörmann, F., Soll, J. and Heins, L. (2002) Protein import into chloroplasts involves redox-regulated proteins. Embo J, 21, 6136–6145.

LaBrant, E., Barnes, A.C. and Roston, R.L. (2018) Lipid transport required to make lipids of photosynthetic membranes. Photosynthesis research, 138, 345–360.

Lavell, A.A. and Benning, C. (2019) Cellular Organization and Regulation of Plant Glycerolipid Metabolism. Plant Cell Physiol, 60, 1176–1183.

Lee, S.B. and Suh, M.C. (2015) Advances in the understanding of cuticular waxes in Arabidopsis thaliana and crop species. Plant Cell Rep, 34, 557–572.

Li-Beisson, Y., Neunzig, J., Lee, Y. and Philippar, K. (2017) Plant membrane-protein mediated intracellular traffic of fatty acids and acyl lipids. Curr Opin Plant Biol, 40, 138–146.

Li-Beisson, Y., Shorrosh, B., Beisson, F., Andersson, M.X., Arondel, V., Bates, P.D., Baud, S., Bird, D., Debono, A., Durrett, T.P., Franke, R.B., Graham, I.A., Katayama, K., Kelly, A.A., Larson, T., Markham, J.E., Miquel, M., Molina, I., Nishida, I., Rowland, O., Samuels, L., Schmid, K.M., Wada, H., Welti, R., Xu, C., Zallot, R. and Ohlrogge, J. (2013) Acyl-lipid metabolism. The Arabidopsis book / American Society of Plant Biologists, 11, e0161.

Li, N., Gügel, I.L., Giavalisco, P., Zeisler, V., Schreiber, L., Soll, J. and Philippar, K. (2015) FAX1, a novel membrane protein mediating plastid fatty acid export. PLoS Biology, 13, e1002053.

Li, N., Meng, H., Li, S., Zhang, Z., Zhao, X., Wang, S., Liu, A., Li, Q., Song, Q., Li, X., Guo, L., Li, H., Zuo, J. and Luo, K. (2020) Two Plastid Fatty Acid Exporters Contribute to Seed Oil Accumulation in Arabidopsis. Plant Physiol, 182, 1910–1919.

Li, N., Xu, C., Li-Beisson, Y. and Philippar, K. (2016) Fatty Acid and Lipid Transport in Plant Cells. Trends Plant Sci, 21, 145–158.

Li, N., Zhang, Y., Meng, H., Li, S., Wang, S., Xiao, Z., Chang, P., Zhang, X., Li, Q., Guo, L., Igarashi, Y. and Luo, F. (2019) Characterization of Fatty Acid Exporters involved in fatty acid transport for oil accumulation in the green alga Chlamydomonas reinhardtii. Biotechnol Biofuels, 12, 14.

Lundquist, P.K., Mantegazza, O., Stefanski, A., Stuhler, K. and Weber, A.P.M. (2017) Surveying the Oligomeric State of Arabidopsis thaliana Chloroplasts. Mol Plant, 10, 197–211.

Mangeon, A., Junqueira, R.M. and Sachetto-Martins, G. (2010) Functional diversity of the plant glycine-rich proteins superfamily. Plant Signal Behav, 5, 99–104.

Michaud, M. and Jouhet, J. (2019) Lipid Trafficking at Membrane Contact Sites During Plant Development and Stress Response. Front Plant Sci, 10, 2.

Mól, A.R., Castro, M.S. and Fontes, W. (2018) NetWheels: A web application to create high quality peptide helical wheel and net projections. 416347.

Narayanaswami, V., Kiss, R.S. and Weers, P.M. (2010) The helix bundle: a reversible lipid binding motif. Comp Biochem Physiol A Mol Integr Physiol, 155, 123–133.

Peter, J., Huleux, M., Spaniol, B., Sommer, F., Neunzig, J., Schroda, M., Li-Beisson, Y. and Philippar, K. (2022) Fatty acid export (FAX) proteins contribute to oil production in the green microalga Chlamydomonas reinhardtii. Front Mol Biosci, 9, 939834.

Philippar, K., Geis, T., Ilkavets, I., Oster, U., Schwenkert, S., Meurer, J. and Soll, J. (2007) Chloroplast biogenesis: the use of mutants to study the etioplast-chloroplast transition. Proc Natl Acad Sci USA, 104, 678–683.

Phillips, M.C. (2013) New insights into the determination of HDL structure by apolipoproteins: Thematic review series: high density lipoprotein structure, function, and metabolism. J Lipid Res, 54, 2034–2048.

Qin, Y., Leydon, A.R., Manziello, A., Pandey, R., Mount, D., Denic, S., Vasic, B., Johnson, M.A. and Palanivelu, R. (2009) Penetration of the stigma and style elicits a novel transcriptome in pollen tubes, pointing to genes critical for growth in a pistil. PLoS Genet, 5, e1000621.

R-Core-Team (2019) R: A language and environment for statistical computing. R Foundation for Statistical Computing, *Vienna, Austria*, URL https://www.R-project.org/.

Schmid, M., Davison, T.S., Henz, S.R., Pape, U.J., Demar, M., Vingron, M., Scholkopf, B., Weigel, D. and Lohmann, J.U. (2005) A gene expression map of Arabidopsis thaliana development. Nature Genetics, 37, 501–506.

Schwacke, R., Schneider, A., van der Graaff, E., Fischer, K., Catoni, E., Desimone, M., Frommer, W.B., Flugge, U.I. and Kunze, R. (2003) ARAMEMNON, a novel database for Arabidopsis integral membrane proteins. Plant Physiol, 131, 16–26.

Selinski, J. and Scheibe, R. (2014) Pollen tube growth: where does the energy come from? Plant Signal Behav, 9, e977200.

Shi, J., Cui, M., Yang, L., Kim, Y.J. and Zhang, D. (2015) Genetic and Biochemical Mechanisms of Pollen Wall Development. Trends Plant Sci, 20, 741–753.

Somerville, C. and Browse, J. (1991) Plant lipids: metabolism, mutants, and membranes. Science, 252, 80–87.

Stengel, A., Benz, P., Balsera, M., Soll, J. and Bolter, B. (2008) TIC62 redox-regulated translocon composition and dynamics. J Biol Chem, 283, 6656–6667.

Takabayashi, A., Takabayashi, S., Takahashi, K., Watanabe, M., Uchida, H., Murakami, A., Fujita, T., Ikeuchi, M. and Tanaka, A. (2017) PCoM-DB Update: A Protein Co-Migration Database for Photosynthetic Organisms. Plant Cell Physiol, 58, e10.

Takemura, T., Imamura, S. and Tanaka, K. (2019) Identification of a chloroplast fatty acid exporter protein, CmFAX1, and triacylglycerol accumulation by its overexpression in the unicellular red alga Cyanidioschyzon merolae. Algal Research, 38, 101396.

Teese, M.G. and Langosch, D. (2015) Role of GxxxG Motifs in Transmembrane Domain Interactions. Biochemistry, 54, 5125–5135.

Tian, Y., Lv, X., Xie, G., Wang, L., Dai, T., Qin, X., Chen, F. and Xu, Y. (2019) FAX2 Mediates Fatty Acid Export from Plastids in Developing Arabidopsis Seeds. Plant Cell Physiol, 60, 2231–2242.

Tian, Y., Lv, X., Xie, G., Zhang, J., Xu, Y. and Chen, F. (2018) Seed-specific overexpression of AtFAX1 increases seed oil content in Arabidopsis. Biochem Biophys Res Commun, 500, 370–375.

Troncoso-Ponce, M.A., Nikovics, K., Marchive, C., Lepiniec, L. and Baud, S. (2015) New insights on the organization and regulation of the fatty acid biosynthetic network in the model higher plant Arabidopsis thaliana. Biochimie, 120, 3–8.

Voinnet, O., Rivas, S., Mestre, P. and Baulcombe, D. (2003) An enhanced transient expression system in plants based on suppression of gene silencing by the p19 protein of tomato bushy stunt virus. Plant J, 33, 949–956.

Voith von Voithenberg, L., Park, J., Stube, R., Lux, C., Lee, Y. and Philippar, K. (2019) A Novel Prokaryote-Type ECF/ABC Transporter Module in Chloroplast Metal Homeostasis. Front Plant Sci, 10, 1264.

von Berlepsch, S., Kunz, H.H., Brodesser, S., Fink, P., Marin, K., Flugge, U.I. and Gierth, M. (2012) The Acyl-Acyl Carrier Protein Synthetase from Synechocystis sp PCC 6803 Mediates Fatty Acid Import. Plant Physiol, 159, 606-+.

Waegemann, K., Eichacker, S. and Soll, J. (1992) Outer Envelope Membranes from Chloroplasts Are Isolated as Right-Side-out Vesicles. Planta, 187, 89–94.

Wan, X., Wu, S., Li, Z., An, X. and Tian, Y. (2020) Lipid Metabolism: Critical Roles in Male Fertility and Other Aspects of Reproductive Development in Plants. Mol Plant, 13, 955–983.

Warnes, G.R., Bolker, B., Bonebakker, L., Gentleman, R., Huber, W., Liaw, A., Lumley, T., Maechler, M., Magnusson, A., Moeller, S., Schwartz, M., Venables, B. and Galili, T. (2020) Various R Programming Tools for Plotting Data. Maintainer: Tal Galil, Repository: CRAN, URL https://github.com/talgalili/gplots.

Xiao, Z., Tang, F., Zhang, L., Li, S., Wang, S., Huo, Q., Yang, B., Zhang, C., Wang, D., Li, Q., Wei, L., Guo, T., Qu, C., Lu, K., Zhang, Y., Guo, L., Li, J. and Li, N. (2021) The Brassica napus fatty acid exporter FAX1-1 contributes to biological yield, seed oil content, and oil quality. Biotechnol Biofuels, 14, 190.

Xu, C. and Shanklin, J. (2016) Triacylglycerol Metabolism, Function, and Accumulation in Plant Vegetative Tissues. Annual review of plant biology, 67, 179–206.

Zhan, H., Xiong, H., Wang, S. and Yang, Z.N. (2018) Anther Endothecium-Derived Very-Long-Chain Fatty Acids Facilitate Pollen Hydration in Arabidopsis. Mol Plant, 11, 1101–1104.

Zhao, X., Li, N., Song, Q., Li, X., Meng, H. and Luo, K. (2021) OPDAT1, a plastid envelope protein involved in 12-oxo-phytodienoic acid export for jasmonic acid biosynthesis in Populus. Tree Physiol, 41, 1714–1728.

Zhu, L., He, S., Liu, Y., Shi, J. and Xu, J. (2020) Arabidopsis FAX1 mediated fatty acid export is required for the transcriptional regulation of anther development and pollen wall formation. Plant Mol Biol, 104, 187–201.

Zou, Z.Y., Tong, F.M., Faergeman, N.J., Borsting, C., Black, P.N. and DiRusso, C.C. (2003) Vectorial acylation in Saccharomyces cerevisiae - Fat1p and fatty acyl-CoA synthetase are interacting components of a fatty acid import complex. J Biol Chem, 278, 16414–16422.

