## Supplemental Tables, Legends for "Plastid fatty acid export (FAX) proteins in *Arabidopsis thaliana* - the role of FAX1 and FAX3 in growth and development"

**Supplementary data**

**Table S1.** *FAX proteins in Arabidopsis and pea.*

| name | AGI code | protein id | pre [aa] | cTP [aa] | mat [kDa] | loc score | exp |
|---|---|---|---|---|---|---|---|
| **Arabidopsis** | | | | | | | |
| FAX1 | At3g57280 | NP_567046 | 226 | 33 | 19.9 | 25.7 (c) | IE[1,2,3] |
| FAX2* | At2g38550 | NP_565892 | 335 | 72 | 28.6 | 24.8 (c) | IE[2],env[4,5] |
| FAX3* | At3g43520 | NP_566866 | 240 | 84 | 15.9 | 24.1 (c) | IE[2,3] |
| FAX4 | At1g33265 | NP_564422 | 177 | 63 | 18.4 | 20.6 (c) | env[4] |
| FAX5 | At1g50740 | NP_564579 | 119 | | 12.5 | 25.7 (sp) | [6] |
| FAX6 | At3g20510 | NP_188687 | 119 | | 12.5 | 25.7 (sp) | - |
| FAX7 | At2g26240 | NP_180192 | 108 | | 11.1 | 8.6 (sp)/7.3 (m) | [7] |
| **Pea** | **GenBank no.** | | | | | | |
| FAX1 | KF981436 | AIX87540 | 232 | 39 | 20.6 | 0.845 (c) | IE[1] |
| FAX2 | MN737494 | QNQ79270 | 306 | 54 | 27.0 | 0.896 (c) | IE[1] |
| FAX3 | MN737495 | QNQ79271 | 226 | 56 | 16.6 | 0.917 (c) | IE[1] |
| FAX5/6** | MN737496 | QNQ79272 | 120 | | 12.4 | 0.971 (o) | - |

Protein names, *Arabidopsis* genome initiative (AGI) codes or GenBank accession numbers for FAX orthologs in pea as well as protein ids (GenPept) are listed. The coding regions for Ps-FAX1 (Li et al., 2015) as well as Ps-FAX2 and Ps-FAX3 (this study) were amplified by RT-PCR from pea seedling RNA (see Supplementary Fig. S9). The sequence of Ps-FAX5/6 was deduced from pea EST dataset (Franssen et al., 2011). For each protein, the amino acid (aa) count of the full preprotein (pre) and of the predicted N-terminal chloroplast targeting peptide (cTP) as well as the calculated molecular weight in kDa of the corresponding mature protein sequence (mat) are indicated. Experimental evidence (exp) for a plastid inner envelope (IE) insertion of FAX1, FAX2 and FAX3 in this study and the respective literature ([1]: Li et al. [2015]; [2]: Ferro et al. [2010]; [3]: Bouchnak et al. [2019]), as well as envelope (env) localization of At-FAX2 ([4]: Li et al. [2020]; [5]: Tian et al. [2019]), and of At-FAX4 ([4]: Li et al. [2020]) are listed. [6]: Please note that only recently localization of the At-FAX5 ortholog in ER membranes was shown in *Chlamydomonas reinhardtii* (Peter et al., 2022). [7]:Although At-FAX7 was identified in two proteomic analyses of *Arabidopsis* mitochondria membranes (Brugiere et al., 2004, Duncan et al., 2011), upon additional GFP-targeting assays by Duncan and coworkers, At-FAX7 was annotated not to be in mitochondria (Duncan et al., 2011). cTPs were predicted with ChloroP (Emanuelsson et al., 1999). The score for localization prediction (loc score) in *Arabidopsis* was obtained from Aramemnon (AramLocCon; Schwacke et al. [2003]), for *Pisum sativum* (pea) proteins from TargetP (Almagro Armenteros et al., 2019). c, predicted chloroplast; m, predicted mitochondria; sp, predicted secretory pathway localization. * Please note that in contrast to NCBI and UniProt annotations, FAX2 and FAX3 relatives are as originally defined by (Li et al., 2015, Tian et al., 2019, Li et al., 2020) and as named in the Aramemnon database as well as in the current manuscript. ** In pea only one protein similar to At-FAX5 and At-FAX6 (64% and 66% aa identity, respectively) was identified. For Ps-FAX5/6, TargetP predicts no signal peptide (o).

**Table S2.** List of oligonucleotides used in this study.

| name | sequence (5'-3') | application |
|---|---|---|
| fax1_LP | TTTCTTCGCAACATTTTGACC | PCR-genotyping of T-DNA insertion lines |
| fax1_RP | AGTGGAGACACTATCAATCCC | |
| LB1 SAIL | GCCTTTTCAGAAATGGATAAATAGCCTTGCTTCC | |
| GABI-LBseq | ATATTGACCATCATACTCATTGC | |
| fax2_LP | AAACCCTAATTCCACCTTG | |
| fax2_RP | GTATGCCTCCGATGACA | |
| Tag5 IJPB | CTACAAATTGCCTTTTCTTATCGAC | |
| fax3_LP | CAATTCAGAACACTTCCGAACC | |
| fax3_RP | GGAACTCAAAATCAAATCCGC | |
| fax4_LP | GAGTTTCGTTGTATAGGAGATTCTGC | |
| fax4_RP | GAAGACTAAACGAGGTAATGGTCTCTG | |
| AtAct2/8_fw | GGTGATGGTGTGTCT | PCR-genotyping qRT-PCR, RT-PCR |
| AtAct2/8_rev | ACTGAGCACAATGTTAC | |
| AtAct2fw | CTCCTGAAGAGCACCC | |
| AtAct2rev | TCAGTAAGGTCACGTCCA | |
| FAX1 LCfw | CCTATGATTCGTCCCCAG | qRT-PCR, RT-PCR |
| FAX1 LCrev | CACTCACAACGAGACCA | |
| FAX2 LC E2 fw | AAGCGAAAGTGGTTAGTGA | |
| FAX2 LC E5 rev | AGGAACGTCGATTTCTG | |
| FAX3 LC-CT fw | TTCTCGTCGGAGTTGG | |
| FAX3 LC-CT rev | GAGTGTTTGCTACTGCTT | |
| FAX4 LC-CT fw | TAGGAGGCGGTCTCTT | |
| FAX4 LC-CT rev | CAACCTCATGCTCTCATC | |
| FAX4 LC-NT fw | TCTATAACCCAATCCAAACCG | |
| FAX4 LC-NT rev | GCCTCCTAGAAGAAGTACAC | |
| Atfax4 fw (cacc) | CACCATGTGGTCTCTTGCGCTAACGTTAC | |
| Atfax4 rev (-stop) | AGGTAGAGGTAAAGGGTCGGGTATGG | |
| FAX5LCfw | GGAAGTATTGCATCTCTCG | |
| FAX5LCrev | ATCCATATACCTTATTTCAACAGT | |
| FAX6LCfw | GGGATGCTTCTCATCGG | |
| FAX6LCrev | CCACCAGTAGCGATCT | |
| FAX7LCfw | TCATCTTTGTCGCAGAAGT | |
| FAX7LCrev | CTTGAGCGAATGAGGC | |
| Ps-FAX2 fw0 | ATTCACATTACTTCATTTCATAAACCCTTC | isolation of Ps-FAX2, Ps-FAX3 cDNA |
| Ps-FAX2-rev | GGTTCAAACTTGGAAGGTGAAGCAGGA | |
| Ps-FAX3 fw | ATGGCGGTTTTAGGCCTTTCG | |
| Ps-FAX3 rev | CGGAATTATGCGTAGTAGTTCGCACTAG | |

**Fig. S1.** Definition of the FAX protein family in *Arabidopsis*.

(A) Sequence alignment of the FAX protein family in *Arabidopsis*. Mature protein sequences of the plastid localized FAX1, FAX2, FAX3, FAX4 are according to ChloroP and TargetP 2.0 predictions of processing sites for stromal peptidases (Emanuelsson et al., 1999, Almagro Armenteros et al., 2019). The four membrane-embedded α-helices (orange boxes) within the Tmemb_14 domain (pfam PF03647 motif, green line) of each FAX protein are depicted according to the Aramemnon consensus prediction AramTmCon (Schwacke et al., 2003). Names, *Arabidopsis* genome initiative codes (AGI) and protein ids (GenPept) for *Arabidopsis* FAX proteins are as follows: At-FAX1 (At3g57280, NP_567046.1), At-FAX2 (At2g38550, NP_565892.1), At-FAX3 (At3g43520, NP_566866.1), At-FAX4 (At1g33265, NP_564422.1), At-FAX5 (At1g50740, NP_564579.1), At-FAX6 (At3g20510, NP_188687.1), and At-FAX7 (At2g26240, NP_180192.1). Please note that in NCBI and UniProt databases, plant FAX2 proteins in contrast to the definition in all original publications, i.e. in Li et al. [2015], Tian et al. [2019], Li et al. [2020], are annotated as FAX3 and *vice versa*. In the Aramemnon plant membrane protein database (Schwacke et al., 2003), in the publications mentioned above and in the current manuscript, however, FAX2 and FAX3 proteins correspond to the AGI codes and protein ids as defined above.

(B) Amino acid (aa) identities in % of mature proteins shown in (A). Values in green are for aa identities within the Tmemb_14 domain only. Please note that the Tmemb_14 domain of FAX1 is most similar to that of FAX5 (36%), FAX6 (38%), and FAX2 (34%), while that of FAX3 strongly resembles FAX7 (56%). Sequences of FAX5 and FAX6 are 81% identical.

**Fig. S2.** FAX1 proteins in different plant species.

(A) Mature protein sequences are depicted according to ChloroP and TargetP 2.0 predictions of processing sites for chloroplast stromal peptidases (Emanuelsson et al., 1999, Almagro Armenteros et al., 2019), for At-FAX1 and Cr-FAX1 see also Peter et al. (2022). The four membrane-embedded α-helices (H1-H4, blue boxes) within the Tmemb_14 domain (pfam PF03647 motif, green line) are depicted according to the Aramemnon consensus prediction AramTmCon (Schwacke et al., 2003) to the sequence of At-FAX1. Conserved motifs, positively (R, K, H; red dots) and negatively (D, E; blue dots) charged amino acid residues (see Fig. 1) as well as the predicted N-terminal, non-hydrophobic α-helix of At-FAX1 (h0, compare Fig. 2A) are indicated. Names in the alignment, different organisms, accession numbers of GenPept entries, and gene numbers (GeneBank, Aramemnon) are as follows: At (*Arabidopsis thaliana*, NP_567046, At3g57280), Br386 (*Brassica rapa*, VDC99386, LR031574), Br405 (*Brassica rapa*, RID54405, CM010634), Br859 (*Brassica rapa*, VDC62859, LR031568), Sl (*Solanum lycopersicum*, NP_001307153, NM_001320224), Mt (*Medicago truncatula,* XP_003625097, XM_003625049), Ps (*Pisum sativum*, AIX87540, KF981436), Bv (*Beta vulgaris*, XP_010672925, XM_010674623), Gm (*Glycine max*, XP_003554154, XM_003554106), Os450 (*Oryza sativa*, XP_015635421, LOC_Os04g39450), Os690 (*Oryza sativa*, BAS97383, LOC_Os06g0301100), Zm081 (*Zea mays* B73, XP_008652081, GRMZM2G046529), Zm990 (*Zea mays* Mo17 inbred line, PWZ45990, GRMZM2G173649), Pp022 (*Physcomitrium patens*, XP_024359022, XM_024503254), Pp899 (*Physcomitrium patens*, XP_024401899, XM_024546131), Pp988 (*Physcomitrium patens*, XP_024403988, XM_024548220), Cr (*Chlamydomonas reinhardtii*, XP_001702596, Cre10.g421750 [Phytozome]; Peter et al. [2022]). Please note that due to uncertain prediction of a chloroplast targeting peptide (cTP), the N-terminal sequence of the Zm081 preprotein has been manually shortened fit to the size of all other FAX1 proteins. Further, all three FAX1-like proteins from *Physcomitrium patens* contain an unusually long FAX2-like apolipoprotein N-terminus (compare Supplementary Fig. S8), which here has been deleted to fit to the alignment.

(B) Amino acid (aa) identities in % of proteins shown in (A). Values in green are for aa identities within the Tmemb_14 domain only.

**Fig. S3.** FAX3 proteins in different plant species.

(A) Mature protein sequences are depicted according to ChloroP and TargetP 2.0 predictions of processing sites for chloroplast stromal peptidases (Emanuelsson et al., 1999, Almagro Armenteros et al., 2019). The four membrane- embedded α-helices (H1-H4, blue boxes) within the Tmemb_14 domain (pfam PF03647 motif, green line) are depicted according to the Aramemnon consensus prediction AramTmCon (Schwacke et al., 2003) to the sequence of At-FAX3. Conserved positively (R, K, H; red dots) and negatively (D, E; blue dots) charged amino acid residues (see Fig. 1) are indicated. The N-terminal poly glycine stretch (poly G, compare Fig. 2E) extends from the conserved proline residue in motif 4 (see [C]) until helix 1. Please note that in NCBI and UniProt databases, plant FAX3 proteins in contrast to the definition in the original publication, i.e. in Li et al. (2015), are annotated as FAX2 and *vice versa*. In the Aramemnon plant membrane protein database (Schwacke et al., 2003), in Li et al. (2015), and in the current manuscript, however, all FAX3 relatives are similar to the protein with the *Arabidopsis* At3g43520 (At-FAX3). Names in the alignment, different organisms, accession numbers of GenPept entries, and gene numbers (GeneBank, Aramemnon) are as follows: At (*Arabidopsis thaliana*, NP_566866, At3g43520), Br (*Brassica rapa*, RID58705, CM010633), Sl (*Solanum lycopersicum*, XP_004251316, XM_004251268), Mt (*Medicago truncatula,* XP_013458374, XM_013602920), Ps (*Pisum sativum*, MN737495), Gm (*Glycine max*, XP_003542391, XM_003542343), Os (*Oryza sativa*, XP_015635691, LOC_Os04g55930), Zm (*Zea mays*, NP_001136902, GRMZM2G386525), Pp701 (*Physcomitrium patens*, XP_024362701, XM_024506933), Pp764 (*Physcomitrium patens*, XP_024362764, XM_024506996), Cr (*Chlamydomonas reinhardtii*, Cre08.g383300 [Phytozome]; Peter et al. [2022]). Please note that the sequence for Pp701 and Pp764 proteins was shortened at the N-terminus to the length of the mature At-FAX3 sequence.

(B) Amino acid (aa) identities in % of proteins shown in (A). Values in green are for aa identities within the Tmemb_14 domain only.

(C) Motif consensus prediction of the mature FAX3 protein sequences shown in (A) according to MEME software (Bailey et al., 2015). Please note that the consensus motifs 4 (blue), 5 (orange), and 3 (green) are in the N-terminal poly glycin stretches (compare Fig. 2E), whereas motifs 1 (red) and 2 (turquoise) are located in the Tmemb_14 domain.

**Fig. S4.** FAX2 proteins in different plant species.

(A) Mature protein sequences are depicted according to ChloroP and TargetP 2.0 predictions of processing sites for chloroplast stromal peptidases (Emanuelsson et al., 1999, Almagro Armenteros et al., 2019). The four membrane- embedded α-helices (H1-H4, blue boxes) within the Tmemb_14 domain (pfam PF03647 motif, green line) are depicted according to the Aramemnon consensus prediction AramTmCon (Schwacke et al., 2003) to the sequence of At-FAX2. Conserved positively (R, K, H; red dots) and negatively (D, E; blue dots) charged amino acid residues (see Fig. 1) as well as the predicted N-terminal, apolipoprotein domain of At-FAX2 (helices ho-h3, orange boxes; compare Fig. 2) are indicated. Please note that in NCBI and UniProt databases, plant FAX2 proteins in contrast to the definition in all original publications, i.e. in Li et al. (2015), Tian et al. (2019), Li et al. (2020), are annotated as FAX3 and *vice versa*. In the Aramemnon plant membrane protein database (Schwacke et al., 2003), in the publications mentioned above and in the current manuscript, however, all FAX2 relatives are similar to the protein with the *Arabidopsis* At2g38550 (At-FAX2). Names in the alignment, different organisms, accession numbers of GenPept entries, and gene numbers (GeneBank, Aramemnon) are as follows: At (*Arabidopsis thaliana*, NP_565892, At2g38550), Br (*Brassica rapa*, XP_009143375, XM_009145127), Sl (*Solanum lycopersicum*, XP_004246287, XM_004246239), Mt (*Medicago truncatula,* XP_003623630, XM_003623582), Ps (*Pisum sativum*, MN737494), Bv (*Beta vulgaris*, XP_010683481, XM_010685179), Gm (*Glycine max*, XP_003552314, XM_003552266), Os (*Oryza sativa*, XP_015621232, Os01g60650), Zm377 (*Zea mays*, NP_001132377, GRMZM2G059353), Zm146 (*Zea mays*, ACG49146, GRMZM2G166441).

(B) Amino acid (aa) identities in % of proteins shown in (A). Values in green are for aa identities within the Tmemb_14 domain only.

**Fig. S5.** FAX4 proteins in different plant species.

(A) Mature protein sequences are depicted according to TargetP 2.0 predictions of processing sites for chloroplast stromal peptidases (Almagro Armenteros et al., 2019). The four membrane-embedded α-helices (H1-H4, blue boxes) within the Tmemb_14 domain (pfam PF03647 motif, green line) are depicted according to the Aramemnon consensus prediction AramTmCon (Schwacke et al., 2003) to the sequence of At-FAX4. Conserved positively (R, K, H; red dots) and negatively (D, E; blue dots) charged amino acid residues (see Fig. 1) are indicated. Names in the alignment, different organisms, accession numbers of UniProt/GenPept entries are as follows: At (*Arabidopsis thaliana*, Q8LPG1, At1g33265), Br (*Brassica rapa*, RID45910), Bn (*Brassica napus*, A0A078J2E9), Dc (*Daucus carota*, A0A175YNA6), Ha (*Helianthus annuus,* A0A251RPY1), Vv (*Vitis vinifera*, F6HUB3), Pt (*Populus trichocarpa*, A0A3N7FVE9), Py (*Prunus yedoensis*, A0A314XXC1), Ma (*Musa acuminata*, M0SFM6), Os (*Oryza sativa*, Q01KJ0), Zm (*Zea mays* B6TKU2), Sm (*Selaginella moellendorffii*, XP_002970789.2), Pp (*Physcomitrium patens*, A0A2K1KQF7). Please note that in contrast to FAX1, FAX2, FAX3 (compare Supplementary Figs. S2, S3, S4), we were unable to identify FAX4 proteins in *Solanum lycopersicon, Medicago truncatula, Pisum sativum, Beta vulgaris and Glycine max.*

(B) Amino acid (aa) identities in % of proteins shown in (A).

**Fig. S6.** Helical wheel projections of the four predicted α-helices in the Tmemb_14 domain of FAX-family proteins.

Helical wheels of α-helices H1-H4 of FAX1 (A), FAX2 (B), FAX3 (C), FAX4 (D), FAX5/6 (E), and FAX7 (G). Helical wheels were produced by NetWheels software (Mól et al. [2018]; settings: α-helix with 3.6 amino acid residues per helix turn) by using the respective consensus sequence for all plant FAX proteins, which is given below each helix projection (compare Supplementary Figs. S1-S5, S7, Könnel et al. [2019]). In case of no distinct consensus, the amino acid of the respective *Arabidopsis* FAX protein sequence was used (green letters). Positively charged (red), negatively charged (blue), polar/uncharged (green), and nonpolar/hydrophobic (yellow) amino acid residues are depicted. Blue dashed lines indicate the separation of hydrophobic and polar helix surface regions along the entire helix cylinder of H3 in FAX1, FAX2, FAX4, FAX5/6 and of H4 in FAX7 (compare Fig. 1).

**Fig. S7.** FAX7 proteins in different plant species.

(A) The four membrane-embedded α-helices (H1-H4, blue boxes) within the Tmemb_14 domain (pfam PF03647 motif, green line) are depicted according to the Aramemnon consensus prediction AramTmCon (Schwacke et al., 2003) to the sequence of At-FAX7. Conserved positively (R, K, H; red dots) and negatively (D, E; blue dots) charged amino acid residues (see Fig. 1) are indicated. Names in the alignment, different organisms, accession numbers of UniProt/GenPept entries are as follows: At (*Arabidopsis thaliana*, O64847, At2g26240), Br (*Brassica rapa*, XP_009117028.1), Bn (*Brassica napus*, A0A078FEW8, +22: please note that the last 22 aa predicted for Bn-FAX7 are not depicted), So (*Spinacia oleracea*, A0A0K9R288), Ma (*Musa acuminata*, M0T184), Ob (*Oryza brachyantha*, J3M249), Zm (*Zea mays* PWZ43434.1, asterisks indicate incomplete open reading frame), Sm-330 (*Selaginella moellendorffii*, D8R330), Sm-G47 (*Selaginella moellendorffii*, D8SG47), Cr (*Chlamydomonas reinhardtii*, Cre09.g387838 [Phytozome]; Peter et al. [2022]).

(B) Amino acid (aa) identities in % of proteins shown in (A).

**Fig. S8.** FAX1-like proteins with a FAX2-like N-terminal apolipoprotein domain in moss and spikemoss.

Mature protein sequences are depicted according to TargetP 2.0 (Almagro Armenteros et al., 2019) predictions of processing sites for chloroplast stromal peptidases in *Physcomitrium patens* (Pp) proteins. Please note that probably due to still incomplete genome annotation the sequences of the *Selaginella moellendorfii* (Sm) proteins are only partial and appear to be truncated at the N- and C-Terminus. The Tmemb_14 domain (pfam PF03647 motif) is depicted as green line. Conserved FAX1-like motifs in Tmemb14 (see Fig. 1, Supplementary Fig. S2) are indicated. Interestingly, the FAX1-like proteins with the characteristic motifs in the C-terminal Tmemb14 domain, in *Physcomitrium* and *Selaginella*, contain an extended N-terminus with a FAX2-like apolipoprotein domain. The secondary structure of the Pp022 N-terminus (modelled at 92% confidence) was predicted by Phyre$^2$ (Kelley et al., 2015) in comparison to the NMR structure of a helix bundle of an insect apolipophorin-III (template d1eq1a: *Manduca sexta* apoliophorin-III, PDB entry 1EQ1). The expected Pp022 apolipoprotein domain is indicated.

Names in the alignment, organisms, accession numbers of GenPept entries, and gene numbers are as follows (for Pp proteins, see also Supplementary Fig. S2): Pp022 (*Physcomitrium patens*, XP_024359022, XM_024503254), Pp899 (*Physcomitrium patens*, XP_024401899, XM_024546131), Pp988 (*Physcomitrium patens*, XP_024403988, XM_024548220), Sm654 (*Selaginella moellendorfii*, EFJ34654.1, GL377569.1), Sm059 (*Selaginella moellendorfii*, EFJ31059.1, GL377574.1).

**Fig. S9.** FAX1, FAX2, FAX3 proteins in *Arabidopsis* and pea.

Amino acid sequences of mature At-FAX1, Ps- FAX1, At-FAX2, Ps-FAX2, At-FAX3, and Ps-FAX3 proteins. The coding regions for Ps-FAX1 (Li et al., 2015) as well as Ps-FAX2 and Ps-FAX3 (this study) were amplified by RT-PCR from pea seedling RNA. For GenBank and GenPept accession numbers see Table S1. Amino acids (aa) conserved in all sequences are shaded in black, those found in 60% of the proteins are highlighted in dark grey, those in 30% in light grey. Sequence identity between *Arabidopsis* and pea FAX1, FAX2, and FAX3 is 52%, 49%, and 56%, respectively. The four membrane-embedded α-helices of the Tmemb_14 domain (light blue boxes) are depicted according to Aramemnon consensus prediction AramTmCon (Schwacke et al., 2003) of *Arabidopsis* proteins. Presumed thermolysin cleavage sites pointing to the intermembrane space (compare Fig. 4) are indicated by orange triangles. Oligopeptides used to generate antisera against Ps-FAX1 in Li et al. (2015) as well as Ps-FAX2 and Ps-FAX3 (this study) are specified by yellow boxes. The most N-terminal peptides, identified after trypsin proteolysis and peptide sequencing of the respective bands that were stained by antisera in SDS gels are highlighted by magenta boxes. For the latter analysis, tryptic digest of PAGE separated proteins and mass spectrometric analysis was essentially performed by Frederik Sommer (TU Kaiserslautern, Germany) as described in (Peter et al., 2022). MS data were searched against proteins from *Arabidopsis thaliana* (for At-FAX1) and *Pisum sativum* derived from UniProt database, supplemented with the coding sequences of Ps-FAX1-3 described in Li et al. (2015) for Ps-FAX1 and in this study for Ps-FAX2 and 3.

**Fig. S10.** Characterization of T-DNA insertion lines in *At-FAX2*, *At-FAX3*, and *At-FAX4*.

(A)-(C) Left, schematic representations of the genomic organization of *At-FAX2* (A), *At-FAX3* (B) and *At-FAX4* (C) according to the Aramemnon database (Schwacke et al., 2003). Black arrows indicate coding sequences of exons, white bars represent introns and untranslated regions (UTR). Please note that the lengths of the respective regions are drawn to scale, except 5'-UTR regions, which have been shortened from the 5'-end. The adenosine nucleotide of the translational start codon ("ATG") is designated as "+1". T-DNA insertion sites are indicated by triangles. Orientation and binding sites of left-border, T-DNA-specific, and of gene-specific oligonucleotide primers used for PCR-genotyping are depicted.

(A) For the *FAX2* gene a total length of 4300 bp (1827 bp 5'UTR, 2052 bp from translational start to stop codon, 421 bp 3'UTR) and six exons are annotated. In the T-DNA insertion line *fax2-1* (FLAG_426D07), the T-DNA has inserted into the first exon at position +233. Please note that *fax2-1* was generated in Wassilewskija (Ws) ecotype and is different from the T-DNA insertion lines in the 5'UTR of *FAX2* in Col-0 background described in Tian et al. (2019), Li et al. (2020). As reported by Tian and coworkers (Tian et al., 2019), but in contrast to Li et al. (2020), we detected wild-type like levels of *FAX2* transcripts in T-DNA insertion lines in the 5'UTR of *FAX2* (not shown).

(B) The *FAX3* gene is 3850 bp long (1837 bp 5'UTR, 1612 bp from translational start to stop codon, 401 bp 3'UTR) and contains three exons. The T-DNA in insertion line *fax3-1* (FLAG_454B11, in Ws ecotype) has inserted into the 5′-UTR at position -23.

(C) The *FAX4* gene is 3400 bp long (1812 bp 5'UTR, 1158 bp from translational start to stop codon, 430 bp 3'UTR) and includes four exons. For *FAX4* three different T-DNA insertion lines were analyzed. In *fax4-1* (SAIL788B10) and *fax4-2* (SAIL1176C06), the T-DNAs have inserted into the 5'-UTR (*fax4-1*, position -64; *fax4-2,* position -11). In the line *fax4-3* (GK-930E07), the T-DNA insertion site is in the first exon at position +118. Please note that the T-DNA insertion line in the 5'UTR of *FAX4* described in Li et al. (2020), i.e. SAIL671D03, is different from the lines *fax4-1*, *fax4-2, fax4-3* characterized in our study.

(A)-(C) Right, PCR-genotyping (gDNA) and RT-PCR (cDNA) analyses of the respective T-DNA mutant lines. A PCR product of actin (435bp in [A, B]; 271bp in [C]) was used as control. <u>gDNA:</u> PCR-genotyping results of *fax2-1* (A), *fax3-1* (B), and *fax4-1, fax4-2, fax4-3* (C) mutant lines. *At-FAX2, At-FAX3,* and *At-FAX4* gene-specific primers in combination with T-DNA-specific left border primers generated fragments of 466 bp (*fax2-1*, [A]), 536 bp (*fax3-1*, [B]), as well as 365 bp, 664 bp, and 567 bp (*fax4-1*, *fax4-2*, *fax4-3*, respectively, [C]) on genomic DNA of heterozygous (he) and homozygous (ho) plants. For verification of PCR products and

T-DNA insertion sites, amplified DNA fragments were sequenced. To identify plants with T-DNA in both alleles of the *FAX* genes, we used gene-specific primers flanking the predicted insertion sites. gDNA from homozygous mutations gave no amplification product, whereas the amplified regions on wild-type (wt) and he DNA were 595 bp (*fax2-1* [A]), 335 bp (*fax3-1* [B]), and 601 bp, (for *fax4-1*, *fax4-2*, *fax4-3* [C]).

<u>cDNA:</u> RT-PCR on cDNA from segregated homozygous (ho) and wild-type plants (wt) of the mutants described above and the *fax1-2/fax3-1* double mutant (*fax1/3* dm). RNA was prepared from the respective plants and reverse transcribed into cDNA as described (Duy et al., 2007). Primer pairs specific for *At-FAX2* (A), *At-FAX3* (B) and *At-FAX4* (C) only in wild-type plants amplified products of 388 bp (*fax2-1* [A]), 303 bp (*fax3-1* [B]), and 321 bp, (*fax4-3* [C]). Thus, *fax2-1*, *fax3-1*, and *fax4-3* homozygous lines are knockouts without RNA of the respective genes. Please note that for *fax4-1*, *fax4-2* in (C), ho plants contained RNA (PCR product of 321 bp on cDNA), indicating that the T-DNA insertion in the 5'UTR of these lines does not lead to the loss of *FAX4* transcripts. Further, the *fax1/3* dm line is homozygous for *fax3-1* but heterozygous for the *fax1-2* allele (product of 265 bp on cDNA).

(D) Segregation analysis of *fax2-1*, *fax3-1,* and *fax4-3* mutants. The progeny of heterozygous single *fax2-1*, *fax3-1,* and *fax4-3* mutant lines was germinated on agar medium and analyzed after three weeks by PCR-genotyping as described in (A-C). The genotype of homozygous (ho), heterozygous (he), and wild-type (wt) mutant alleles is given in percent. no.: number of plantlets analyzed.

**Fig. S11.** Expression of *Arabidopsis FAX* genes in rosette leaves of FAX2-, FAX3-, and FAX4 loss-of-function mutants.

The transcript content of *At-FAX1*, *At-FAX2*, *At-FAX3*, *At-FAX4*, *At-FAX5*, *At-FAX6*, and *At-FAX7* was analyzed by quantitative real-time RT-PCR in rosette leaves of 4-week-old *fax2-1* (A), *fax3-1* (B), and *fax4-3* (C) knockouts (black bars) as well as the corresponding segregated wild types (WT2, WT3) and Col-0 (white bars). The mRNA of each *Arabidopsis FAX* gene was quantified relative to that of actin 2/8 and normalized to the respective amount in wild type, which was set to 100 (a. u., arbitrary units). Data points represent mean values from three independent biological replicates ($n = 3 \pm SD$). Please note that for each biological replicate, rosette tissue from three individual plants was pooled prior to RNA extraction. Asterisks indicate significantly different values when compared to wild type (*: $p < 0.05$, **: $p < 0.01$, Student's t-test).

### References

**Almagro Armenteros, J.J., Salvatore, M., Emanuelsson, O., Winther, O., von Heijne, G., Elofsson, A. and Nielsen, H.** (2019) Detecting sequence signals in targeting peptides using deep learning. *Life Science Alliance*, **2**, e201900429.

**Bailey, T.L., Johnson, J., Grant, C.E. and Noble, W.S.** (2015) The MEME Suite. *Nucleic Acids Res*, **43**, 39-49.

**Bouchnak, I., Brugiere, S., Moyet, L., Le Gall, S., Salvi, D., Kuntz, M., Tardif, M. and Rolland, N.** (2019) Unraveling Hidden Components of the Chloroplast Envelope Proteome: Opportunities and Limits of Better MS Sensitivity. *Mol Cell Proteomics*, **18**, 1285-1306.

**Brugiere, S., Kowalski, S., Ferro, M., Seigneurin-Berny, D., Miras, S., Salvi, D., Ravanel, S., d'Herin, P., Garin, J., Bourguignon, J., Joyard, J. and Rolland, N.** (2004) The hydrophobic proteome of mitochondrial membranes from Arabidopsis cell suspensions. *Phytochemistry*, **65**, 1693-1707.

**Duncan, O., Taylor, N.L., Carrie, C., Eubel, H., Kubiszewski-Jakubiak, S., Zhang, B., Narsai, R., Millar, A.H. and Whelan, J.** (2011) Multiple lines of evidence localize signaling, morphology, and lipid biosynthesis machinery to the mitochondrial outer membrane of Arabidopsis. *Plant Physiol*, **157**, 1093-1113.

**Duy, D., Wanner, G., Meda, A.R., von Wiren, N., Soll, J. and Philippar, K.** (2007) PIC1, an ancient permease in Arabidopsis chloroplasts, mediates iron transport. *Plant Cell*, **19**, 986-1006.

**Emanuelsson, O., Nielsen, H. and Von Heijne, G.** (1999) ChloroP, a neural network-based method for predicting chloroplast transit peptides and their cleavage sites. *Protein Sci*, **8**, 978-984.

**Ferro, M., Brugiere, S., Salvi, D., Seigneurin-Berny, D., Court, M., Moyet, L., Ramus, C., Miras, S., Mellal, M., Le Gall, S., Kieffer-Jaquinod, S., Bruley, C., Garin, J., Joyard, J., Masselon, C. and Rolland, N.** (2010) AT_CHLORO, a comprehensive chloroplast proteome database with subplastidial localization and curated information on envelope proteins. *Mol Cell Proteomics*, **9**, 1063-1084.

**Franssen, S.U., Shrestha, R.P., Bräutigam, A., Bornberg-Bauer, E. and Weber, A.P.** (2011) Comprehensive transcriptome analysis of the highly complex Pisum sativum genome using next generation sequencing. *BMC Genomics*, **12**, 227.

**Kelley, L.A., Mezulis, S., Yates, C.M., Wass, M.N. and Sternberg, M.J.** (2015) The Phyre2 web portal for protein modeling, prediction and analysis. *Nature protocols*, **10**, 845-858.

**Könnel, A., Bugaeva, W., Guegel, I.L. and Philippar, K.** (2019) BANFF: bending of bilayer membranes by amphiphilic alpha-helices is necessary for form and function of organelles (1). *Biochem Cell Biol*, **97**, 243-256.

**Li, N., Gügel, I.L., Giavalisco, P., Zeisler, V., Schreiber, L., Soll, J. and Philippar, K.** (2015) FAX1, a novel membrane protein mediating plastid fatty acid export. *PLoS Biology*, **13**, e1002053.

**Li, N., Meng, H., Li, S., Zhang, Z., Zhao, X., Wang, S., Liu, A., Li, Q., Song, Q., Li, X., Guo, L., Li, H., Zuo, J. and Luo, K.** (2020) Two novel plastid fatty acid exporters contribute to seed oil accumulation in Arabidopsis. *Plant Physiol*, **Preview, published on February 4**, doi: 10.1104/pp.1119.01344.

**Mól, A.R., Castro, M.S. and Fontes, W.** (2018) NetWheels: A web application to create high quality peptide helical wheel and net projections. 416347.

**Peter, J., Huleux, M., Spaniol, B., Sommer, F., Neunzig, J., Schroda, M., Li-Beisson, Y. and Philippar, K.** (2022) Fatty acid export (FAX) proteins contribute to oil production in the green microalga Chlamydomonas reinhardtii. *Frontiers in molecular biosciences*, **9**, 939834.

**Schwacke, R., Schneider, A., van der Graaff, E., Fischer, K., Catoni, E., Desimone, M., Frommer, W.B., Flugge, U.I. and Kunze, R.** (2003) ARAMEMNON, a novel database for Arabidopsis integral membrane proteins. *Plant Physiol*, **131**, 16-26.

**Tian, Y., Lv, X., Xie, G., Wang, L., Dai, T., Qin, X., Chen, F. and Xu, Y.** (2019) FAX2 Mediates Fatty Acid Export from Plastids in Developing Arabidopsis Seeds. *Plant Cell Physiol*, **60**, 2231-2242.
