## Supplemental Fig S1-S5,S7-S9 for "Plastid fatty acid export (FAX) proteins in *Arabidopsis thaliana* - the role of FAX1 and FAX3 in growth and development"

**A**

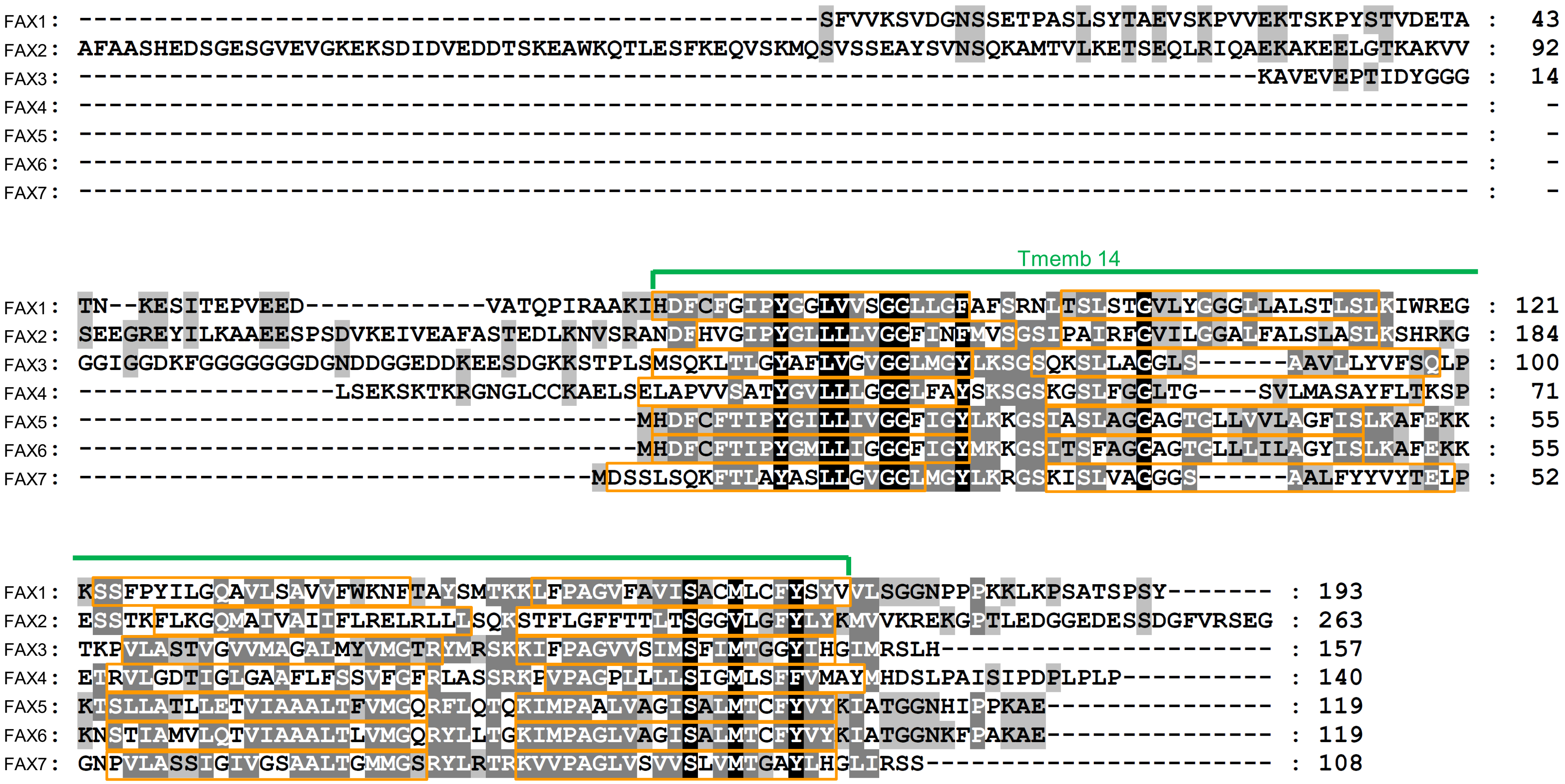
FAX1 : -------------------------------------------------SFVVKSVDGNSSETPASLSYTAEVSKPVVEKTSKPYSTVDETA : 43
FAX2 : AFAASHEDSGESGVEVGKEKSDIDVEDDTSKEAWKQTLESFKEQVSKMQSVSSEAYSVNSQKAMTVLKETSEQLRIQAEKAKEELGTKAKVV : 92
FAX3 : ------------------------------------------------------------------------------KAVEVEPTIDYGGG : 14
FAX4 : -------------------------------------------------------------------------------------------- : -
FAX5 : -------------------------------------------------------------------------------------------- : -
FAX6 : -------------------------------------------------------------------------------------------- : -
FAX7 : -------------------------------------------------------------------------------------------- : -
Tmemb 14
FAX1 : TN--KESITEPVEED------------VATQPIRAAKIHDFCFGIPYGGLVVSGGLLGFAFSRNLTSLSTGVLYGGGLLALSTLSLKIWREG : 121
FAX2 : SEEGREYILKAAEESPSDVKEIVEAFASTEDLKNVSRANDFHVGIPYGLLLLVGGFINFMVSGSIPAIRFGVILGGALFALSLASLKSHRKG : 184
FAX3 : GGIGGDKFGGGGGGGDGNDDGGEDDKEESDGKKSTPLSMSQKLTLGYAFLVGVGGLMGYLKSGSQKSLLAGGLS------AAVILYVFSQLP : 100
FAX4 : -----------------LSEKSKTKRGNGLCCKAELSELAPVVSATYGVLLLGGGLFAYSKSGSKGSLFGGLTG----SVLMASAYFLTKSP : 71
FAX5 : -------------------------------------MHDFCFTIPYGILLIVGGFIGYLKKGSIASLAGGAGTGLLVVLAGFISLKAFEKK : 55
FAX6 : -------------------------------------MHDFCFTIPYGMLLIGGGFIGYMKKGSITSFAGGAGTGLLLILAGYISLKAFEKK : 55
FAX7 : ----------------------------------MDSSLSQKFTLAYASLLGVGGLMGYLKRGSKISLVAGGGS------AALFYVYTELP : 52
FAX1 : KSSFPYILGQAVLSAVVFWKNFTAYSMTKKLFPAGVFAVISACMLCFYSYVVLSGGNPPPKKLKPSATSPSY------- : 193
FAX2 : ESSTKFLKGQMAIVAIIFLRELRLLLSQKSTFLGFFTTLTSGGVLGFYLYKMVVKREKGPTLEDGGEDESSDGFVRSEG : 263
FAX3 : TKPVLASTVGVVMAGALMYVMGTRYMRSKKIFPAGVVSIMSFIMTGGYIHGIMRSLH---------------------- : 157
FAX4 : ETRVLGDTIGLGAAFLFSSVFGFRLASSRKPVPAGPLLLLSIGMLSFFVMAYMHDSLPAISIPDPLPLP---------- : 140
FAX5 : KTSLLATLLETVIAAALTFVMGQRFLQTQKIMPAALVAGISALMTCFYVYKIATGGNHIPPKAE--------------- : 119
FAX6 : KNSTIAMVLQTVIAAALTLVMGQRYLLTGKIMPAGLVAGISALMTCFYVYKIATGGNKFPAKAE--------------- : 119
FAX7 : GNPVLASSIGIVGSAALTGMMGSRYLRTRKVVPAGLVSVVSLVMTGAYLHGLIRSS----------------------- : 108

**B**

| | FAX1 | FAX2 | FAX3 | FAX4 | FAX5 | FAX6 | FAX7 |
|---|---|---|---|---|---|---|---|
| FAX1 | **100%** | | | | | | |
| FAX2 | **20%**<br>34% | **100%** | | | | | |
| FAX3 | **12%**<br>24% | **6%**<br>13% | **100%** | | | | |
| FAX4 | **10%**<br>17% | **8%**<br>19% | **18%**<br>29% | **100%** | | | |
| FAX5 | **22%**<br>36% | **12%**<br>26% | **21%**<br>33% | **18%**<br>25% | **100%** | | |
| FAX6 | **23%**<br>38% | **14%**<br>29% | **20%**<br>31% | **18%**<br>24% | **81%**<br>80% | **100%** | |
| FAX7 | **11%**<br>21% | **6%**<br>14% | **39%**<br>56% | **24%**<br>30% | **33%**<br>37% | **32%**<br>37% | **100%** |

Figure S2

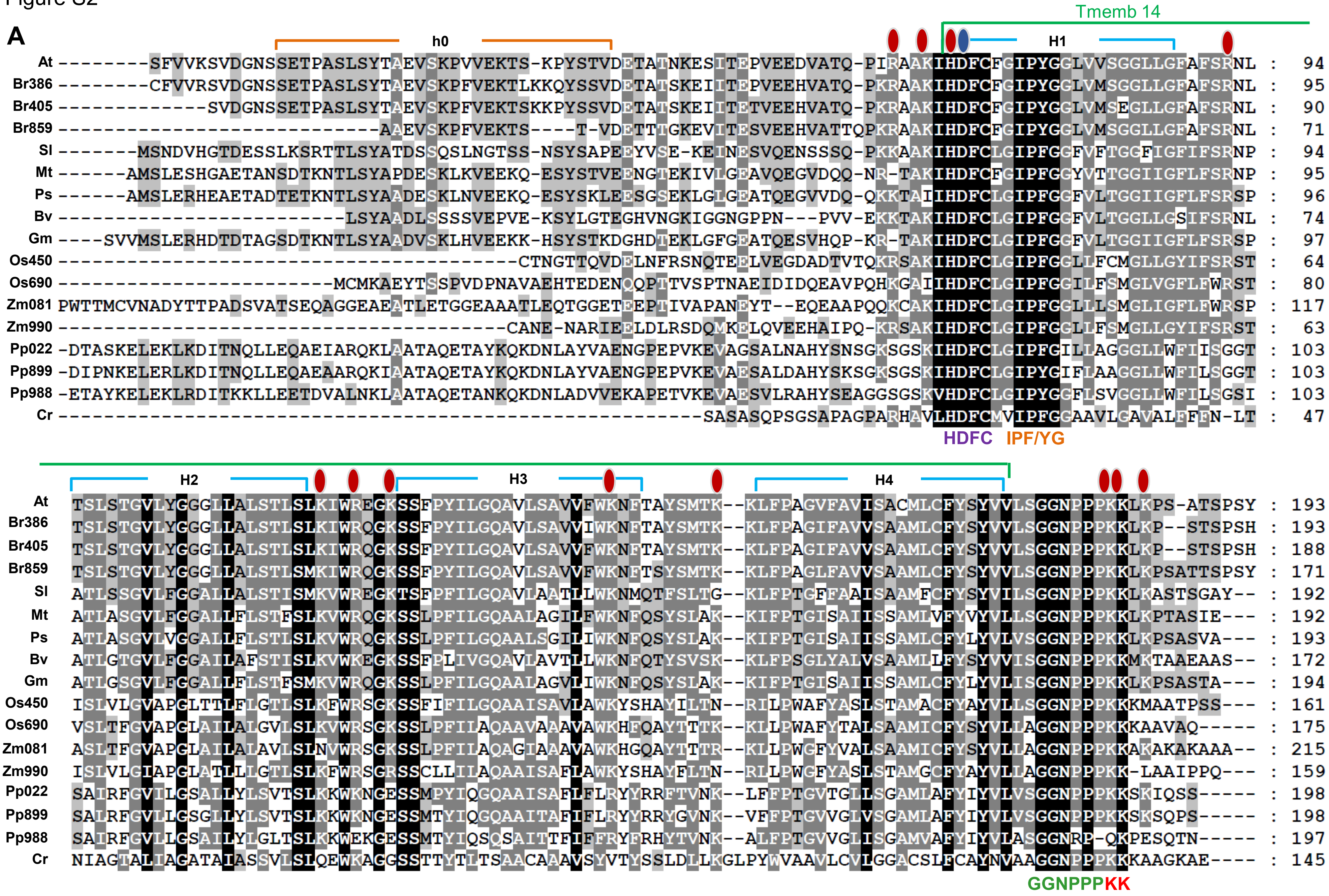
A
h0
Tmemb 14
H1
At --------SFVVKSVDGNSSETPASLSYTAEVSKPVVEKTS-KPYSTVDETATNKESITEPVEEDVATQ-PIRAAKIHDFCFGIPYGGLVVSGGLLGFAFSRNL : 94
Br386 --------CFVVRSVDGNSSETPASLSYTAEVSKPFVEKTLKKQYSSVDETATSKEIITEPVEEHVATQ-PKRAAKIHDFCFGIPYGGLVMSGGLLGFAFSRNL : 95
Br405 -------------SVDGNSSETPASLSYTAEVSKPFVEKTSKKPYSSVDETATSKEIITETVEEHVATQ-PKRAAKIHDFCFGIPYGGLVMSEGLLGFAFSRNL : 90
Br859 ---------------------------AAEVSKPFVEKTS----T-VDETTTGKEVITESVEEHVATTQPKRAAKIHDFCFGIPYGGLVMSGGLLGFAFSRNL : 71
Sl -------MSNDVHGTDESSLKSRTTLSYATDSSQSLNGTSS-NSYSAPEEYVSE-KEINESVQENSSSQ-PKKAAKIHDFCLGIPFGGFVFTGGFIGFIFSRNP : 94
Mt ------AMSLESHGAETANSDTKNTLSYAPDESKLKVEEKQ-ESYSTVEENGTEKIVLGEAVQEGVDQQ-NR-TAKIHDFCFGIPFGGYVTTGGIIGFLFSRNP : 95
Ps ------AMSLERHEAETADTETKNTLSYAADESKLNVEEKQ-ESYSKLEESGSEKLGLGEATQEGVVDQ-QKKTAIIHDFCLGIPFGGFVLTGGIIGFLFSRSP : 96
Bv -------------------------LSYAADLSSSSVEPVE-KSYLGTEGHVNGKIGGNGPPN---PVV-EKKTAKIHDFCLGIPFGGFVLTGGLLGSIFSRNL : 74
Gm ----SVVMSLERHDTDTAGSDTKNTLSYAADVSKLHVEEKK-HSYSTKDGHDTEKLGFGEATQESVHQP-KR-TAKIHDFCLGIPFGGFVLTGGIIGFLFSRSP : 97
Os450 ---------------------------------------CTNGTTQVDELNFRSNQTEELVEGDADTVTQKRSAKIHDFCLGIPFGGLLFCMGLLGYIFSRST : 64
Os690 ------------------------MCMKAEYTSSPVDPNAVAEHTEDENQQPTTVSPTNAEIDIDQEAVPQHKGAIIHDFCLGIPFGGILFSMGLVGFLFWRST : 80
Zm081 PWTTMCVNADYTTPADSVATSEQAGGEAEATLETGGEAAATLEQTGGETEEPTIVAPANEYT--EQEAAPQQKCAKIHDFCLGIPFGGLLLSMGLIGFLFWRSP : 117
Zm990 --------------------------------------CANE-NARIEELDLRSDQMKELQVEEHAIPQ-KRSAKIHDFCLGIPFGGLLFSMGLLGYIFSRST : 63
Pp022 -DTASKELEKLKDITNQLLEQAEIARQKLAATAQETAYKQKDNLAYVAENGPEPVKEVAGSALNAHYSNSGKSGSKIHDFCLGIPFGILLAGGLLWFIISGGT : 103
Pp899 -DIPNKELERLKDITNQLLEQAEAARQKIAATAQETAYKQKDNLAYVAENGPEPVKEVAESALDAHYSKSGKSGSKIHDFCLGIPYGIFLAAGGLLWFILSGGT : 103
Pp988 -ETAYKELEKLRDITKKLLEETDVALNKLAATAQETANKQKDNLADVVEKAPETVKEVAESVLRAHYSEAGGSGSKVHDFCLGIPYGGFLSVGGLLWFILSGSI : 103
Cr --------------------------------------------------------SASASQPSGSAPAGPARHAVLHDFCMVIPFGGAAVLGAVALFFFN-LT : 47
HDFC
IPF/YG
H2
H3
H4
At TSLSTGVLYGGGLLALSTLSLKIWREGKSSFPYILGQAVLSAVVFWKNFTAYSMTK--KLFPAGVFAVISACMLCFYSYVVLSGGNPPPKKLKPS-ATSPSY : 193
Br386 TSLSTGVLYGGGLLALSTLSLKIWRQGKSSFPYILGQAVLSAVVIWKNFTAYSMTK--KLFPAGIFAVVSAAMLCFYSYVVLSGGNPPPKKLKP--STSPSH : 193
Br405 TSLSTGVLYGGGLLALSTLSLKIWRQGKSSFPYILGQAVLSAVVFWKNFTAYSMTK--KLFPAGIFAVVSAAMLCFYSYVVLSGGNPPPKKLKP--STSPSH : 188
Br859 TSLSTGVLYGGGLLALSTLSMKIWRQGKSSFPYILGQAVLSAVVFWKNFTSYSMTK--KLFPAGLFAVVSAAMLCFYSYVVLSGGNPPPKKLKPSATTSPSY : 171
Sl ATLSSGVLFGGALLALSTISMKVWREGKTSFPFILGQAVLAATLLWKNMQTFSLTG--KLFPTGFFAAISAAMFCFYSYVILSGGNPPPKKLKASTSGAY-- : 192
Mt ATLASGVLFGGALLFLSTFSLKVWRQGKSSLPFILGQAALAGILFWKNFQSYSLAK--KIFPTGISAIISSAMLVFYVYVLLSGGNPPPKKLKPTASIE--- : 192
Ps ATLASGVLVGGALLFLSTLSLKVWRQGKSSLPFILGQAALSGILIWKNFQSYSLAK--KIFPTGISAIISSAMLCFYLYVLVSGGNPPPKKLKPSASVA--- : 193
Bv ATLGTGVLFGGAILAFSTISLKVWKEGKSSFPLIVGQAVLAVTLLWKNFQTYSVSK--KLFPSGLYALVSAAMLLFYSYVVISGGNPPPKKMKTAAEAAS-- : 172
Gm ATLGSGVLFGGALLFLSTFSMKVWRQGKSSLPFILGQAALAGVLIWKNFQSYSLAK--KIFPTGISAIISSAMLCFYLYVLISGGNPPPKKLKPSASTA--- : 194
Os450 ISLVLGVAPGLTTLFLGTLSLKFWRSGKSSFIFILGQAAISAVLAWKYSHAYILTN--RILPWAFYASLSTAMACFYAYVLLSGGNPPPKKKMAATPSS--- : 161
Os690 VSLTFGVAPGLAILALGVLSLKVWRSGKSSLPFILAQAAVAAAVAWKHFQAYTTTK--KLLPWAFYTALSAAMICFYSYVLLAGGNPPPKKKAAVAQ----- : 175
Zm081 ASLTFGVAPGLAILALAVLSLNVWRSGKSSLPFILAQAGIAAAVAWKHGQAYTTTR--KLLPWGFYVALSAAMICFYSYVLLAGGNPPPKKAKAKAKAAA-- : 215
Zm990 ISLVLGIAPGLATLLLGTLSLKFWRSGRSSCLLILAQAAISAFLAWKYSHAYFLTN--RILPWGFYASLSTAMGCFYAYVLLAGGNPPPKK-LAAIPPQ--- : 159
Pp022 SAIRFGVILGSALLYLSVTSLKKWKNGESSMPYIQGQAAISAFLFLRYYRRFTVNK--LFFPTGVTGLLSGAMLAFYIYVLVSGGNPPPKKSKIQSS----- : 198
Pp899 SALRFGVLLGSGLLYLSVTSLKKWKNGESSMTYIQGQAAITAFIFLRYYRRYGVNK--VFFPTGVVGLVSGAMLAFYIYVLVSGGNPPPKKSKSQPS----- : 198
Pp988 SAIRFGVLLGSAILYLGLTSLKKWEKGESSMTYIQSQSAITTFIFFRYFRHYTVNK--ALFPTGVVGLISGAMVAFYIYVLASGGNRP-QKPESQTN----- : 197
Cr NIAGTALIAGATAIASSVLSLQEWKAGGSSTTYTLTSAACAAAVSYVTYSSLDLLKGLPYWVAAVLCVIGGACSLFCAYNVAAGGNPPPKKKAAGKAE---- : 145
GGNPPPKK

**B**

| | At | Br389 | Br405 | Br859 | Sl | Mt | Ps | Bv | Gm | Os450 | Os690 | Zm081 | Zm990 | Pp022 | Pp899 | Pp988 | Cr |
|---|---|---|---|---|---|---|---|---|---|---|---|---|---|---|---|---|---|
| At | **100%** | | | | | | | | | | | | | | | | |
| Br389 | **89%**<br>94% | **100%** | | | | | | | | | | | | | | | |
| Br405 | **88%**<br>94% | **94%**<br>98% | **100%** | | | | | | | | | | | | | | |
| Br859 | **77%**<br>93% | **78%**<br>96% | **81%**<br>96% | **100%** | | | | | | | | | | | | | |
| Sl | **54%**<br>67% | **53%**<br>66% | **52%**<br>65% | **50%**<br>67% | **100%** | | | | | | | | | | | | |
| Mt | **53%**<br>63% | **53%**<br>64% | **53%**<br>64% | **50%**<br>64% | **58%**<br>72% | **100%** | | | | | | | | | | | |
| Ps | **51%**<br>63% | **52%**<br>66% | **51%**<br>64% | **48%**<br>64% | **59%**<br>74% | **82%**<br>90% | **100%** | | | | | | | | | | |
| Bv | **48%**<br>66% | **49%**<br>67% | **50%**<br>66% | **52%**<br>67% | **54%**<br>75% | **50%**<br>66% | **52%**<br>66% | **100%** | | | | | | | | | |
| Gm | **47%**<br>61% | **47%**<br>64% | **46%**<br>62% | **44%**<br>64% | **58%**<br>53% | **79%**<br>90% | **81%**<br>94% | **48%**<br>68% | **100%** | | | | | | | | |
| Os450 | **41%**<br>55% | **41%**<br>56% | **42%**<br>56% | **46%**<br>54% | **38%**<br>57% | **37%**<br>51% | **38%**<br>56% | **37%**<br>49% | **36%**<br>54% | **100%** | | | | | | | |
| Os690 | **38%**<br>57% | **38%**<br>58% | **39%**<br>58% | **42%**<br>56% | **38%**<br>59% | **37%**<br>54% | **40%**<br>57% | **39%**<br>54% | **37%**<br>56% | **50%**<br>67% | **100%** | | | | | | |
| Zm081 | **37%**<br>56% | **38%**<br>57% | **39%**<br>57% | **43%**<br>55% | **37%**<br>51% | **35%**<br>54% | **38%**<br>58% | **40%**<br>54% | **37%**<br>57% | **39%**<br>63% | **56%**<br>87% | **100%** | | | | | |
| Zm990 | **37%**<br>53% | **38%**<br>54% | **39%**<br>54% | **42%**<br>52% | **36%**<br>44% | **34%**<br>47% | **36%**<br>52% | **37%**<br>49% | **34%**<br>49% | **72%**<br>86% | **48%**<br>67% | **40%**<br>65% | **100%** | | | | |
| Pp022 | **33%**<br>47% | **32%**<br>46% | **32%**<br>46% | **32%**<br>46% | **34%**<br>42% | **32%**<br>46% | **33%**<br>47% | **32%**<br>47% | **33%**<br>45% | **31%**<br>46% | **31%**<br>44% | **28%**<br>43% | **32%**<br>47% | **100%** | | | |
| Pp899 | **31%**<br>49% | **34%**<br>49% | **34%**<br>49% | **33%**<br>49% | **33%**<br>40% | **32%**<br>45% | **33%**<br>46% | **32%**<br>48% | **33%**<br>45% | **30%**<br>43% | **29%**<br>41% | **27%**<br>40% | **30%**<br>43% | **86%**<br>85% | **100%** | | |
| Pp988 | **31%**<br>45% | **30%**<br>43% | **30%**<br>43% | **29%**<br>43% | **30%**<br>40% | **30%**<br>43% | **31%**<br>45% | **29%**<br>46% | **31%**<br>44% | **28%**<br>41% | **29%**<br>44% | **25%**<br>41% | **28%**<br>44% | **68%**<br>72% | **70%**<br>78% | **100%** | |
| Cr | **24%**<br>31% | **23%**<br>30% | **28%**<br>29% | **27%**<br>30% | **20%**<br>25% | **20%**<br>28% | **20%**<br>28% | **22%**<br>28% | **19%**<br>28% | **26%**<br>28% | **27%**<br>32% | **20%**<br>28% | **26%**<br>27% | **21%**<br>31% | **20%**<br>29% | **16%**<br>24% | **100%** |

Figure S3

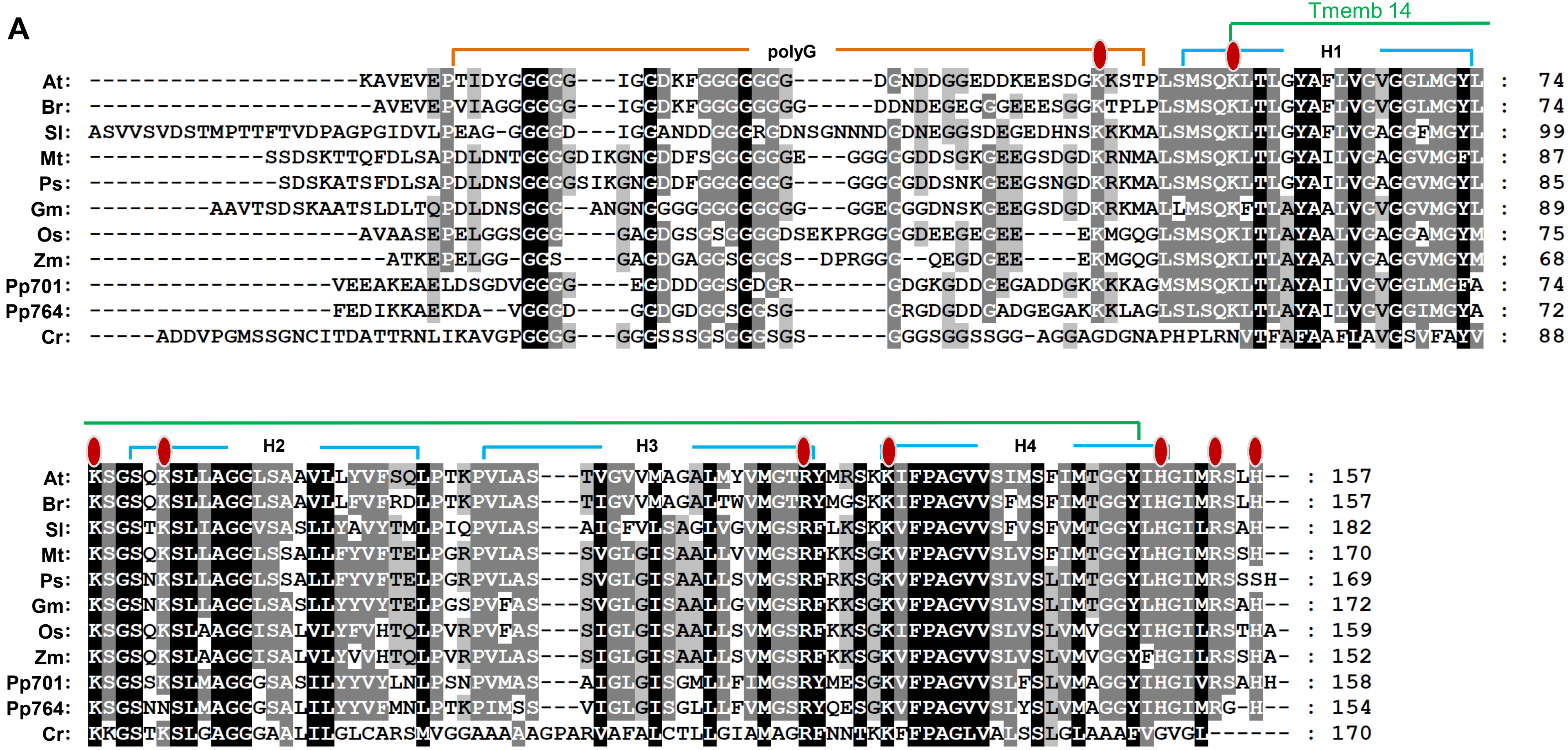

**B**

| | At | Br | Sl | Mt | Ps | Gm | Os | Zm | Pp701 | Pp764 | Cr |
|---|---|---|---|---|---|---|---|---|---|---|---|
| At | **100%** | | | | | | | | | | |
| Br | **84%**<br>91% | **100%** | | | | | | | | | |
| Sl | **51%**<br>67% | **50%**<br>69% | **100%** | | | | | | | | |
| Mt | **57%**<br>71% | **55%**<br>69% | **50%**<br>73% | **100%** | | | | | | | |
| Ps | **56%**<br>70% | **55%**<br>68% | **50%**<br>73% | **90%**<br>94% | **100%** | | | | | | |
| Gm | **51%**<br>67% | **50%**<br>65% | **52%**<br>74% | **74%**<br>85% | **76%**<br>87% | **100%** | | | | | |
| Os | **53%**<br>64% | **54%**<br>65% | **50%**<br>69% | **59%**<br>75% | **60%**<br>76% | **59%**<br>79% | **100%** | | | | |
| Zm | **51%**<br>64% | **52%**<br>64% | **50%**<br>72% | **60%**<br>79% | **61%**<br>80% | **58%**<br>80% | **84%**<br>93% | **100%** | | | |
| Pp701 | **54%**<br>64% | **51%**<br>64% | **47%**<br>66% | **52%**<br>68% | **53%**<br>68% | **57%**<br>72% | **57%**<br>70% | **58%**<br>71% | **100%** | | |
| Pp764 | **53%**<br>63% | **51%**<br>63% | **43%**<br>61% | **51%**<br>65% | **53%**<br>68% | **54%**<br>69% | **54%**<br>69% | **56%**<br>70% | **70%**<br>82% | **100%** | |
| Cr | **24%**<br>37% | **25%**<br>39% | **25%**<br>39% | **25%**<br>40% | **25%**<br>39% | **28%**<br>41% | **28%**<br>40% | **25%**<br>37% | **25%**<br>40% | **27%**<br>41% | **100%** |

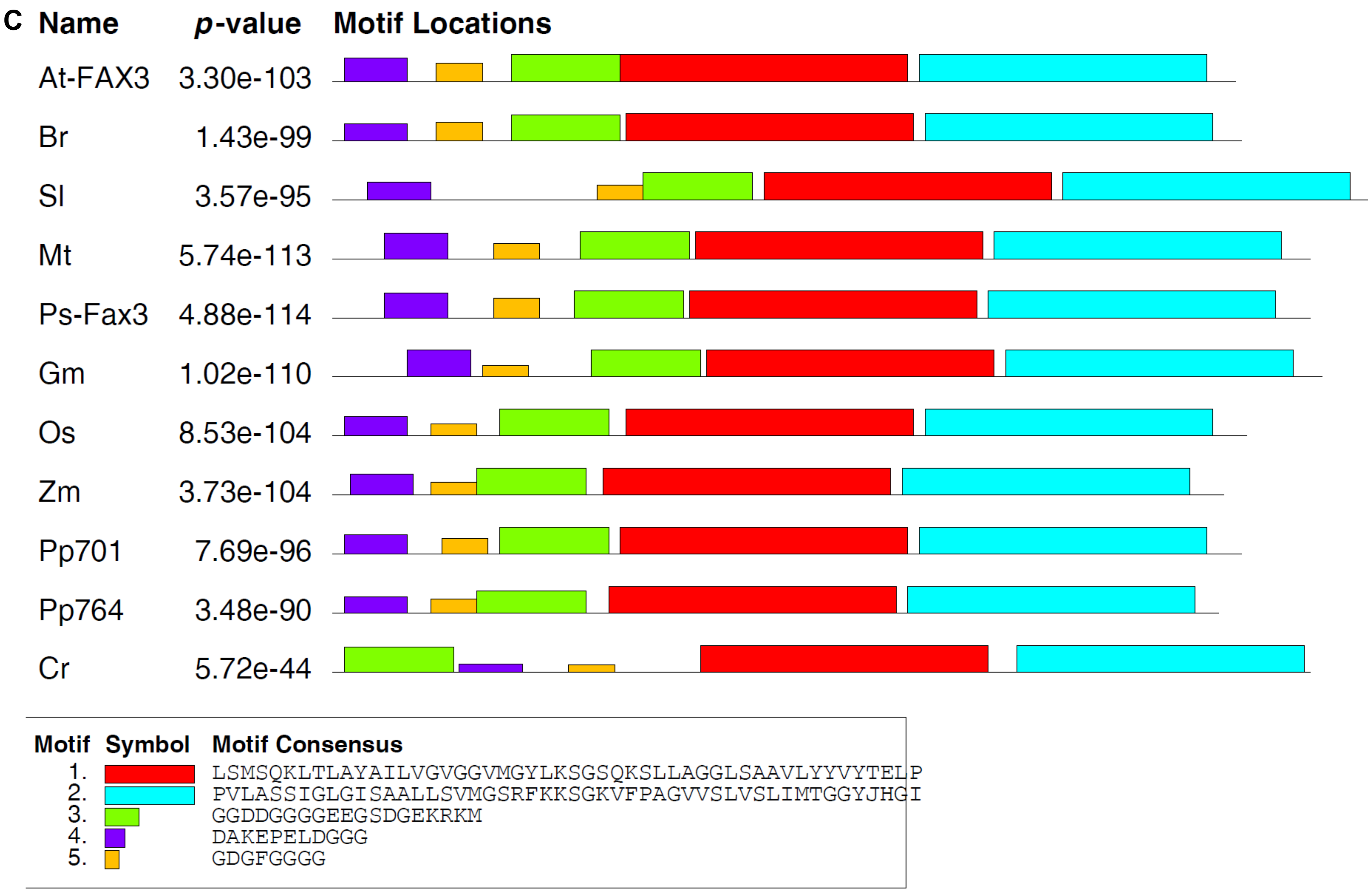
C
Name
p-value
Motif Locations
At-FAX3 3.30e-103
Br 1.43e-99
Sl 3.57e-95
Mt 5.74e-113
Ps-Fax3 4.88e-114
Gm 1.02e-110
Os 8.53e-104
Zm 3.73e-104
Pp701 7.69e-96
Pp764 3.48e-90
Cr 5.72e-44
Motif Symbol Motif Consensus
1. LSMSQKLTLAYAILVGVGGVMGYLKSGSQKSLLAGGLSAAVLYYVYTELP
2. PVLASSIGLGISAALLSVMGSRFKKSGKVFPAGVVSLVSLIMTGGYJHGI
3. GGDDGGGGEEGSDGEKRKM
4. DAKEPELDGGG
5. GDGFGGGG

Figure S4

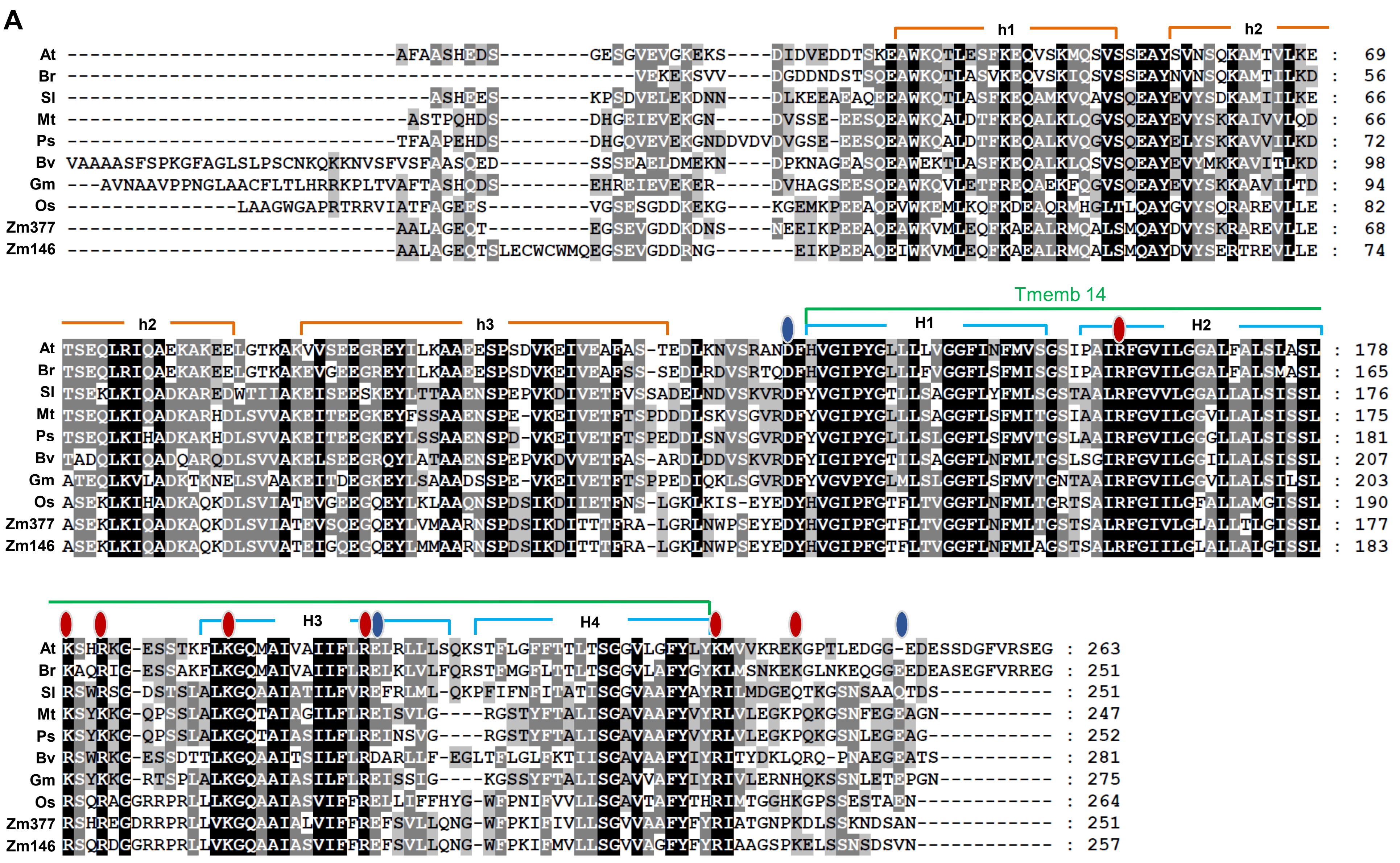

**B**

| | At | Br | Sl | Mt | Ps | Bv | Gm | Os | Zm377 | Zm146 |
|---|---|---|---|---|---|---|---|---|---|---|
| At | **100%** | | | | | | | | | |
| Br | **74%**<br>83% | **100%** | | | | | | | | |
| Sl | **50%**<br>57% | **48%**<br>57% | **100%** | | | | | | | |
| Mt | **48%**<br>60% | **48%**<br>60% | **59%**<br>65% | **100%** | | | | | | |
| Ps | **49%**<br>59% | **47%**<br>59% | **58%**<br>64% | **86%**<br>91% | **100%** | | | | | |
| Bv | **43%**<br>60% | **41%**<br>55% | **53%**<br>63% | **51%**<br>65% | **50%**<br>65% | **100%** | | | | |
| Gm | **42%**<br>55% | **39%**<br>54% | **47%**<br>61% | **65%**<br>81% | **65%**<br>83% | **46%**<br>61% | **100%** | | | |
| Os | **37%**<br>48% | **35%**<br>48% | **41%**<br>50% | **40%**<br>51% | **41%**<br>52% | **38%**<br>53% | **38%**<br>53% | **100%** | | |
| Zm377 | **38%**<br>49% | **35%**<br>46% | **47%**<br>57% | **44%**<br>53% | **43%**<br>50% | **38%**<br>51% | **38%**<br>50% | **67%**<br>76% | **100%** | |
| Zm146 | **36%**<br>50% | **33%**<br>48% | **44%**<br>56% | **43%**<br>53% | **41%**<br>51% | **36%**<br>52% | **37%**<br>51% | **65%**<br>79% | **85%**<br>91% | **100%** |

Figure S5

**A**

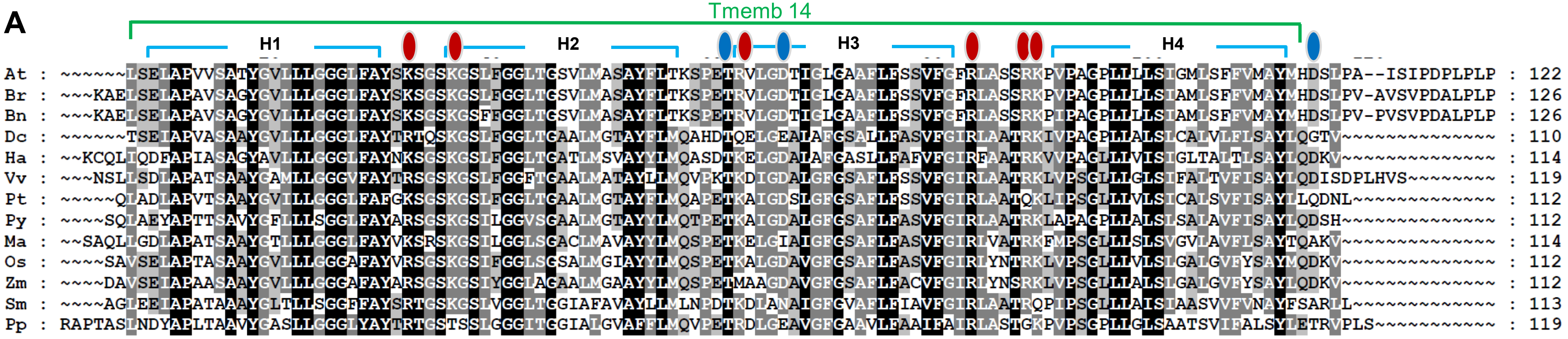


**B**

| | At | Br | Bn | Dc | Ha | Vv | Pt | Py | Ma | Os | Zm | Sm | Pp |
|---|---|---|---|---|---|---|---|---|---|---|---|---|---|
| At | **100%** | | | | | | | | | | | | |
| Br | **92%** | **100%** | | | | | | | | | | | |
| Bn | **89%** | **97%** | **100%** | | | | | | | | | | |
| Dc | **54%** | **51%** | **50%** | **100%** | | | | | | | | | |
| Ha | **49%** | **48%** | **47%** | **66%** | **100%** | | | | | | | | |
| Vv | **54%** | **55%** | **54%** | **63%** | **57%** | **100%** | | | | | | | |
| Pt | **56%** | **54%** | **54%** | **66%** | **59%** | **72%** | **100%** | | | | | | |
| Py | **52%** | **51%** | **51%** | **67%** | **57%** | **68%** | **71%** | **100%** | | | | | |
| Ma | **52%** | **51%** | **51%** | **61%** | **64%** | **64%** | **64%** | **67%** | **100%** | | | | |
| Os | **57%** | **55%** | **54%** | **67%** | **61%** | **67%** | **68%** | **74%** | **71%** | **100%** | | | |
| Zm | **53%** | **52%** | **51%** | **67%** | **58%** | **64%** | **64%** | **72%** | **67%** | **87%** | **100%** | | |
| Sm | **47%** | **49%** | **49%** | **53%** | **47%** | **57%** | **55%** | **53%** | **55%** | **53%** | **53%** | **100%** | |
| Pp | **42%** | **43%** | **42%** | **50%** | **42%** | **50%** | **47%** | **47%** | **46%** | **46%** | **45%** | **52%** | **100%** |

Figure S7

**A**

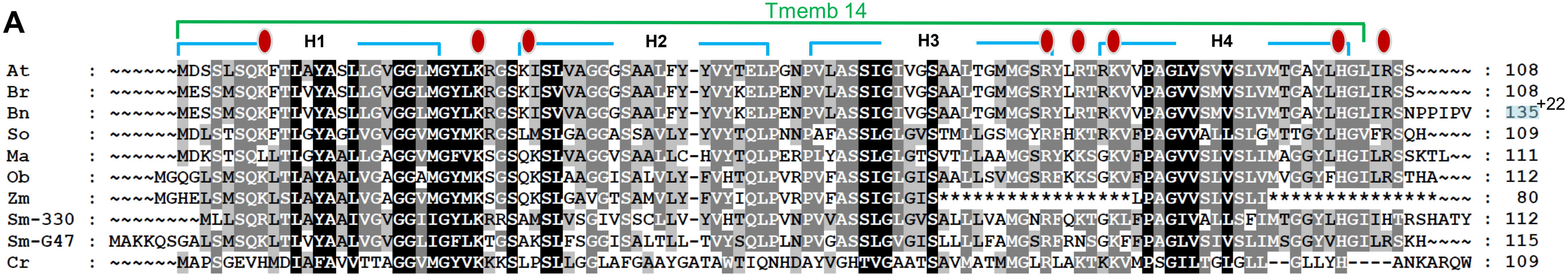


**B**

| | At | Br | Bn | So | Ma | Ob | Zm | Sm-330 | Sm-G47 | Cr |
|---|---|---|---|---|---|---|---|---|---|---|
| At | **100%** | | | | | | | | | |
| Br | **92%** | **100%** | | | | | | | | |
| Bn | **73%** | **79%** | **100%** | | | | | | | |
| So | **61%** | **59%** | **48%** | **100%** | | | | | | |
| Ma | **58%** | **57%** | **46%** | **62%** | **100%** | | | | | |
| Ob | **57%** | **56%** | **45%** | **62%** | **68%** | **100%** | | | | |
| Zm | **40%** | **41%** | **33%** | **46%** | **47%** | **60%** | **100%** | | | |
| Sm-330 | **50%** | **46%** | **39%** | **57%** | **53%** | **53%** | **52%** | **100%** | | |
| Sm-G47 | **51%** | **51%** | **41%** | **53%** | **58%** | **63%** | **55%** | **52%** | **100%** | |
| Cr | **26%** | **26%** | **21%** | **29%** | **28%** | **26%** | **30%** | **24%** | **20%** | **100%** |

Figure S8

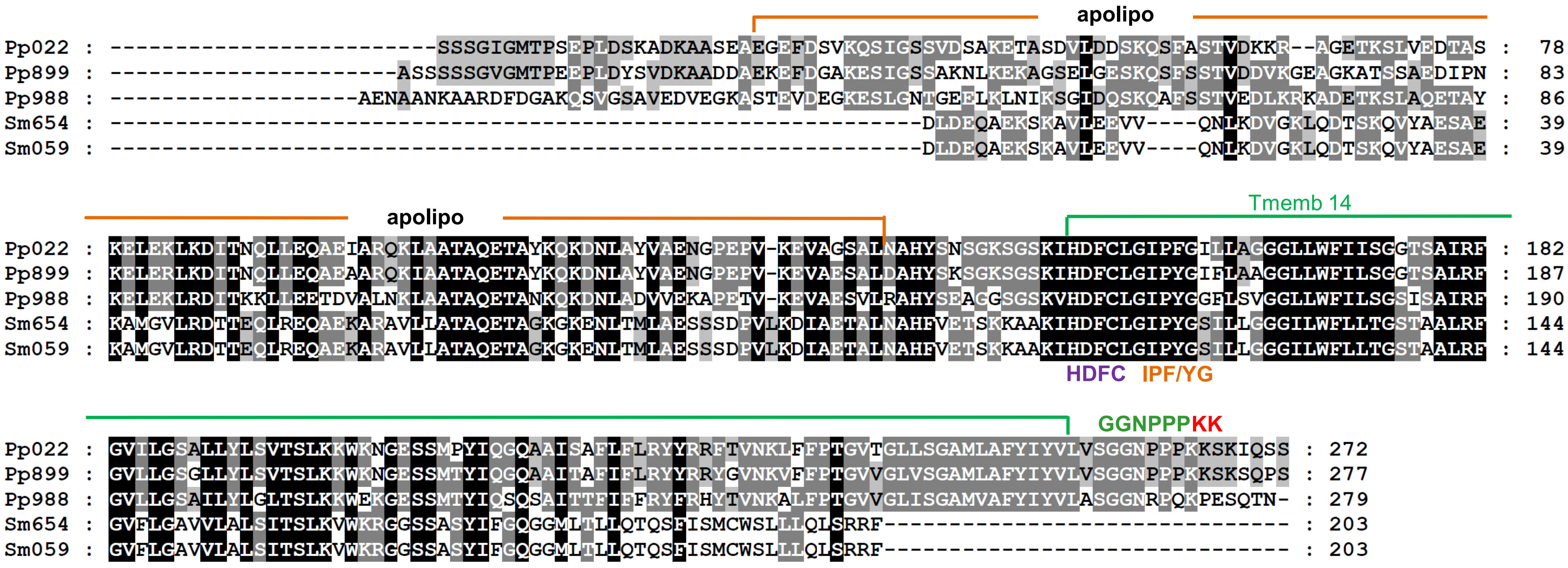
apolipo
Pp022 : ---------------------------SSSGIGMTPSEPLDSKADKAASEAEGEFDSVKQSIGSSVDSAKETASDVLDDSKQSFASTVDKKR--AGETKSLVEDTAS : 78
Pp899 : -----------------------ASSSSSGVGMTPEEPLDYSVDKAADDAEKEFDGAKESIGSSAKNLKEKAGSELGESKQSFSSTVDDVKGEAGKATSSAEDIPN : 83
Pp988 : --------------------AENAANKAARDFDGAKQSVGSAVEDVEGKASTEVDEGKESLGNTGEELKLNIKSGIDQSKQAFSSTVEDLKRKADETKSLAQETAY : 86
Sm654 : -----------------------------------------------------------------DLDEQAEKSKAVLEEVV----QNLKDVGKLQDTSKQVYAESAE : 39
Sm059 : -----------------------------------------------------------------DLDEQAEKSKAVLEEVV----QNLKDVGKLQDTSKQVYAESAE : 39
apolipo
Tmemb 14
Pp022 : KELEKLKDITNQLLEQAEIARQKLAATAQETAYKQKDNLAYVAENGPEPV-KEVAGSALNAHYSNSGKSGSKIHDFCLGIPFGILLAGGGLLWFIISGGTSAIRF : 182
Pp899 : KELERLKDITNQLLEQAEAARQKIAATAQETAYKQKDNLAYVAENGPEPV-KEVAESALDAHYSKSGKSGSKIHDFCLGIPYGIFLAAGGLLWFILSGGTSALRF : 187
Pp988 : KELEKLRDITKKLLEETDVALNKLAATAQETANKQKDNLADVVEKAPETV-KEVAESVLRAHYSEAGGSGSKVHDFCLGIPYGGFLSVGGLLWFILSGSISAIRF : 190
Sm654 : KAMGVLRDTTEQLREQAEKARAVLLATAQETAGKGKENLTMLAESSSDPVLKDIAETALNAHFVETSKKAAKIHDFCLGIPYGSILLGGGILWFLLTGSTAALRF : 144
Sm059 : KAMGVLRDTTEQLREQAEKARAVLLATAQETAGKGKENLTMLAESSSDPVLKDIAETALNAHFVETSKKAAKIHDFCLGIPYGSILLGGGILWFLLTGSTAALRF : 144
HDFC
IPF/YG
GGNPPPKK
Pp022 : GVILGSALLYLSVTSLKKWKNGESSMPYIQGQAAISAFLFLRYYRRFTVNKLFFPTGVTGLLSGAMLAFYIYVLVSGGNPPPKKSKIQSS : 272
Pp899 : GVLLGSGLLYLSVTSLKKWKNGESSMTYIQGQAAITAFIFLRYYRRYGVNKVFFPTGVVGLVSGAMLAFYIYVLVSGGNPPPKKSKSQPS : 277
Pp988 : GVLLGSAILYLGLTSLKKWEKGESSMTYIQSQSAITTFIFFRYFRHYTVNKALFPTGVVGLISGAMVAFYIYVLASGGNRPQKPESQTN- : 279
Sm654 : GVFLGAVVLALSITSLKVWKRGGSSASYIFGQGGMLTLLQTQSFISMCWSLLLQLSRRF------------------------------- : 203
Sm059 : GVFLGAVVLALSITSLKVWKRGGSSASYIFGQGGMLTLLQTQSFISMCWSLLLQLSRRF------------------------------- : 203

Figure S9

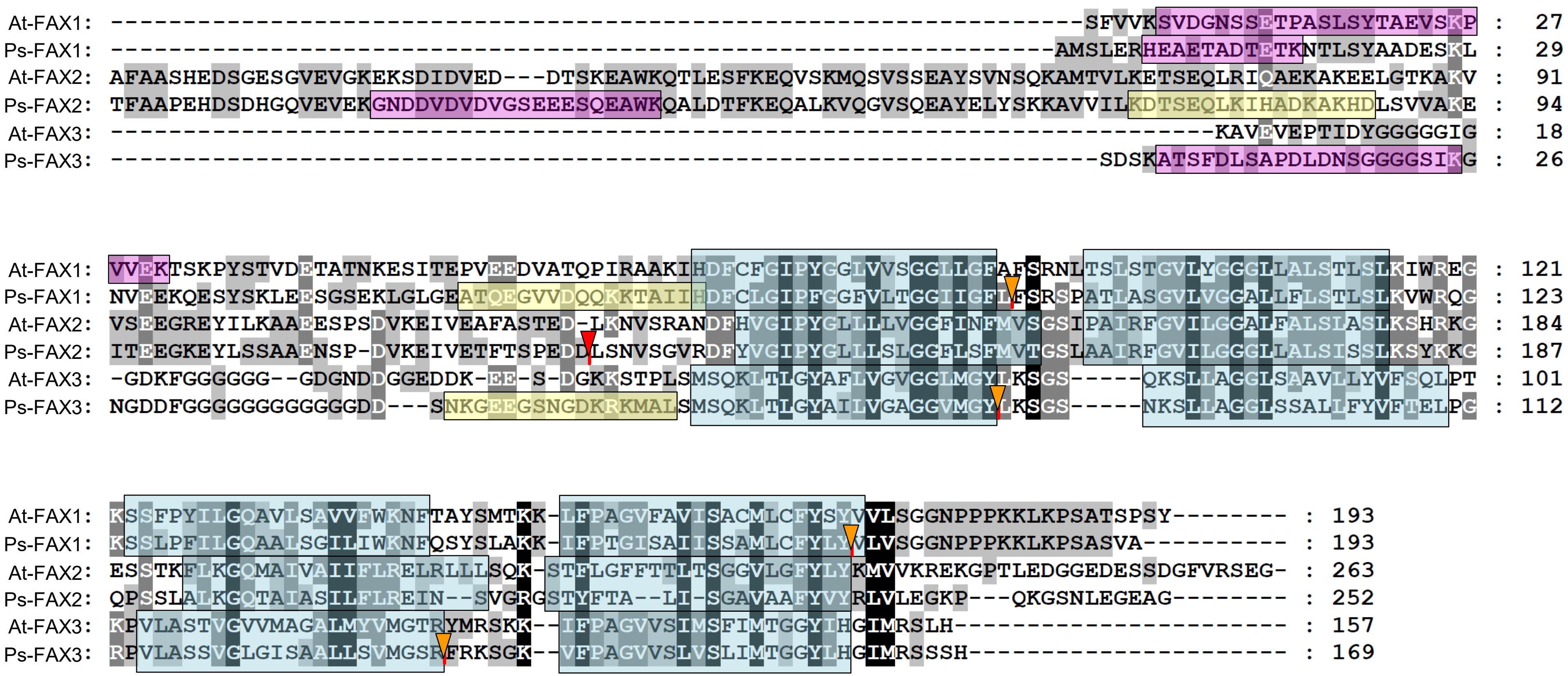
At-FAX1: -------------------------------------------------------------------SFVVKSVDGNSSETPASLSYTAEVSKP : 27
Ps-FAX1: -----------------------------------------------------------------AMSLERHEAETADTETKNTLSYAADESKL : 29
At-FAX2: AFAASHEDSGESGVEVGKEKSDIDVED---DTSKEAWKQTLESFKEQVSKMQSVSSEAYSVNSQKAMTVLKETSEQLRIQAEKAKEELGTKAKV : 91
Ps-FAX2: TFAAPEHDSDHGQVEVEKGNDDVDVDVGSEEESQEAWKQALDTFKEQALKVQGVSQEAYELYSKKAVVILKDTSEQLKIHADKAKHDLSVVAKE : 94
At-FAX3: -----------------------------------------------------------------------------KAVEVEPTIDYGGGGIG : 18
Ps-FAX3: --------------------------------------------------------------------SDSKATSFDLSAPDLDNSGGGGSIKG : 26
At-FAX1: VVEKTSKPYSTVDETATNKESITEPVEEDVATQPIRAAKIHDFCFGIPYGGLVVSGGLLGFAFSRNLTSLSTGVLYGGGLLALSTLSLKIWREG : 121
Ps-FAX1: NVEEKQESYSKLEESGSEKLGLGEATQEGVVDQQKKTAIIHDFCLGIPFGGFVLTGGIIGFLFSRSPATLASGVLVGGALLFLSTLSLKVWRQG : 123
At-FAX2: VSEEGREYILKAAEESPSDVKEIVEAFASTED-LKNVSRANDFHVGIPYGLLLLVGGFINFMVSGSIPAIRFGVILGGALFALSIASLKSHRKG : 184
Ps-FAX2: ITEEGKEYLSSAAENSP-DVKEIVETFTSPEDDLSNVSGVRDFYVGIPYGLLLSLGGFLSFMVTGSLAAIRFGVILGGGLLALSISSLKSYKKG : 187
At-FAX3: -GDKFGGGGGG--GDGNDDGGEDDK-EE-S-DGKKSTPLSMSQKLTLGYAFLVGVGGLMGYLKSGS-----QKSLLAGGLSAAVLLYVFSQLPT : 101
Ps-FAX3: NGDDFGGGGGGGGGGGDD---SNKGEEGSNGDKRKMALSMSQKLTLGYAILVGAGGVMGYLKSGS-----NKSLLAGGLSSALLFYVFTELPG : 112
At-FAX1: KSSFPYILGQAVLSAVVFWKNFTAYSMTKK-LFPAGVFAVISACMLCFYSYVVLSGGNPPPKKLKPSATSPSY-------- : 193
Ps-FAX1: KSSLPFILGQAALSGILIWKNFQSYSLAKK-IFPTGISAIISSAMLCFYLYVLVSGGNPPPKKLKPSASVA---------- : 193
At-FAX2: ESSTKFLKGQMAIVAIIFLRELRLLLSQK-STFLGFFTTLTSGGVLGFYLYKMVVKREKGPTLEDGGEDESSDGFVRSEG- : 263
Ps-FAX2: QPSSLALKGQTAIASILFLREIN--SVGRGSTYFTA--LI-SGAVAAFYVYRLVLEGKP---QKGSNLEGEAG-------- : 252
At-FAX3: KPVLASTVGVVMAGALMYVMGTRYMRSKK--IFPAGVVSIMSFIMTGGYIHGIMRSLH---------------------- : 157
Ps-FAX3: RPVLASSVGLGISAALLSVMGSRFRKSGK--VFPAGVVSLVSLIMTGGYLHGIMRSSH---------------------- : 169
