## Supplementary figures and images for "Plastid fatty acid export (FAX) proteins in *Arabidopsis thaliana* - the role of FAX1 and FAX3 in growth and development"

### Supplemental Fig S6, S10,11

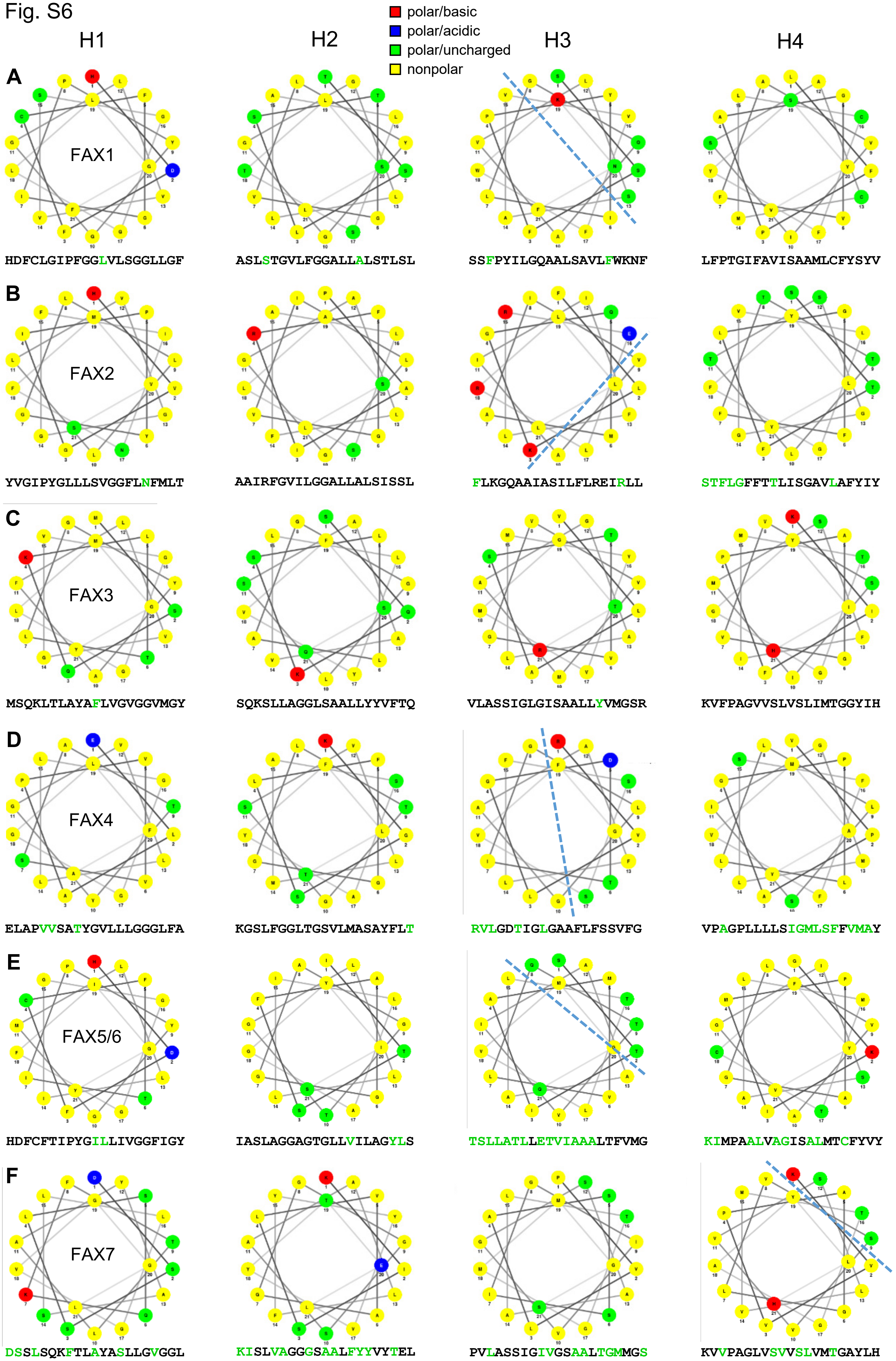

Fig. S10

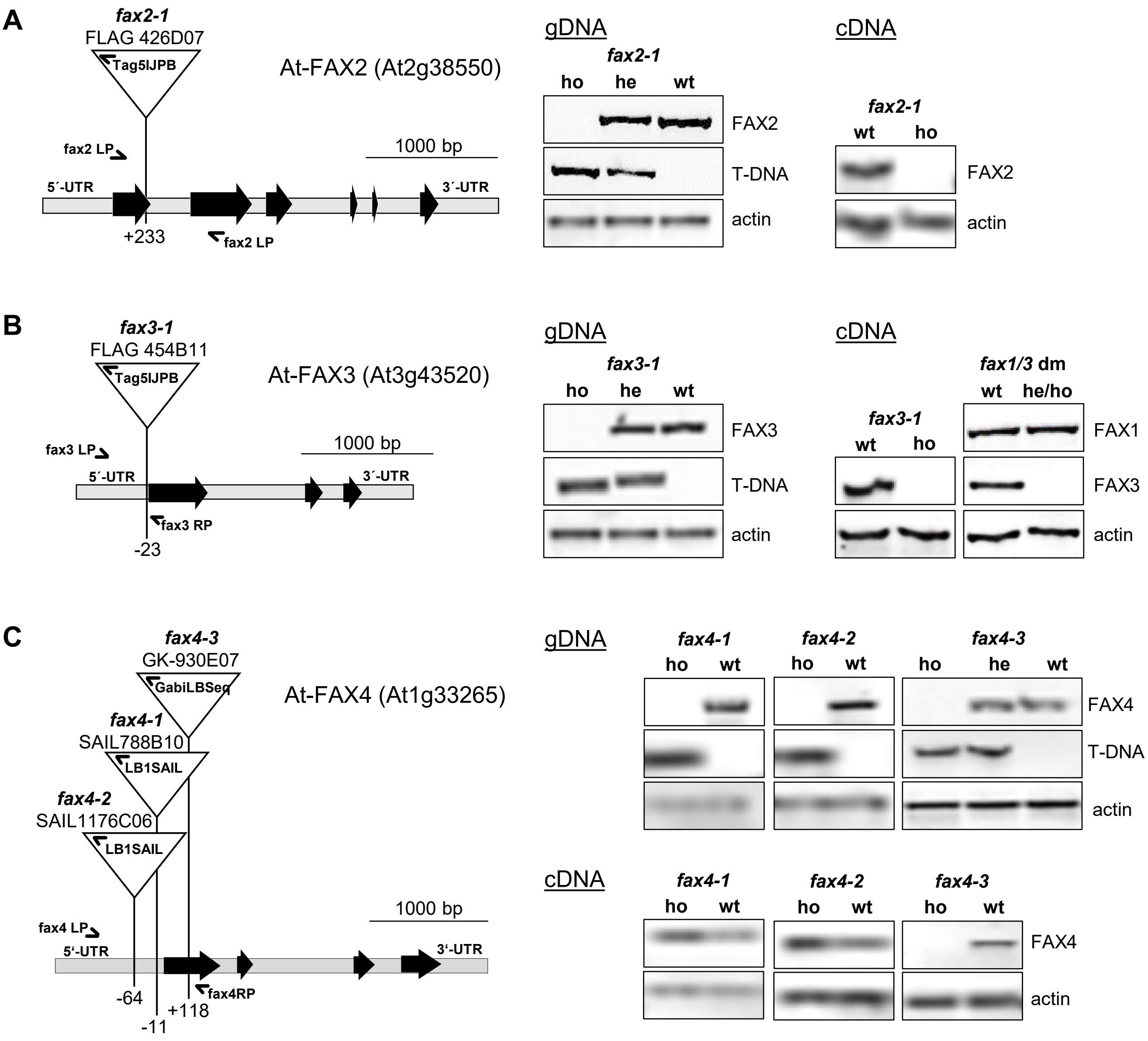


**D**

| | no. | ho [%] | he [%] | wt [%] |
|---|---|---|---|---|
| **agar** | | | | |
| *fax2-1* | 146 | 28.8 | 44.5 | 26.7 |
| *fax3-1* | 122 | 23.0 | 36.9 | 40.2 |
| *fax4-3* | 74 | 24.3 | 44.6 | 31.1 |

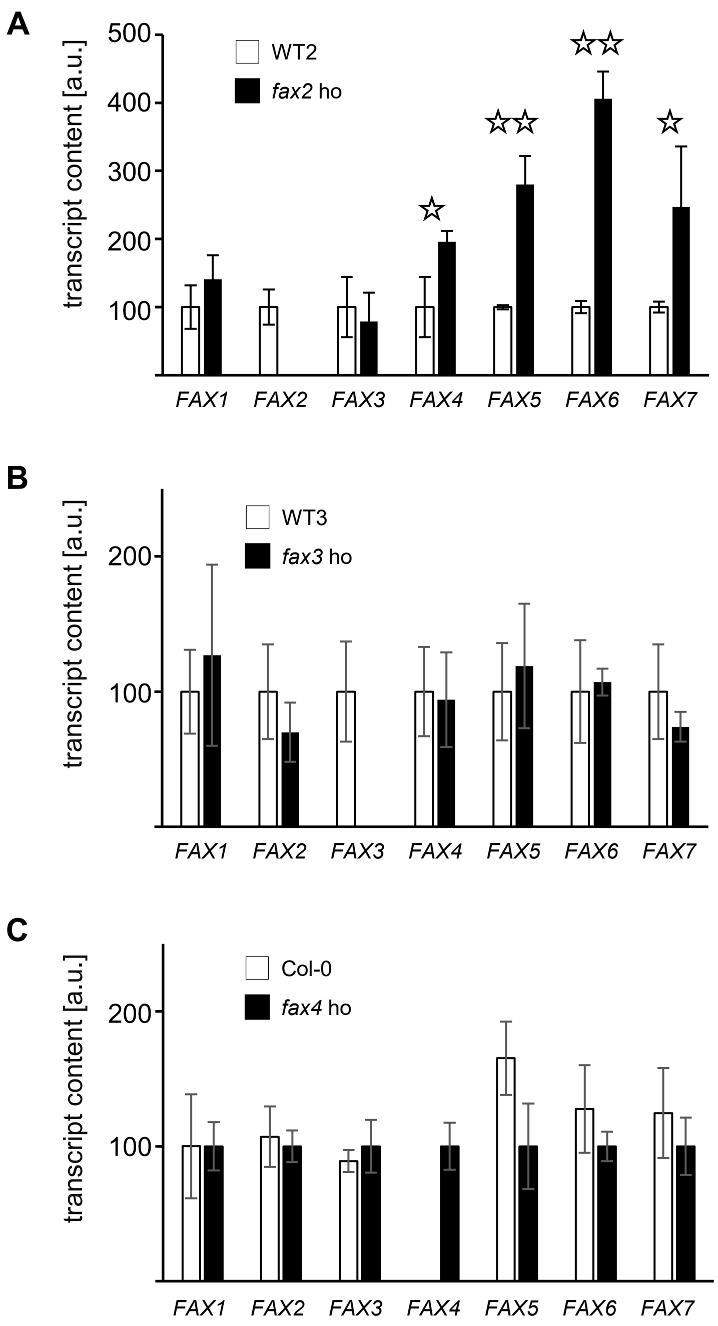
